# Identification of hypoxia-specific biomarkers in salmonids using RNA-sequencing and validation using high-throughput qPCR

**DOI:** 10.1101/2020.05.14.086090

**Authors:** Arash Akbarzadeh, Aimee Lee S. Houde, Ben J.G. Sutherland, Oliver P. Günther, Kristina M. Miller

## Abstract

Identifying early gene expression responses to hypoxia (i.e., low dissolved oxygen) as a tool to assess the degree of exposure to this stressor is crucial for salmonids, because they are increasingly exposed to hypoxic stress due to anthropogenic habitat change, e.g., global warming, excessive nutrient loading, and persistent algal blooms. Our goal was to discover and validate gill gene expression biomarkers specific to the hypoxia response in salmonids across multi-stressor conditions. Gill tissue was collected from 24 freshwater juvenile Chinook salmon (*Oncorhynchus tshawytscha*), held in normoxia [dissolved oxygen (DO) > 8 mg L^−1^] and hypoxia (DO = 4□5 mg L^−1^) in 10 and 18°C temperatures for up to six days. RNA-sequencing (RNA-seq) was then used to discover 240 differentially expressed genes between hypoxic and normoxic conditions, but not affected by temperature. The most significantly differentially expressed genes had functional roles in the cell cycle and suppression of cell proliferation associated with hypoxic conditions. The most significant genes (n = 30) were selected for real-time qPCR assay development. These assays demonstrated a strong correlation (r = 0.88; p < 0.001) between the expression values from RNA-seq and the fold changes from qPCR. Further, qPCR of the 30 candidate hypoxia biomarkers was applied to an additional 322 Chinook salmon exposed to hypoxic and normoxic conditions to reveal the top biomarkers to define hypoxic stress. Multivariate analyses revealed that smolt stage, water salinity, and morbidity status were relevant factors to consider with the expression of these genes in relation to hypoxic stress. These hypoxia candidate genes will be put into application screening Chinook salmon to determine the identity of stressors impacting the fish.

## Introduction

Aquatic ecosystems are increasingly being affected by a complex mixture of anthropogenic stressors (Hering *et al*. 2015). In recent years, the prevalence and intensity of aquatic hypoxic episodes has increased worldwide, resulting in aquatic environments with large variations in dissolved oxygen (Leveelahti *et al*. 2011; Robertson *et al*. 2014). Both natural and anthropogenic drivers can be responsible for dissolved oxygen (DO) depletion in aquatic ecosystems. Natural hydrodynamic conditions, such as the oxidation of organic matter and the release of gases (e.g., methane or carbon dioxide), may make certain aquatic ecosystems prone to oxygen depletion. However, climate change can also reduce the solubility of oxygen because of increased thermal stratification and enhanced microbial activity, all of which would cause DO depletion (Weinke and Biddanda 2018). Moreover, agriculture, industry, and urbanization can lead to nutrient loading (i.e., eutrophication) and consequent persistent algal blooms in aquatic ecosystems, which also affect both the supply and uptake of DO (Friedrich *et al*. 2014). The combined effects of warming and eutrophication in aquatic ecosystems may cause dramatic declines in DO, which can lead to hypoxic conditions.

Hypoxia is defined as any level of DO that is low enough to negatively impact the behaviour and/or physiology of an organism. However, the impacts of different hypoxia levels and their effects on fish physiology or behavior is very species-specific (Pollock *et al*. 2007). For example, cyprinid fish, such as the goldfish (*Carassius auratus*) and the crucian carp (*C. carassius*), exhibit a striking capacity to survive and remain active for long periods under very low DO, even tolerating anoxia (Gattuso *et al*. 2018). In contrast, DO below 5 mg/L is the threshold for hypoxic stress in salmon (Lucas and Southgate. 2013; Olsvic *et al*. 2013; Del Rio *et al*. 2019).

Hypoxia can adversely impact a wide variety of biochemical and physiological processes in fish, such as cell cycle, fish growth, and aerobic metabolism (Zhang *et al*. 2009; Douxfils *et al*. 2012). Hypoxia can also negatively impact normal biological functions, resulting in suppressed development, reduced activity, feeding and growth, disturbed endocrine function, and impaired reproductive performance (Callier *et al*. 2013; Wang *et al*. 2016; Abdel-Tawwab *et al*. 2019). Under extreme hypoxic conditions, mass mortality of wild populations can occur (Douxfils et al. 2012; Long et al. 2015; Closs *et al*. 2016). Owing to the significance of hypoxic stress in fish, the study of the effects of hypoxia on fishes has flourished, and received much attention by fish physiologists (Pollock *et al*. 2007; Richards *et al*. 2009).

Salmon can be negatively impacted by hypoxia through impacts on swim performance, growth, and development, which may ultimately affect fitness and survival. Typically, fish respond to hypoxia through a number of behavioural, morphological, physiological, and molecular changes. Behaviourally, fish can actively avoid hypoxic areas or reduce energy demands by decreasing swimming activity. Morphologically, fish can adapt by increasing the respiratory surface area on the gills. Physiological and molecular adaptations include enhanced production of respiratory proteins (e.g. hemoglobin, myoglobin, and neuroglobin) to increase oxygen carrying capacity, upregulation of genes encoding enzymes within the glycolytic pathway, and downregulation of genes involved in aerobic energy production and other energy-consuming processes (Tiedke *et al*. 2015). Differential regulation of genes involved in general metabolism, catabolism, and the ubiquitin-proteasome pathway have also been demonstrated in fish as responses to low DO (Ju *et al*. 2007; Zhang *et al*. 2012; Olsvik *et al*. 2013).

Every year, billions of juvenile salmon pass through estuaries on the North American Pacific coast, where anthropogenic impacts and climate change may increase the chance of encountering hypoxic conditions (Birtwell and Kruzynski 1989). Given the intensive migrations these fish must endure to reach optimal feeding grounds in the ocean and their need to evade predators along the way (Quinn, 2005), if exposure to low DO results in reduced swimming performance, as demonstrated by Bjornn and Reiser (1991), that could have dire, indirect consequences on survival. Eutrophic lakes can also become hypoxic (Conley *et al*. 2009), which may be particularly problematic for fry and adult salmon using these habitats, e.g., Sockeye salmon *Oncorhynchus nerka* in Cultus Lake, BC (Putt *et al*. 2019), and Okanagan system lakes (Hyatt et al. 2003). Low DO can also have adverse effects in combination with other stressors such as thermal stress (Del Rio et al. 2019), and under extreme conditions low DO can be lethal to salmonids (Carter 2005).

Measuring DO concentrations in aquatic ecosystems is one aspect of monitoring hypoxic stress. However, frequent DO measurements over large regions and long periods of time are impractical, unless sampling moorings are continuously used. Furthermore, such environmental measurements do not directly assess whether hypoxic stress was experienced by a fish. We know that low DO conditions have serious consequences for farmed salmon (e.g. Burt *et al*. 2012; Solstorm *et al*. 2018), but it is not clear whether migratory fish inhabiting or moving through such conditions can behaviourally moderate their exposure or whether they are also negatively affected. The application of biomarkers to detect the specific stress response to low DO, and to other stressors for that matter, could thus be an important approach (Zhang *et al*. 2012; Akbarzadeh *et al*. 2018), and be a more integrative technique (Froehlich *et al*. 2015; Houde *et al*. 2019a). Biomarkers can be defined as measurable biochemical, cellular, and/or physiological changes in an organism caused by perturbations in the environmental conditions, such as hypoxia. The ideal biomarker should be specific to the stressor of interest, be easy to assay, and be relatively unaffected by sampling procedures (Zhang *et al*. 2012).

A major advantage of the biomarker approach is the ability to detect sub-lethal impacts at low levels of stressor intensity. Moreover, the inferred consequences of sub-lethal exposure can be made more robust because the biomarkers can also provide detailed physiological information of an organism (Froehlich *et al*. 2015). Molecular ecologists have become interested in studying early genetic responses of various organisms to stressors such as hypoxia (Nikinmaa and Rees 2005; Boswell *et al*. 2009). Several approaches have been used to identify hypoxia responsive genes in fishes and their altered expression levels, including microarrays, real-time qPCR, and proteomics (Rashid *et al*. 2017). RNA-seq has been more recently used to identify the transcriptional responses of fish to hypoxia at the whole transcriptome level without a reference genome (Olsvic *et al*. 2013; Long *et al*. 2015; Zhang *et al*. 2016).

As most genes and proteins can be involved in several physiological pathways, it may not be realistic to obtain such a specific predictor with a single biomarker. Miller *et al*. (2017) demonstrated that the detection of a viral disease state in salmon could be accomplished by the co-regulation of as few as seven biomarkers. This approach has led to the discovery of several novel viruses in salmon (Mordecai *et al*. 2020) and the elucidation of the developmental pathway of a natural disease outbreak, from a viral carrier state to disease causing death (Di Cicco *et al*. 2018). Recently, salmonid biomarker panels have been developed and validated for the detection of thermal and salinity stress (Akbarzadeh *et al*. 2018; Houde *et al*. 2019a), but the identification of a panel specific to hypoxic stress activated in gill tissue has proven to be more difficult. This may be because of a lack of available transcriptional studies to use to identify candidate hypoxia biomarkers in salmon gill, whereas such studies were available for thermal and salinity stress. For hypoxia, only two genes (i.e., *hypoxia inducible factor 1 alpha* [*hif1a*] and *cytochrome C oxidase subunit 6B1* [*cox6b1*]) of the 24 candidate hypoxia genes applied to the gill tissue of juvenile ocean-type Chinook salmon (*O. tshawytscha*) exposed to combinations of salinity, temperature, and DO manipulations were primarily influenced by DO. However, these two genes alone were not sufficient to separate salmon exposed to normoxia from those exposed to hypoxia across differing salinities and temperatures.

Our objective here was to discover and validate a biomarker panel of at least eight genes specific to the hypoxia response across several conditions. To this end, we applied RNA-seq (Wang *et al*. 2009; Ozsolak and Milos 2011) with best practices for analysis (Conesa et al. 2016) to discover additional candidate genes for the response to hypoxia in salmonids. It is well known that the gill is a multifunctional organ (Evans et al. 2005). Oxygenation in the gill is only dependent on the ambient oxygen level from direct contact with the external environment, whereas oxygenation in all other tissues also depends on circulatory oxygen perfusion. Therefore, temperature and hypoxia effects can only be separated using gill tissue. In this regard, gill tissue is ideal for identifying specific hypoxia biomarkers in salmon. In total, gill tissue from 24 freshwater (FW) Chinook salmon juveniles were characterized by RNA-sequencing after exposure to normoxia or hypoxia within 10 and 18°C temperatures (n = 6 fish per condition). These fish were sourced from a previous exposure experiment (Houde *et al*. 2019a). TaqMan assays were then designed to the top 30 candidate hypoxia genes (based on p-values) that were consistent across temperatures. Biomarkers were validated using 301 Chinook salmon from the challenge study that were not used for sequencing. Recent studies have shown that intermittent diel cycles of hypoxia (12 h normoxia: 12 h hypoxia) might give different results as compared to constant (sustained) hypoxia (Borowiec et al. 2015). Therefore, in this study we exposed fish to one hypoxia level and one hypoxia duration.

Our ultimate aim is to add a validated hypoxia-specific biomarker panel to similar panels for thermal, salinity, and general stress, as well as panels indicating different disease states (viral disease, inflammation/wounding response, humoral/cellular response, and imminent mortality) applied simultaneously within a newly developed microfluidics-based salmon “Fit-Chip” tool (Miller et al. 2018). The Fit-Chips will then be applied to examine the synergistic interplay between stress and disease, to elucidate the combinations of stressors most impactful to salmon survival, and to identify habitats where fish are the most compromised. This information can then be used to develop plans for environmental mitigation.

## Materials and Methods

### Experimental set-up

The experiment was approved by the Fisheries and Oceans Canada (DFO) Pacific Region Animal Care Committee (2017-002) following the Canadian Council of Animal Care Standards. Sub-yearling Chinook salmon were provided by Big Qualicum Hatchery, Qualicum, British Columbia (BC), Canada, and were reared in communal tanks at the Pacific Biological Station, Nanaimo, BC. Juveniles were exposed to FW (< 14°C), light at the natural cycle, and fed a ration of commercial pellets (2% body mass, Bio-Oregon) every 1–2 days until used in the experiment. Four trials were conducted over the smoltification period, covering pre-smolt (March), smolt (May), and de-smolt (two trials, June and August) stages based on expected differences in seawater (SW) mortality, gill Na^+^/K^+^-ATPase activity, and body variables (see Houde *et al*. 2019a).

During the experiment, as detailed in Houde et al. (2019a) and in brief here, juveniles were exposed to 18 possible water treatments: three salinities (FW at 0 PSU, brackish (BW) at 20 PSU, and SW at 28 or 29 PSU), three temperatures (10, 14, and 18°C), and two dissolved oxygen (DO) concentrations (hypoxia 4□5 mg L^−1^ and normoxia > 8 mg L^−1^) in all combinations. Each treatment was represented by two (replicate) 30 L pot tanks with tight fitting lids that limited gas exchange. Water at the desired salinity and temperature was divided into two PVC columns: one with media for typical aeration for normoxia and the other with a ceramic air stone delivering very small nitrogen bubbles (5–10 μm) for hypoxia. Nitrogen was supplied to the system using portable liquid units or compressed gas bottles (Praxair). Hypoxia conditions were measured to ensure target levels using nine DO probes, where a single probe was present in one of the two replicate tanks. Each probe was connected to a Point Four RIU3 monitor-controller (Pentair) for turning the nitrogen regulator on or off as required to keep DO in range. Collectively, the controllers were connected to a Point Four LC3 central water system (Pentair).

### Juvenile handling

Prior to treatment exposure, 12□16 juveniles were acclimated to each tank (n = 18 tanks) for six days under the control conditions (i.e., FW, 13□14°C, normoxia), and fed a ration every day. Water conditions were then changed over 1□2 days. On day 1, higher salinity water was introduced over 2□3 h in morning, then temperature was changed by 2°C in the early afternoon, and dissolved oxygen was set to 6.5□8 mg L^−1^ in the mid-afternoon using the LC3 system. On the morning of day 2, temperature was changed by the remaining 2°C and DO was set to 4□5 mg L^−1^. Juveniles were then exposed to full treatment conditions for six days and were fed rations with 48 h starvation before final sampling. On day six, juveniles were euthanized in an overdose of TMS (250 mg L^−1^, buffered for FW treatments) using normoxic water of the same salinity and temperature. Gill tissue from the left side was dissected, placed in RNAlater (Invitrogen) for 24 h in a 4°C fridge, and then stored in a −80°C freezer until used for RNA extraction.

As previously described by Houde et al. (2019a), the two SW/18°C-hypoxia tanks in the June, de-smolt trial (i.e., trial 3) had lower DO than intended in the SW treatment due to a programming failure (3.3–4.1 mg L^−1^ instead of 4–5 mg L^−1^). After almost two days of exposure, 11 of 24 juveniles were dead or moribund, and the remaining 13 juveniles were ethically euthanized. Because of a poor separation between normoxia and hypoxia using the general hypoxia candidate genes previously applied (see Houde *et al*. 2019a), these fish were used as an ‘extreme hypoxia’ condition. Greater details on juvenile handling and sampling are described in Houde *et al*. (2019a)

### RNA extraction and sample selection

Gill tissues from all of the samples described above were homogenized in TRIzol (Ambion) and 1–bromo–3–chloropropane (BCP) reagent with stainless steel beads using a mixer mill (Retch Inc., MM301) in 30 Hz for 3 min. The aqueous supernatant of the homogenate was removed and then used for extraction of RNA using the ‘No-Spin Procedure’ of MagMAX-96 Total RNA Isolation kits (Ambion) as per manufacturer’s instructions and a Biomek FXP automation workstation (Beckman-Coulter). The resulting total RNA was stored at −80°C.

A subset of 24 individuals was selected for RNA-seq from the two de-smolt trials (trials 3 and 4). The FW individuals were selected as there is an immediate need for application of the developed panel in freshwater. The more extreme temperatures were selected to ensure robustness of markers across different temperatures. Two individuals from each tank replicate were selected from Trial 3, and one individual from each tank replicate from Trial 4. Conditions included were two temperatures (10°C and 18°C) and two DO conditions (normoxia and hypoxia).

### Library preparation and sequencing

RNA-seq library preparation and sequencing was performed by D-Mark at the University of California, Davis, California, USA. Total RNA integrity of extracted gill tissue samples was measured using a BioAnalyzer 2100 (Agilent). Samples with RNA integrity numbers (RIN) ≥ 6 were selected for libraries and sequencing; one hypoxia-18°C individual from the June trial was substituted with a similar individual from the August trial to meet this standard.

Libraries were constructed using NEBNext Ultra RNA Library Prep for Illumina kits (New England Biolabs kits E7490 and E7530), as per manufacturer’s instructions. Briefly, mRNA was enriched using oligo (dT) magnetic beads, fragmented, and reverse transcribed to cDNA. Samples were ligated with NEBNext multiplex oligo adaptor kits (New England Biolabs) for barcoding individual samples. Ligated DNA was then amplified by PCR. Libraries were quantified using a Qubit 2.0 fluorimeter and quality checked for insert size using a BioAnalyzer, then precisely quantified using KAPA Biosystems universal Illumina library quantification kit (Roche). Barcoded, prepared samples were pooled into two pools (with 10 and 14 individuals each) for sequencing on a HiSeq 4000 (Illumina) using paired-end (PE) 150 mode without PhiX with the aim to obtain approximately 10 million PE reads per sample. Samples from both DO conditions were present in approximate equal ratio in both lanes.

### Sequence read processing

All bioinformatics analyses are reported in detail including all scripts on GitHub (see *Data Accessibility*). Raw reads were quality checked with FastQC 0.11.5 (Andrews 2016), with individual sample files compiled using MultiQC 1.0 (Ewels *et al*. 2016). Adaptors, poor quality sequences (Phred = 2, MacManes 2014), and reads with a length less than 80 bp were removed using Trimmomatic v0.36 (Bolger *et al*. 2014) using flags *illuminaclip* (2:30:10), *slidingwindow* (20:2), *leading* (2) and *trailing* (2), and *minlen* (80). Trimmed reads were quality checked again with FastQC and MultiQC.

The Tuxedo protocol was followed for read alignments and assembling transcripts for quantification (Pertea *et al*. 2016). Trimmed PE reads were aligned to an indexed reference genome of Chinook salmon (Otsh_v1.0, RefSeq ID 6017098, Christensen *et al*. 2018) using the spliced aligner HISAT v2.1.0 (Kim *et al*. 2015) allowing up to 40 multi-mapping alignments (*k* = 40). The annotated reference genome contained 87,036 transcripts, and 82,000 intron chains. After alignment, the SAM file per individual was converted to a sorted BAM file with samtools v1.5 (Li et al. 2009). Alignments were then assembled into potential transcripts per individual with the reference genome as a guide using StringTie 1.3.4 (Pertea *et al*. 2015). Individual assemblies were merged to provide a final assembly, i.e. a combination of known transcripts from the reference genome annotation and novel transcripts not in the reference. The guided assembly was compared to the reference genome using *gffcompare* 0.10.4 from StringTie to identify novel features. This combined gff was then used to quantify read counts per transcript for each individual using StringTie. Counts per individual were merged into a single table and converted to transcript counts using the *prepDE.py* script provided by StringTie.

### Statistical analysis for discovering candidate hypoxia genes

After converting to gene counts, RNA-seq analyses were conducted using R 3.4.4 (R Core Team, 2018). The transcript counts per individual were analyzed using the *edgeR* v.3.7 package (Robinson *et al*. 2010) and general pipeline (Anders *et al*. 2013). Gene counts were filtered for low expressed genes using a 0.4 count-per-million (cpm) threshold, corresponding to a count of 5 reads in the sample with the fewest reads. A gene was removed if it was below this threshold in at least half the individuals (*n* = 12). Filtered counts were analyzed using multi-dimensional scaling (MDS) method to evaluate the effects of different factors (i.e., temperature, hypoxia, and trial) on overall gene expression. Then, the filtered counts were normalized and dispersions estimated using a model design containing the four treatments with no intercept. A robust quasi-likelihood generalized linear model was then fit to the counts. Given the potential issues of separating thermal stress from hypoxic stress responses (Pörtner 2010), quasi-likelihood F-tests used models with paired contrasts within temperatures, i.e., normoxia-10°C vs. hypoxia-10°C and normoxia-18°C vs. hypoxia-18°C. No differential expression was observed when the false discovery rate was applied (*q* < 0.05, Benjamini and Yekutieli 2001), and so for this discovery work, the unadjusted *p*-values were used. Differentially expressed genes (*p* < 0.05) in common for both contrasts were isolated using the *systemPipeR* package (Backman and Girke 2016) and retained if the log_2_ fold change direction was the same for both contrasts.

The subset of differentially expressed genes from above were subjected to a principal component analysis (PCA) using the scaled and normalized counts (cpm) of individuals. The PC axis associated with the separation of normoxia and hypoxia was identified. Candidate genes were ranked based on the significance of a Pearson correlation with this PC axis. The top 30 candidate genes based on correlation were examined for available mRNA sequences for several species of salmonids using a custom database (Akbarzadeh *et al*. 2018) and the public NCBI repository, as a goal of this work is to design primers that will work for multiple salmonid species.

### Development of RT-qPCR assays for validating hypoxia biomarkers

Universal salmonid (across nine salmonid species) real-time qPCR assays (Table S1) were developed for the top 30 candidate genes discovered using the RNA-seq data. The assay primers and TaqMan probes for the sequences were produced using Primer Express 3.0.1 (Thermo Fisher Scientific, Waltham, MA) with primer melting temperature (Tm) between 58–60 °C, and probe Tm between 68–70 °C as default.

To test the efficiency of the 30 discovered hypoxia genes across salmonid species, cDNA from RNA extractions of pooled tissues from each of the nine salmonid species, i.e. Chinook salmon (*O. tshawytscha*), coho salmon (*O. kisutch*), chum salmon (*O. keta*), pink salmon (*O. gorbuscha*), sockeye salmon (*O. nerka*), Atlantic salmon (*Salmo salar*), Artic charr (*Salvelinus alpinus*), rainbow trout (*O. mykiss*) and bull trout (*Salvelinus confluentus*), were serially diluted from 1/5 to 1/625 in five dilutions. Specific target amplification (STA), was performed to enrich targeted sequences within the pools using 3.76 μL 1X TaqMan PreAmp master mix (Applied Biosystems), 0.2 μM of each of the primers, and 1.24 μL of cDNA, as previously described (Akbarzadeh et al. 2018; Houde et al. 2019a). Samples were run on a 14 cycle PCR program, with excess primers removed with EXO-SAP-IT (Affymetrix), and then amplified samples were diluted 1/5 in DNA suspension buffer. The diluted samples and assays were run in singletons following the platform instructions (Fluidigm). For sample reactions, 3.0 μL 2X TaqMan mastermix (Life Technologies), 0.3 μL 20X GE sample loading reagent, and 2.7 μL STA product were used. For assay reactions, 3.3 μL 2X assay loading reagent, 0.7 μL DNA suspension buffer, 1.08 μL forward and reverse primers (50 uM), and 1.2 μL probe (10 uM) were used. The PCR was 50°C for 2 min, 95°C for 10 min, followed by 40 cycles of 95°C for 15 s, and then 60°C for 1 min. Data were extracted using the Real-Time PCR Analysis Software (Fluidigm) using Ct thresholds set manually for each assay. PCR efficiencies for each assay were calculated using (10^1/slope^ - 1) × 100, where the slope was estimated by plotting the Ct over the serial dilutions of cDNA.

Assay efficiencies ranged between 0.71 and 1.56 among species. Two assays were not amplified in all salmonid species, including *cyclin dependent kinase inhibitor 1B* (*CDKN1B*) and *receptor activity-modifying protein 1* (*RAMP1*) in rainbow trout, and *condensin-2 complex subunit G2* (*NCAPG2*) in chum and sockeye salmon. *Structural maintenance of chromosome protein 4* (*SMC4*) did not amplify in Chinook salmon and Arctic charr, and generally showed poor efficiencies across the remaining salmonid species. Therefore, *SMC4* was excluded from further analysis in the present study.

In total, 346 individuals were examined for gill gene expression. This number includes the RNA-seq samples from discovery analysis (n = 24) that were used for validating the transfer between platforms (RNA-seq to qPCR), and the extreme hypoxia samples that were either moribund or recently dead (n = 22). The rest of the data were comprised of 212 live-sampled juveniles from the four trials that had been exposed to the full treatment for six days, and 88 distressed juveniles (mostly from trials 1, 3, and 4, i.e. pre-smolt and de-smolt stages). The subset of juveniles (*n* = 43) used for analyzing the extreme hypoxia response were all from SW/18°C in trial 3: 11 live-sampled individuals exposed to extreme hypoxia for approximately two days, 11 distressed individuals exposed to extreme hypoxia, 11 live-sampled fish kept in normal oxygen for six days, and 11 live-sampled individuals exposed to hypoxia for six days. The technical roadmap of the experimental method is illustrated in Fig. 1.

**Fig. 1.**
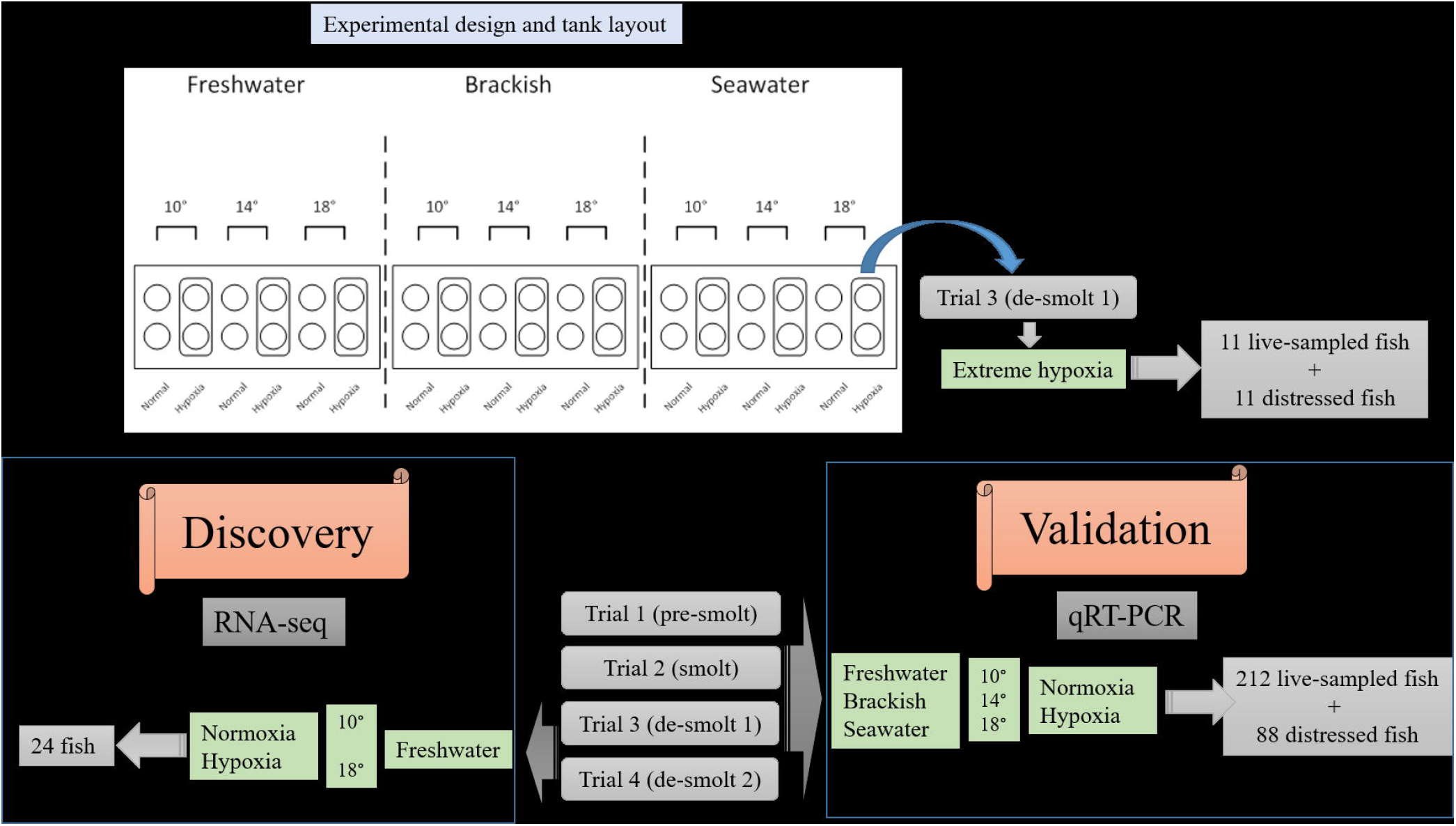
Technical road map of the experimental methods used for discovery (RNA-seq) and validation (q-PCR) of hypoxia biomarkers. Juvenile Chinook salmon were exposed to 18 possible water treatments in 30 L pot tanks (circles): three salinities (FW at 0 PSU, brackish (BW) at 20 PSU, and SW at 28 or 29 PSU), three temperatures (10, 14, and 18°C), and two dissolved oxygen (DO) concentrations (hypoxia 4□5 mg L^−1^ and normoxia > 8 mg L^−1^) in all combinations, each in two replicates. Four trials were conducted over the smoltification period, covering pre-smolt (March), smolt (May), and de-smolt 1 (June) and de-smolt 2 (August) stages. A subset of 24 FW individuals kept at two temperatures (10°C and 18°C) and two DO conditions (normoxia and hypoxia) was selected for RNA-seq from the two de-smolt trials (trials 3 and 4). A group of 212 live-sampled (no signs of morbidity), and 88 distressed juveniles (moribund and recently dead) from the four trials exposed to the full treatment were used for qRT-PCR. Twenty two individuals from two SW/18°C-hypoxia tanks in the trial 3 had lower DO than intended in the SW treatment (3.3–4.1 mg L^−1^ instead of 4–5 mg L^−1^), and therefore were used as an ‘extreme hypoxia’ condition.

Gill tissue homogenization, RNA extraction and quantification, cDNA synthesis, and STA were conducted as described above and previously (Houde *et al*. 2019a). The 96.96 gene expression dynamic array (Fluidigm Corporation, CA, USA) was applied and generally followed Miller *et al*. (2016). qRT-PCR data were analysed with Real-Time PCR Analysis 3 Software (Fluidigm Corporation, CA, USA). The expression of the 29 discovered hypoxia genes were normalized to the expression of the housekeeping gene, S100 calcium binding protein (78d16.1), which was found to be the most suitable housekeeping gene in NormFinder analysis, as previously described (Houde *et al*. 2019a). Sample gene expression was normalized with the ΔΔCt method (Livak and Schmittgen 2001) using the inter-array calibrator sample. Gene expression was then log transformed: log_2_ (2^−ΔΔCt^).

### Comparing RNA-seq results to the designed qRT-PCR targets

To validate the transfer of the discovered markers from RNA-seq platform to qRT-PCR, a correlation analysis was performed between the normalized counts (cpm) of RNA-seq data and the fold changes obtained from the qRT-PCR platform for the same 24 samples used for RNA-seq. Fold changes in expression values of RNA-seq data (on the X-axis) were plotted against fold change values obtained from RT-qPCR (on the Y-axis) on a scatter graph.

### Statistical analysis to validate hypoxia biomarkers

Analyses were performed using R 3.4.4 (R Core Team, 2018) at a significance level of α = 0.05. As the expression levels of hypoxia genes were also influenced by water salinity and smolt stage, further analyses were separately carried out for each group. The dataset for each group was divided into a two-thirds training set and a one-third testing set. The training set was subjected to a Shrunken Centroid method (Tibshirani et al. 2002) to identify a classifier for hypoxia vs. normoxia fish for each group. This method uses an internal cross-validation for threshold selection and returned a reduced list of the most influential genes for robust classification. Then, the selected genes were subjected to PCA analysis using the data from the training set that excluded the 24 RNA-seq samples. This PCA was then applied to the testing set for visualization of unsupervised group separation within each group using the *fviz_pca* function of the *factoextra* R package (https://cran.r-project.org/web/packages/factoextra/index.html). The classification ability was also examined by subjecting the resulting biomarkers to linear discriminant analysis (LDA) using the training set, followed by determining classification performance on the testing set. A similar approach was used for examining mortality (dead or moribund vs. live), and the extreme hypoxia group of SW-18°C in trial 3.

## Data availability

The analysis pipeline and scripts for the RNA-seq analysis is described in more detail on GitHub: https://github.com/bensutherland/Simple_reads_to_counts. The RNA-seq reads have been uploaded to the NCBI Sequence Read Archive (SRA) under BioProject PRJNA635140, with accession numbers SAMN15020676-SAMN15020699. https://www.ncbi.nlm.nih.gov/bioproject/PRJNA635140.

The qRT-PCR results and all the supplemental tables and figures are available at Figshare.

## Results

### Sequencing and gene count overview

Across the 24 individuals for RNA-seq, there was a sequencing depth range of 9.4 to 16.9 million trimmed PE reads (9.4□17.0 million raw reads). The overall alignment rate of the PE reads to the reference Chinook salmon genome ranged from 78.2 to 80.5% per sample. The final gene transcript annotation file (gff), comprised of merged reference guided individual transcript assemblies, contained 8.5% novel introns, 21.3% novel exons, and 56.7% novel loci. Using this final assembly, there were read counts for 127,887 transcripts. After the application of a low expression filter, counts were retained for 48,768 genes. The MDS plot using the overall filtered RNA-seq data showed that dimension 1 (dim 1) clearly separated samples kept in 10 vs. 18°C (i.e., large influence of temperature), and dimension 2 (dim 2) suggested separation of normoxia vs. hypoxia fish kept at 18 °C (Fig. 2). No trial effect was seen using the overall, filtered gene expression data (48, 768 genes).

**Fig. 2.**
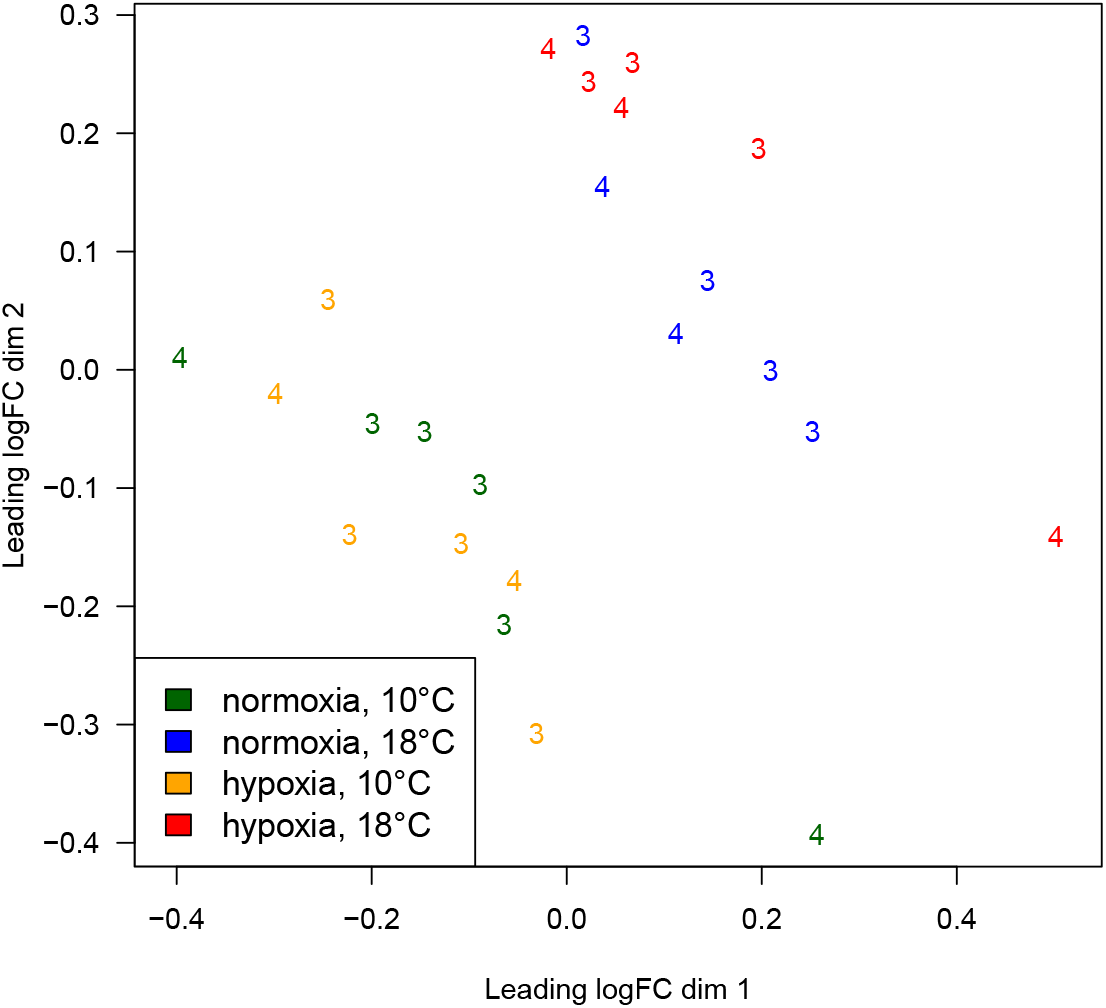
Multi-dimensional scaling (MDS) plot to evaluate the effects of different factors (i.e. temperature, hypoxia, trial) on overall gene expression (3= trial 3, de-smolt; 4= trial 4, de-smolt).

### Discovery of candidate hypoxia genes

The number of upregulated and downregulated genes (p < 0.05) between normoxia and hypoxia treatments was similar within each temperature: at 10 °C, there were 1,926 upregulated and 1,441 downregulated genes from hypoxia. At 18 °C, there were 1,742 upregulated and 1,347 downregulated genes from hypoxia. In total, 349 genes were found to be differentially regulated in both comparisons, and when requiring that these genes follow the same direction of regulation, this included 240 genes (i.e., 96 upregulated genes and 144 downregulated genes).

The PCA using only the 240 candidate hypoxia-response genes revealed that normoxia and hypoxia separation occurred along PC1, as expected given that these genes were selected based on their hypoxia response. There was also a separation of the 10 and 18°C groups along PC2 (Fig. 3). To avoid selecting genes responding to temperature in addition to hypoxia, genes were excluded when they were correlated with PC2 (*p* < 0.1; Table S2). Downregulated genes from hypoxia showed higher correlations with PC1 than did upregulated genes. The top 20 downregulated genes (*r* = −0.85 to −0.95) and the top 10 upregulated genes (*r* = 0.73 to 0.81) were investigated further for assay development. The top 20 down-regulated genes were mostly involved in DNA synthesis, cell division, and cell cycle, while the top 10 up-regulated genes were involved in the cell cycle arrest, oxidative stress response, ion transport, cellular calcium regulation, peptide amidation, and neuropeptide processing (Table 1).

**Fig. 3.**
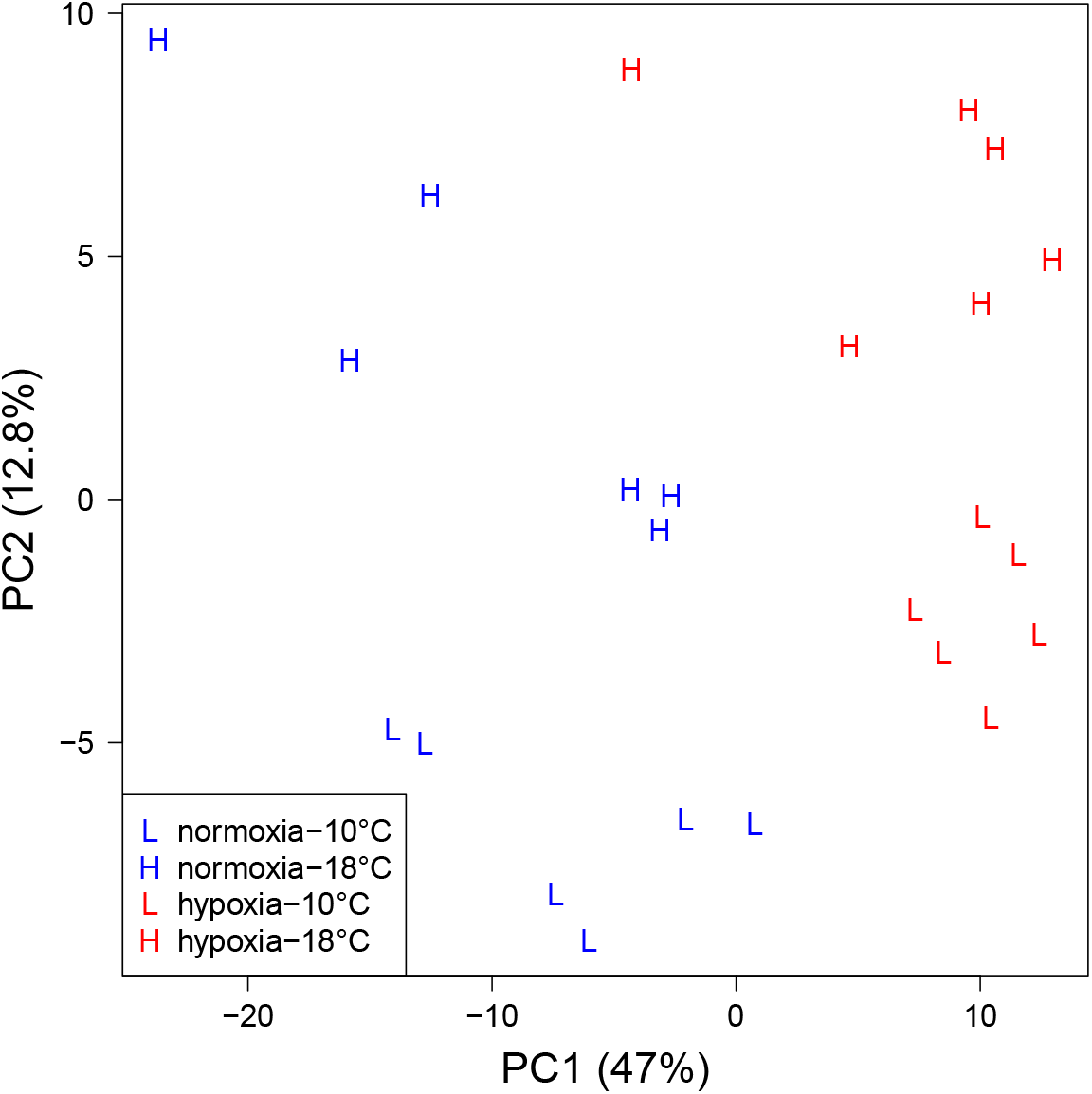
Canonical plot of the first two principal components of 240 candidate gill hypoxia genes.

**Table 1.**
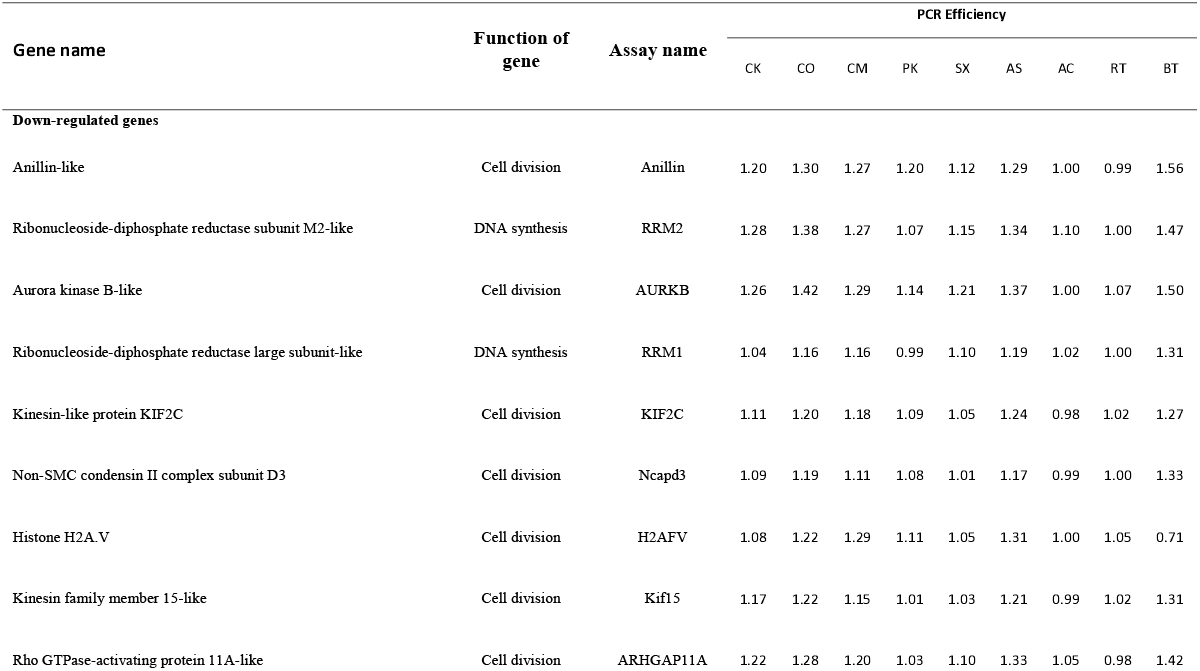
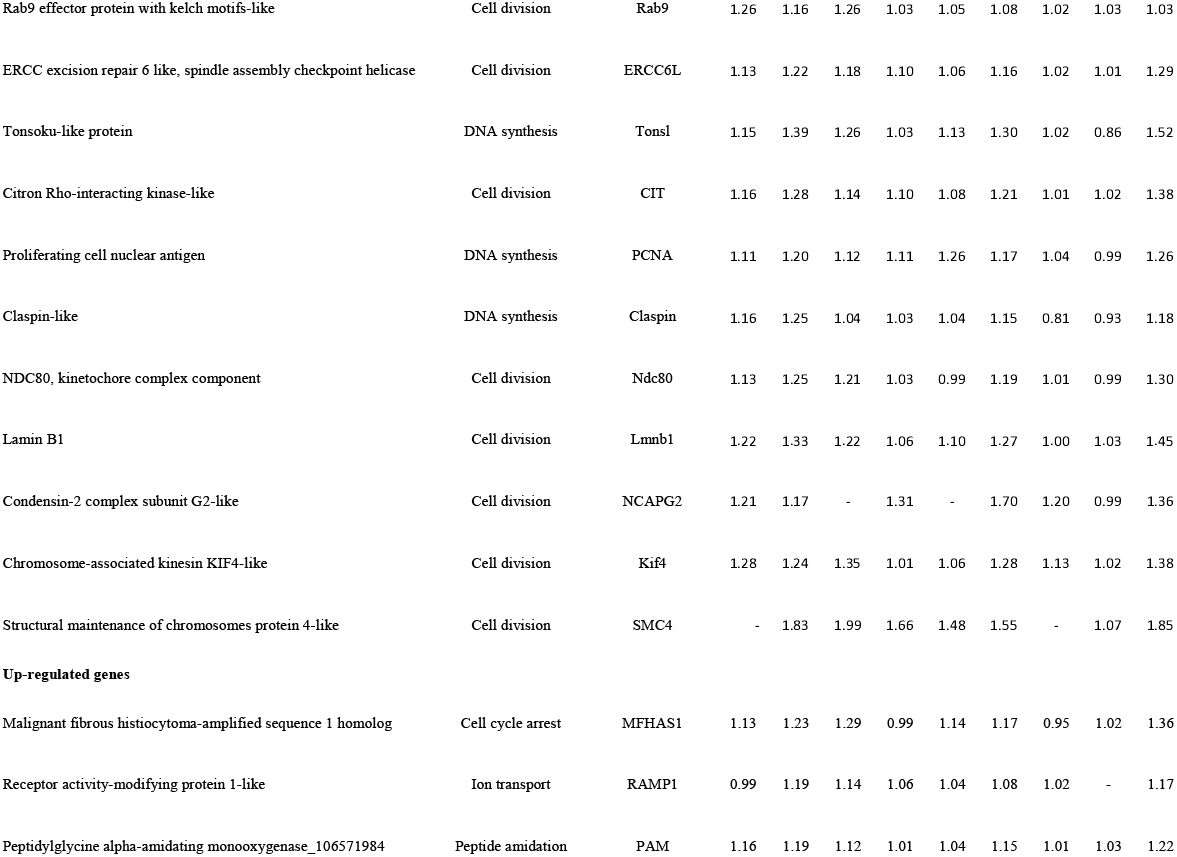
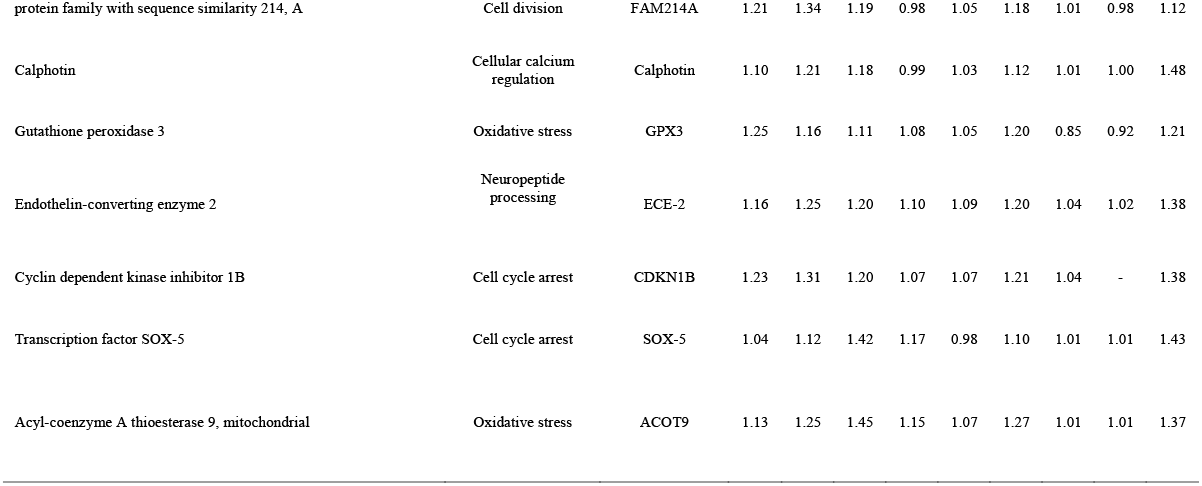
Top 20 downregulated and 10 upregulated candidate genes from the RNA-seq analysis, and efficiencies for candidate hypoxia stress genes. Genes are presented in order of the significance of the correlation with PC1, which separated normoxia and hypoxia (data are in Table S1). Species abbreviations for efficieny: CK= Chinook salmon (*Oncorhynchus tshawytscha*), CO= Coho salmon (*O. kisutch*), CM= Chum salmon (*O. keta*), PK= Pink salmon (*O. gorbuscha*), SX= Sockeye salmon (*O. nerka*), AS= Atlantic salmon (*Salmo salar*), AC= Artic charr (*Salvelinus alpinus*), RT= Rainbow trout (*O. mykiss*), and BT= Bull trout (*Salvelinus confluentus*).

### Correlation between RNA-seq and qRT-PCR data

The validation of the design of the primers for qPCR compared to the results from RNA-seq was conducted using a correlation analysis between the log_2_ fold change from the RNA-seq data and the log_2_ fold change of the qRT-PCR platform for the same 24 samples used for both platforms. A high value for the Pearson correlation coefficient (r = 0.88; p < 0.001) indicated a positive correlation between platforms (Fig. 4). From 29 genes tested, 15 genes showed significant correlation (*p* < 0.05) between two platforms (Table S3).

**Fig. 4.**
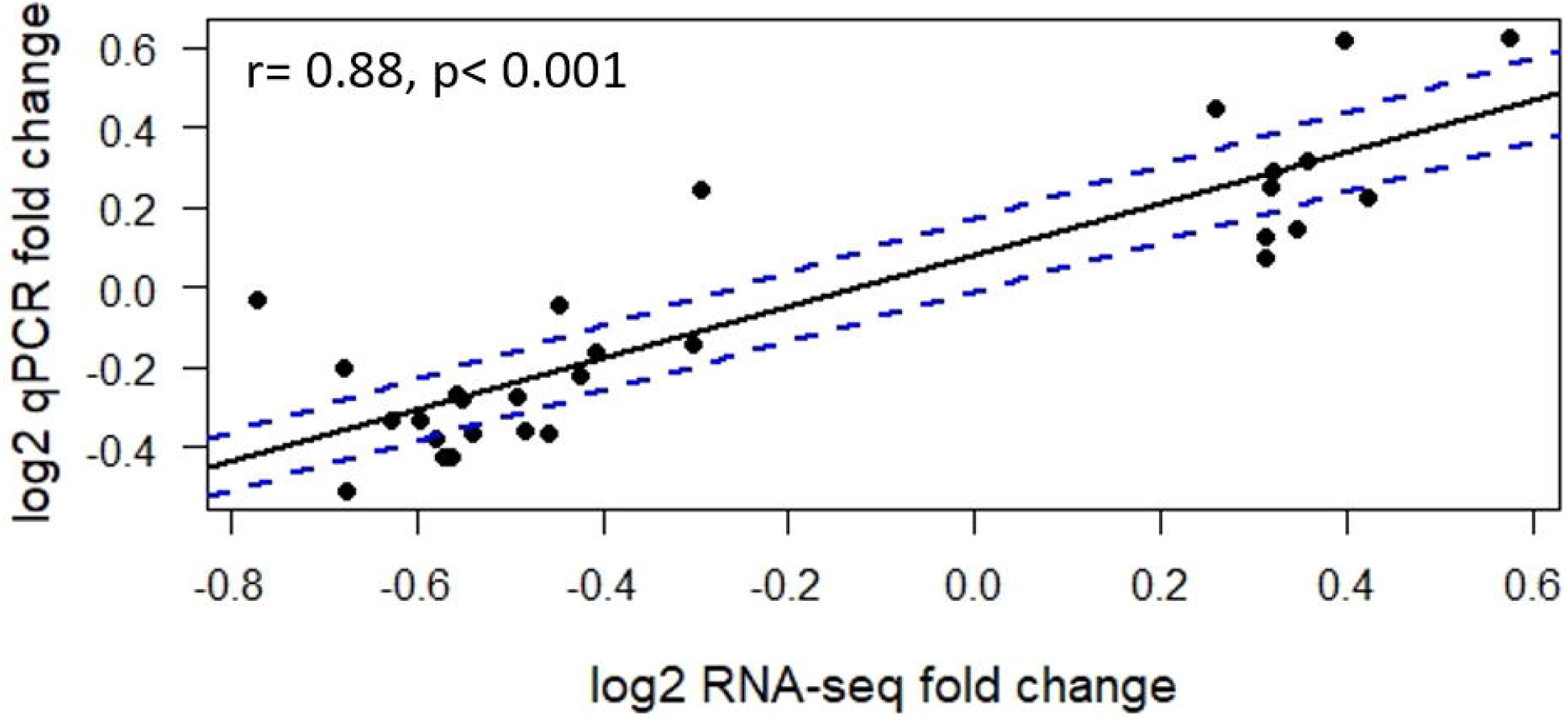
Validating RNA-seq platform with qRT-PCR data using the same 24 samples used in RNA-seq. Fold changes of gene expression detected by RNA-seq were plotted against the data of qRT-PCR. The reference line indicates the linear relationship between the results of RNA-seq and qRT-PCR.

Moreover, there was a high separation between hypoxia and normoxia along PC1 using the gene expression data of 24 independent RNA-seq samples (Fig. 5). Using the top seven genes obtained from the Shrunken Centroid method on 15 significant correlated genes, the classification ability for normoxia was 100.0% and hypoxia was 75.0% with the average classification of 87.5%.

**Fig. 5.**
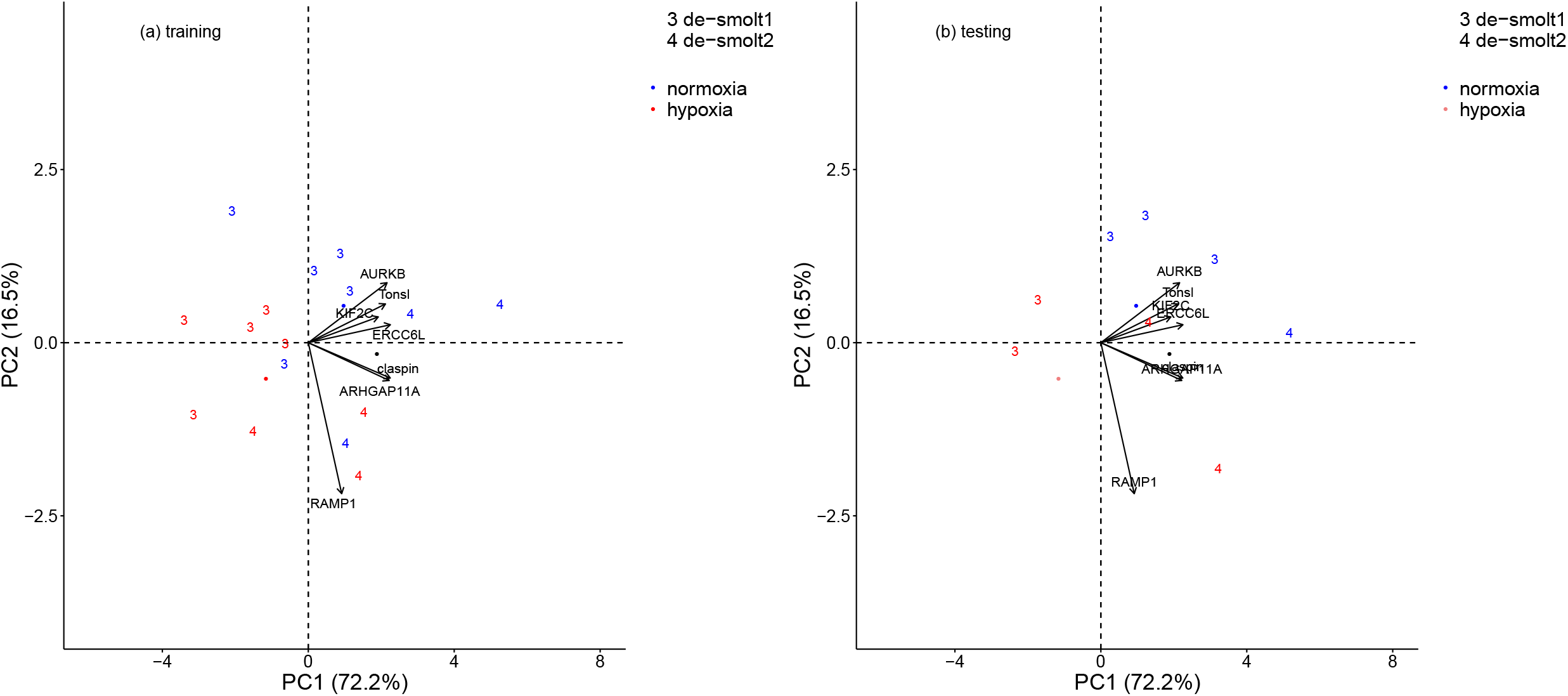
Canonical plots of the first two principal components of the identified hypoxia biomarkers using the 7 hypoxia biomarkers returned by the Shrunken Centroid method on significant correlated genes between RNA-seq and qRT-PCR platforms for 24 samples used in RNA-seq analysis. Temperature treatment symbols are 10 and 18 °C groups. Principal component analysis was performed on the training set (left panel) and then applied to the testing set (right panel). Centroids are represented by the largest point of the same colour. Arrows represent loading vectors of the biomarkers using the training set.

### Hypoxia biomarkers

The qRT-PCR gene expression results for the 29 discovered hypoxia genes including all 300 live-sampled and distressed fish kept at different combinations of temperature, salinity and oxygen for six days, as well as 22 live-sampled and distressed fish exposed to extreme hypoxia for two days are presented in Fig. S1. Moreover, the initial results of the PCA analysis on all live-sampled and distressed fish together using the selected genes from Shrunken Centroid method showed that PC1 was associated with smolt stage, water salinity, and mortality (*data not shown*). Given the influence of smolt stage, salinity, and mortality on the candidate hypoxia biomarkers, we separated the data first by the three smolt stages (live-sampled only) and distressed, and then by the three salinities such that there were 11 groups. Distressed fish in BW data was not analyzed due to too few samples (n = 3) of normoxia fish in this salinity environment.

The results of the PCA analyses and LDA scores for normoxia vs. hypoxia within each group using the most contributing hypoxia biomarkers are shown in Figures S2-S5 and Table 2. Effectiveness of biomarkers were evaluated based on the ability to separate samples within the PCA and low LDA test error for both normoxic and hypoxic conditions.

**Table 2.**
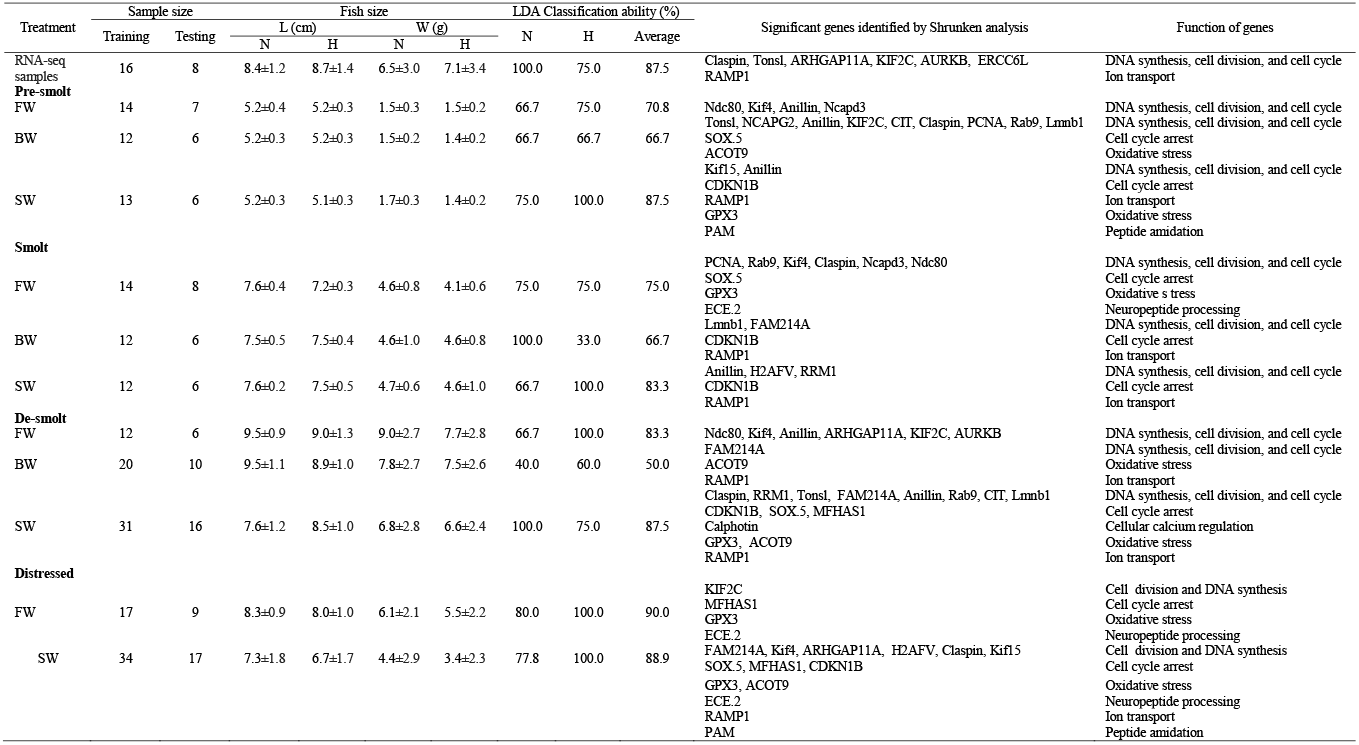
Classification ability of the normoxia and hypoxia groups for 24 samples used in RNA-seq, and across the smoltification process of Chinook salmon, and the distressed (moribund/recently dead) fish using the hypoxia biomarkers. Biomarkers identified by Shrunken Centroid method for each group were placed into a linear discriminant analysis (LDA) for classifying groups within each treatment and mortality. LDA training set used two-thirds of the entire dataset. Presented are the classifications on the remaining one-third testing set. Columns are the LDA classification groups and rows are the real group. Salinity symbols are FW for freshwater, BW for brackish, and SW for seawater groups. Dissolved oxygen symbols are N for normoxia and H for hypoxia groups. Fish size (mean ± standard deviation) symbols are L for length and W for weight.

Pre-smolt fish transferred to SW showed a distinct separation of normoxia and hypoxia groups along PC2, having the highest average LDA classification accuracy (87.5%) (Fig. S2a; Table 2). The pre-smolts in SW had higher LDA classification accuracy than the LDA results for fish in FW or BW.

Smolts in FW or SW had distinct separations of normoxia and hypoxia samples along PC2 (Fig. S3a, b). The SW smolts showed the highest LDA classification accuracy (83.3%), although this was biased towards identifying samples in the hypoxia condition (Table 2). The most balanced LDA classification between hypoxic and normoxic classifications in the smolts was 75% as observed in smolts in FW (Table 2).

De-smolt fish in FW had a distinct separation for normoxia and hypoxia groups along PC2 with an average LDA classification accuracy of 83.3% but biased towards hypoxia (Fig. S4; Table 2). Although the highest average LDA classification accuracy for de-smolts was 87.5% (in SW), there was no clear separation for the PCA using the first two axes (Table 2). Moreover, both trials of de-smolt fish were also separated along PC2 in all salinity environments (Fig. S4).

Distressed fish in both FW and SW showed the highest PCA separation and classification ability with an average classification accuracy of 90.0 and 88.9%, respectively (Fig. S5; Table 2).

The PCA analysis on the extreme hypoxia group of SW/18°C in trial 3 showed a stronger separation of normoxia and live-extreme hypoxia exposed fish than normoxia and moderate hypoxia groups along PC1. Distressed extreme hypoxia exposed fish were also separated from other groups in PCA analysis (Fig. 6).

**Fig. 6.**
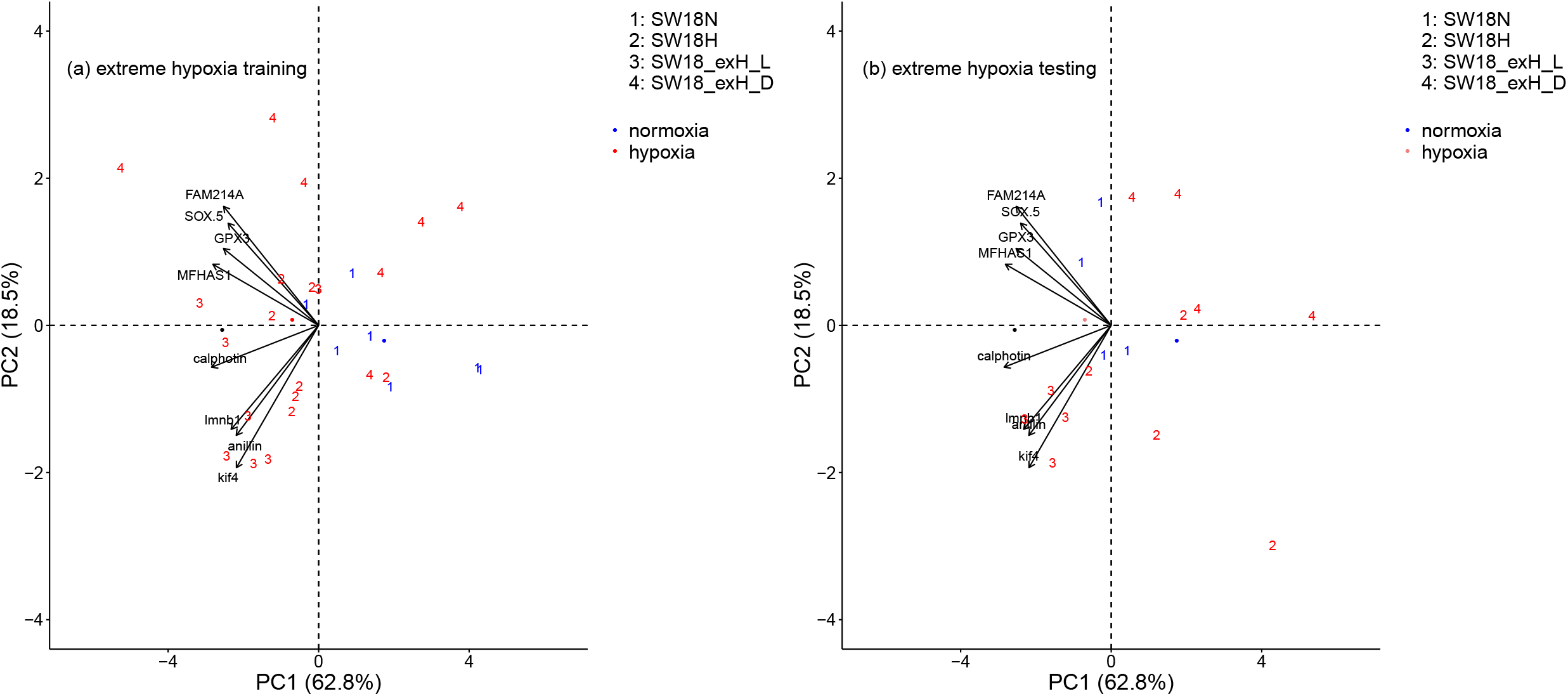
Canonical plots of the first two principal components using 8 hypoxia biomarkers returned by the Shrunken Centroid method for the extreme hypoxia group of SW/18°C in trial 3. 11 live individuals exposed to extreme hypoxia for approximately two days (SW18_exH_L), 11 dead or moribund individuals exposed to extreme hypoxia (SW18_exH_D), 11 live fish kept in normal oxygen for six days (SW18N), and 11 live individuals exposed to designed hypoxia for six days (SW18H). Principal component analysis was performed on the training set (left panel) and then applied to the testing set (right panel). Centroids are represented by the largest point of the same colour. Arrows represent loading vectors of the biomarkers using the training set.

In sum, the conditions that showed the most potential for separating fish kept in normoxic from hypoxic conditions were pre-smolt and smolt stages in SW, and de-smolt stage in FW. On the contrary, all conditions in BW, pre-smolt and smolt in FW, and de-smolt in SW were not able to clearly separate fish kept in normoxic from hypoxic conditions.

## Discussion

Characterizing early genetic responses to hypoxia to facilitate the development of a tool to assess hypoxia exposure is crucial towards managing salmonid fish that are increasingly exposed to hypoxic stress globally. This is particularly relevant considering the strong influence anthropogenic effects, such as excessive nutrient loading, can have on hypoxia levels and occurrence, for example on the west coast of North America. The present study aimed to discover and validate gill gene expression biomarkers specific to hypoxic stress in salmonids. Using high-throughput RNA-seq analysis, we discovered 30 candidate genes that were differentially expressed between hypoxic and normoxic conditions, but were not affected by temperature. The discovered genes were then designed to qRT-PCR assays, and then used to query a larger set of samples by high-throughput microfluidics qPCR. These qPCR data were then used to characterize and identify the best biomarkers for differentiating exposure to normoxic and hypoxic conditions.

### Biological and molecular functions of candidate hypoxia genes

It should be noted that while transcript changes are expected to reflect protein and physiological effects, a strong correlation between gene and protein expression is not always the case owing to post-transcriptional and post-translational modifications (Maier *et al*. 2009; Schwänhausser *et al*. 2011; Kanerva *et al*. 2014). However, here we discuss transcripts in relation to their expected protein products and functions. All of the top 20 downregulated genes in the hypoxia treatment discovered by the RNA-seq analysis were involved in functions related to DNA synthesis, cell division, and cell cycle. The concordant downregulation of these genes during hypoxia suggests that cell proliferation may be suppressed in the Chinook salmon gill tissue. Hypoxia has been shown to inhibit cell proliferation in numerous cell types (Hubbi and Semenza 2015). A decreased number of cells may allow for decreased oxygen demand under hypoxia (Hubbi and Semenza 2015). These hypoxia-suppressed genes may reflect a physiological alteration that halts cell proliferation and translational activities to allow the cells to preserve energy consumption under metabolic stress (Chi *et al*. 2006). The downregulation of genes involved in cell cycle progression occurs in response to acute and chronic hypoxia in zebrafish embryos (Ton *et al*. 2003) and in hepatic cells of hybrid striped bass (*Morone saxatilis* × *Morone chrysops*) (Beck *et al*. 2016). Therefore, the downregulation of cell proliferation in the gill may be a physiological strategy in Chinook salmon to cope with hypoxic water.

Consistent with the suppression of genes involved in cell proliferation, theexpression of genes involved in blocking the cell cycle progression i.e., *transcription factor SOX-5 (SOX-5), CDKN1B*, and *malignant fibrous histiocytoma-amplified sequence 1 homolog* (*MFHAS1*) were among the top 10 hypoxia-induced upregulated genes discovered in this study. Sox-5 is well known for its function in controlling cell cycle and sequential generation of distinct corticofugal neuron subtypes (Hao *et al*. 2014). The upregulation of *SOX-5* occurs in response to hypoxia in human stem cells (Khan *et al*. 2007). Moreover, the upregulation of *CDKN1B*, which blocks cell cycle progression from G1 to S phase, is also a well-known response to hypoxia (Wang *et al*. 2003; Peng *et al*. 2015). MFHAS1 is known as an oncogene that may play an important role in signal transduction and the control of cell growth in humans (Sakabe *et al*. 1999). Collectively, this suggests that hypoxia may elevate expression of genes involved in inhibiting cell growth and proliferation, thus potentially reducing growth rates. Furthermore, this is in line with the lower growth performance of these juveniles kept in hypoxia compared to normoxia (see Houde *et al*. 2019a).

Top 10 hypoxia-induced upregulated genes included those involved in the oxidative stress response (e.g., *glutathione peroxidase 3; GPx-3*, and *Acyl-coenzyme A thioesterase 9, mitochondrial; ACOT*), ion transport (e.g., *RAMP1*), peptide amidation (e.g., *peptidylglycine alpha-amidating monooxygenase; PAM*), and neuropeptide processing (e.g., *endothelin-converting enzyme 2; ECE-2*). GPx-3 is a selenocysteine-containing protein with antioxidant properties. Hypoxia is known as a strong transcriptional regulator of *GPx-3* expression. Increased expression of oxidative stress genes under hypoxic conditions has been already demonstrated in fish (Zhang *et al*. 2016). RAMP1 is a member of the RAMP family of single-transmembrane-domain proteins, called receptor (calcitonin) activity modifying proteins (RAMPs). Chronic hypoxia was reported to enhance mRNA levels for *RAMP1* in rats (Qing *et al*. 2001; Cueille *et al*. 2005). It is stated that in hypoxia, upregulation of *RAMP1* primarily regulates the calcitonin gene-related peptide reception (Qing and Keith 2003). Moreover, PAM catalyzes the amidation of peptides, a process important for peptide stability and biological activity. PAM-dependent amidation has the potential to signal oxygen, so that PAM is strikingly sensitive to hypoxia in cells (Sharma *et al*. 2009; Simpson *et al*. 2015). PAM has catalytic activity on a range of substrates, in different cell types, and is as sensitive to hypoxia as the transcription factor, hypoxia inducible factor (HIF) (Simpson *et al*. 2015). *ECE-2* has been observed to be upregulated in response to hypoxia in juvenile salmon in this study. ECE-2 belongs to a family of membrane-bound metalloproteases involved in neuropeptide processing (Mzhavia *et al*. 2003). The true physiological function of ECE-2 remains largely to be determined (Ouimet *et al*. 2010), but it may be a novel HIF-target gene in hypoxia, where HIF-1 binds to transcription regulatory regions of the *ECE-1* gene during hypoxia (Khamaisi *et al*. 2015). However, no information is available regarding this mechanism for the *ECE-2* gene. The upregulation of *GPX3, Sox-5, CDKN1B*, and *ECE-2* observed in this study was in line with the higher expression of the transcription factor *HIF-1A* (see Houde *et al*. 2019a) in response to hypoxia. Notably, and in line with observations from the current study in Chinook salmon, HIF-1A is a key regulator of the cellular and systemic homeostatic response to hypoxia through the activation of transcription of many genes, including those involved in cell cycle arrest (i.e. *CDKN1B*), antioxidant activity (i.e. *GPX3*), and other genes whose protein products increase oxygen delivery or facilitate metabolic adaptation to hypoxia (Lando *et al*. 2003; Wang *et al*. 2003; Zagórska and Dulak 2004). A recent transcriptome study in a catfish (*Pelteobagrus vachelli*) identified 70 candidate genes in the HIF-1 signaling pathway in response to acute hypoxia and reoxygenation (Zhang *et al*. 2017).

### Differing effects dependent on smolt stage and environmental salinity

Juvenile Chinook salmon exposed to normoxia and hypoxia under various conditions were separated to different levels of success using the biomarkers developed here. Gene expression patterns and multivariate analyses using the discovered hypoxia genes showed that at the level of hypoxia induced in our study, the hypoxia response in juvenile Chinook salmon was highly affected by the developmental process of three factors: smoltification (i.e., smolt stage), water salinity, and mortality status. It is known that metabolic responses can be affected by both smolt stage and salinity in salmon (Morgan and Iwama 1991; Hvas *et al*. 2018).

Across all smolt stages, the results showed lower classification ability of fish in BW than in FW or SW. It is known that salmon in BW are near their isosmotic environment (i.e., the gradient between blood and environmental salinity is lower), such that the oxygen consumption and metabolic rate or energy expenditure may be lower than that in FW or SW to maintain ion- and osmo-regulation (Morgan and Iwama 1991, 1998). These juveniles may have better aerobic performance and may experience less hypoxic stress in this environment (Hvas *et al*. 2018). This was in line with our previous observation of lower Chinook salmon juvenile mortality in BW than in either FW or SW (Houde *et al*. 2019a). Altogether, these results suggest that juvenile Chinook salmon are more resilient to low oxygen levels in brackish conditions than in either full seawater or freshwater.

Pre-smolt Chinook salmon were responsive to hypoxia in SW but not in FW. Juvenile salmonids that are physiologically unprepared for the transition from FW to SW (i.e., pre-smolts) experience higher mortality in SW (Houde *et al*. 2019a,b). Pre-smolts in SW, as opposed to FW, may display higher oxygen consumption as a stress response, and higher energy expenditure to maintain ion- and osmo-regulation (Morgan and Iwana 1991, 1998). Furthermore, oxygen consumption typically is the lowest in the environment that is natural for a given life stage (Ern *et al*. 2014). It is likely that pre-smolt juvenile Chinook salmon in FW experienced lower physiological disturbance than those in SW, such that they were more prepared to cope with hypoxic conditions in FW than in SW.

Smolt and de-smolt juvenile Chinook salmon were responsive to hypoxia in both FW and SW. During smoltification, physiological adaptations to SW increase and adaptations to FW decrease. For example, the gill gene expression of SW and FW isoforms of *Na^+^/K^+^-ATPase* follows this pattern (Nilsen *et al*. 2007). De-smoltification occurs when smolts do not migrate to SW within their smolt window, and remain in FW too long, such that they revert to a more FW-adapted physiology. Although certain physiological features in de-smolts do revert back to FW forms, including ion- and osmo-regulation, other features remain in the SW form, e.g., higher metabolic rate (McCormick *et al*. 1998). Considering the lower oxygen requirements in the most natural environment for a particular life stage (Ern *et al*. 2014), it is possible that smolts and de-smolts both may be less able to cope with hypoxic conditions in FW and SW because they are not fully adapted to either environment.

The post-smolt stage was not examined in the present study but may reveal further insights on the response to hypoxia by salinity and smolt stage. De-smolts may occur in the hatchery setting, whereas smolts in the natural setting typically transition to SW and become post-smolts (McCormick *et al*. 1998). Therefore, the hypothesis of lower oxygen consumption or energetic cost in the natural salinity environment for a given life stage (Morgan and Iwana 1991, 1998; Ern *et al*. 2014), may be further examined using post-smolts, and this would be a valuable direction for future work. It is predicted that the post-smolt pattern may be similar to pre-smolts but in the opposite direction, i.e., post-smolts are more prepared to cope with hypoxic conditions in SW than in FW. Indeed, aerobic scope, but not standard or maximum metabolic rate, is lowest for post-smolt Atlantic salmon in SW than FW and BW (Hvas *et al*. 2018). Further studies examining post-smolts using the three salinities and the candidate hypoxia genes are warranted.

Across smolt stages and salinities, our results showed that three genes involved in cell cycle and DNA synthesis (i.e. *anillin, CDKN1B*, and *Kif4*) were among the most universal genes differentially expressed in response to hypoxia. However, some genes involved in ion transportation (i.e. RAMP1) were differentially expressed in saline water (i.e. brackish water and seawater). In pre-smolt fish kept in FW and BW, the hypoxia responsive genes were mostly involved in cell cycle and DNA synthesis, whereas in SW, the genes involved in ion transportation and oxidative stress also contributed to the hypoxia response. The smolt and de-smolt fish also showed differences in ion transportation genes related to hypoxia when in SW conditions. Previous studies demonstrated that the hypoxia exposure of gill in fish is associated with an ionoregulatory disturbance. Indeed, adjustments of gill morphology, which includes the morphological changes in the gills, along with possible changes in gill ventilation and perfusion to secure oxygen uptake, have a negative effect on the ionoregulatory status of the fish (Matey *et al*. 2008). Therefore, our results suggest that hypoxia may also affect the expression of osmoregulation and ionoregulation genes in gill tissue, which is important given that this tissue is the primary organ involved in both osmoregulation and ionoregulation (Houde *et al*. 2019b).

### Mortality

The results of this study showed that the top candidate hypoxia genes were highly responsive to hypoxia in moribund and recently dead fish in both FW and SW environments, suggesting the active regulation of hypoxia genes in moribund and recently dead fish. These results confirm previous findings of post-mortem activity of hypoxia responsive genes in human (Ferreira *et al*. 2018), mouse and zebrafish (Pozhitkov *et al*. 2016). It has been previously suggested that the genes that function in DNA synthesis, deactivation of immune system, cell necrosis, stress response, and glycolysis of carbohydrates are also the major group of genes that show changes in transcription following death. Hypoxia is believed to play a major role in the initial pre- to post-mortem transition by activation of glycolysis (Ferreira *et al*. 2018). In our study, genes involved in cell cycle and DNA synthesis, oxidative stress, and neuropeptide processing were among the most significant contributed genes responsive to hypoxia in distressed fish in both FW and SW environments. The cellular hypoxic response to morbidity is likely due to reduced blood circulation and consequently a considerable reduction of available oxygen in cells. Our validated hypoxia biomarkers may be applicable to elucidating the causes of salmon mortality and the environmental habitats where salmon experience episodes of hypoxia. Although further confirmatory studies are required, these results demonstrate that this high-throughput gene expression approach may have a high potential for being a useful tool to discover hypoxic mortality in salmonids, and the synergistic relationships between hypoxic stress and disease, if applied on salmon Fit-Chips.

### Extreme hypoxia

The most robust hypoxia response was observed in individuals exposed accidently to more severe hypoxic conditions in SW/18°C than the fish exposed to the more moderate hypoxia conditions designed in the original study as shown by the PCA analysis; this may suggest that the moderate hypoxia conditions were not strong enough to induce a consistent hypoxia response across all individuals. These results were obtained on a limited number of fish due to an instrument issue during laboratory handling. Conceivably, a stronger hypoxia across all treatments may have shown a more robust hypoxia response in salmon, but stronger hypoxia was not chosen because of ethical considerations. In addition, the RNA-seq analysis across the transcriptome revealed small fold-changes and no significant genes using false discovery rate (q-values), supporting a weak hypoxia response using the experimental conditions, although this is probably also impacted by sample size in the discovery component of this study. Further experiments using stronger hypoxic conditions across different salinities, temperatures, and smolt stages are needed to confirm this hypothesis.

## Conclusions

Gill gene expression biomarkers have been previously identified for salinity and temperature stressors across multi-stressor conditions, but hypoxia biomarkers sufficient to identify fish exposed to moderate hypoxia (Houde et al. 2019a). Here, candidate hypoxia genes were discovered through RNA-seq analysis, and then validated using microfluidics qPCR to reveal sets of biomarkers that were responsive to hypoxic stress in juvenile Chinook salmon. Moreover, the expression of these candidate hypoxia genes, and presumably of the hypoxia response, in juvenile Chinook salmon was affected by smolt stage, induced mortality, and water salinity. Our study is the first of its kind dealing with this range of factors from the gene expression perspective. Following the biomarker discovery of 30 candidates by RNA-seq, different validated biomarker sets were identified that were optimal for each of the specific combinations of factors within the dataset (i.e., smolt stage, mortality, salinity). The genes involved in cell cycle and DNA synthesis were among the most universal biomarkers, whereas ion transportation genes were more specific signals within saline water. Since the biomarkers were discovered on specific water salinity levels, smolt stages and water temperatures, no consistent gene expression model for all conditions was observed at the level of hypoxia exposure conducted in our study. Interestingly, this study also provides some evidence for post-mortem regulation of hypoxia genes in fish. The main challenge facing the development of a hypoxia gene expression biomarker panel for salmonids appears to be the relatively minor response compared to other stressors such as temperature, which made the removal of temperature response genes in this study an important part of the design. The present study provides a set of biomarkers that, when combined with factors smolt stage, salinity, and mortality, can be used to determine whether an individual has recently experienced hypoxic stress. Moreover, our study confirms that fish might be highly resilient to mild hypoxia when they are in their natural salinity environment, less so when outside of the salinity environment expected for that development stage, and can be stressed by extreme hypoxia conditions. The biomarkers identified here will be added to the continued development of stressor-specific biomarkers being collected to understand the abiotic and biotic stressors that salmonids face in their natural environment.

## Acknowledgements

Funding for this research was provided by the Pacific Salmon Commission, Pacific Salmon Foundation, and Fisheries, Oceans and the Canadian Coastguard (DFO) Genomics Research and Development Initiative (GRDI) Fund to KMM. We thank DFO Aquarium services for help in the design and assembly of the experiment. We also thank X. He for gene expression analysis advice.

## Notes

https://github.com/bensutherland/Simple_reads_to_counts

### Competing Interest Statement

The authors have declared no competing interest.

### Summary of Updates

Identifying early gene expression responses to hypoxia (i.e., low dissolved oxygen) as a tool to assess the degree of exposure to this stressor is crucial for salmonids, because they are increasingly exposed to hypoxic stress due to anthropogenic habitat change, e.g., global warming, excessive nutrient loading, and persistent algal blooms. Our goal was to discover and validate gill gene expression biomarkers specific to the hypoxia response in salmonids across multi-stressor conditions. Gill tissue was collected from 24 freshwater juvenile Chinook salmon (Oncorhynchus tshawytscha), held in normoxia [dissolved oxygen (DO) > 8 mg L-1] and hypoxia (DO = 4‒5 mg L-1) in 10 and 18 C temperatures for up to six days. RNA-sequencing (RNA-seq) was then used to discover 240 differentially expressed genes between hypoxic and normoxic conditions, but not affected by temperature. The most significantly differentially expressed genes had functional roles in the cell cycle and suppression of cell proliferation associated with hypoxic conditions. The most significant genes (n = 30) were selected for real-time qPCR assay development. These assays demonstrated a strong correlation (r = 0.88; p < 0.001) between the expression values from RNA-seq and the fold changes from qPCR. Further, qPCR of the 30 candidate hypoxia biomarkers was applied to an additional 322 Chinook salmon exposed to hypoxic and normoxic conditions to reveal the top biomarkers to define hypoxic stress. Multivariate analyses revealed that smolt stage, water salinity, and morbidity status were relevant factors to consider with the expression of these genes in relation to hypoxic stress. These hypoxia candidate genes will be put into application screening Chinook salmon to determine the identity of stressors impacting the fish.

