## Supplemental Fig. 1 for "Identification of hypoxia-specific biomarkers in salmonids using RNA-sequencing and validation using high-throughput qPCR"

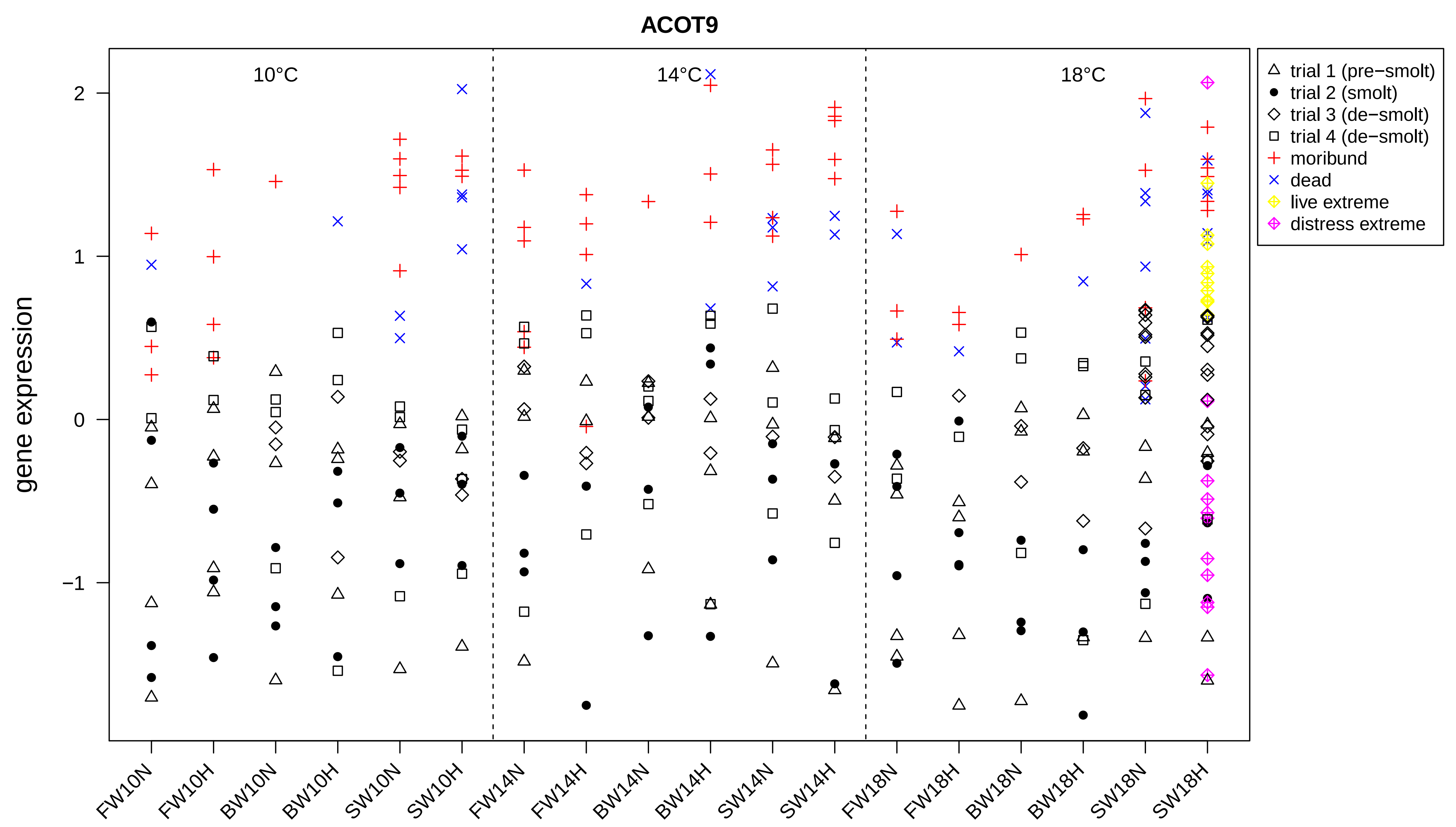

ACOT9
10°C
14°C
18°C
gene expression
2
1
0
−1
FW10N
FW10H
BW10N
BW10H
SW10N
SW10H
FW14N
FW14H
BW14N
BW14H
SW14N
SW14H
FW18N
FW18H
BW18N
BW18H
SW18N
SW18H
trial 1 (pre−smolt)
trial 2 (smolt)
trial 3 (de−smolt)
trial 4 (de−smolt)
moribund
dead
live extreme
distress extreme

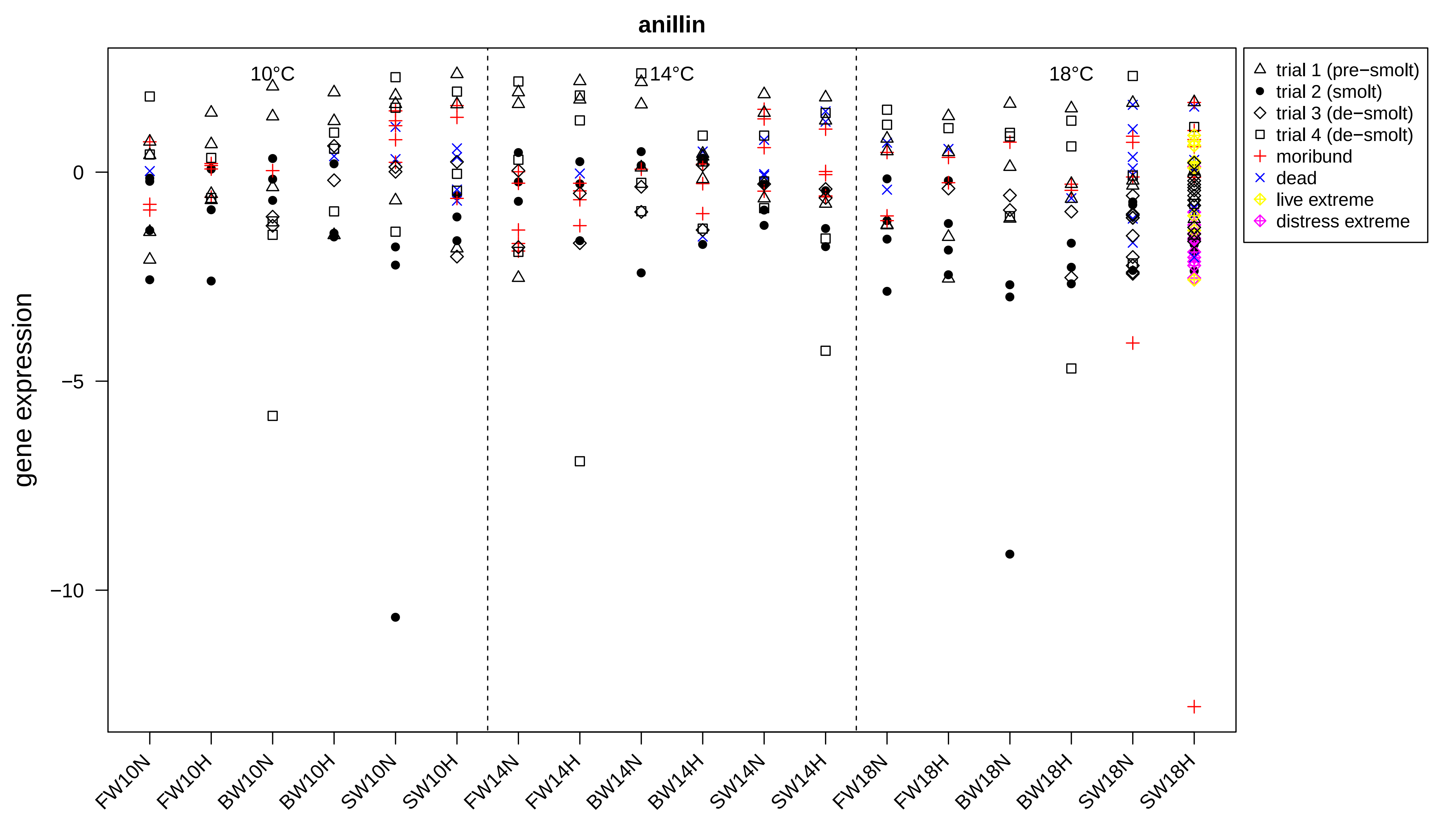
anillin
10°C
14°C
18°C
gene expression
0
−5
−10
FW10N
FW10H
BW10N
BW10H
SW10N
SW10H
FW14N
FW14H
BW14N
BW14H
SW14N
SW14H
FW18N
FW18H
BW18N
BW18H
SW18N
SW18H
trial 1 (pre−smolt)
trial 2 (smolt)
trial 3 (de−smolt)
trial 4 (de−smolt)
moribund
dead
live extreme
distress extreme

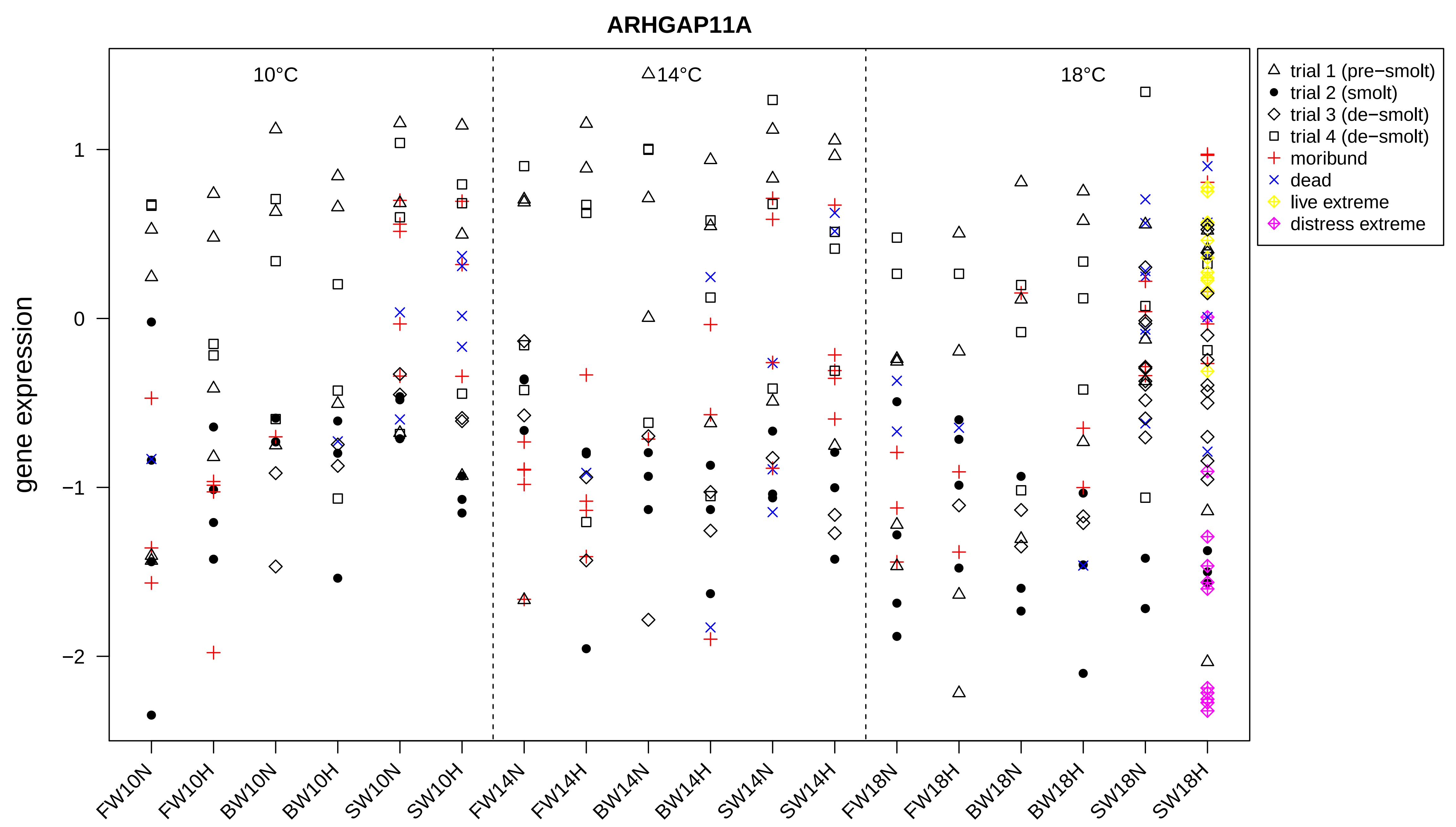
ARHGAP11A
10°C
14°C
18°C
gene expression
1
0
−1
−2
FW10N
FW10H
BW10N
BW10H
SW10N
SW10H
FW14N
FW14H
BW14N
BW14H
SW14N
SW14H
FW18N
FW18H
BW18N
BW18H
SW18N
SW18H
trial 1 (pre−smolt)
trial 2 (smolt)
trial 3 (de−smolt)
trial 4 (de−smolt)
moribund
dead
live extreme
distress extreme

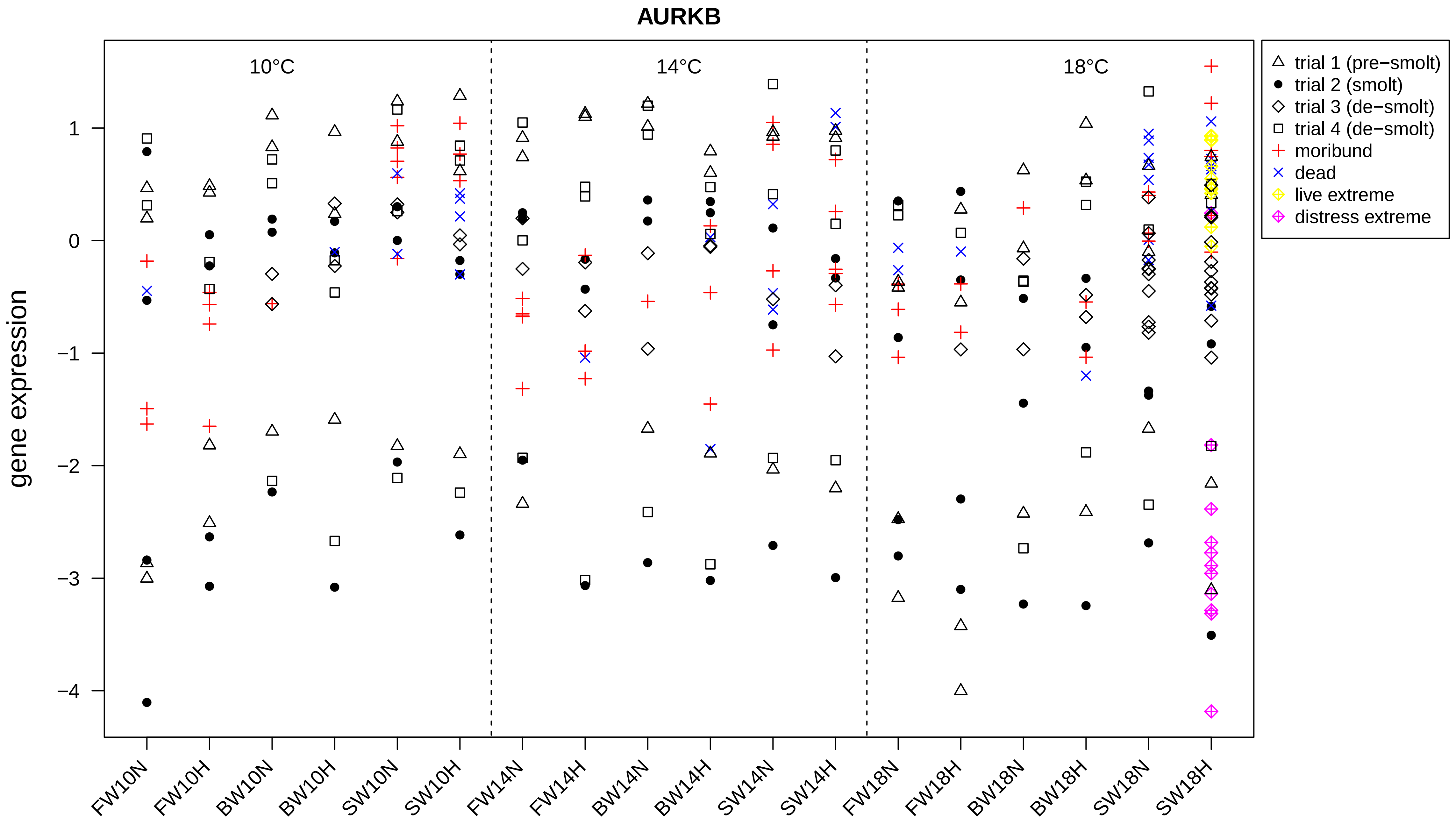
AURKB
10°C
14°C
18°C
gene expression
1
0
−1
−2
−3
−4
FW10N
FW10H
BW10N
BW10H
SW10N
SW10H
FW14N
FW14H
BW14N
BW14H
SW14N
SW14H
FW18N
FW18H
BW18N
BW18H
SW18N
SW18H
trial 1 (pre−smolt)
trial 2 (smolt)
trial 3 (de−smolt)
trial 4 (de−smolt)
moribund
dead
live extreme
distress extreme

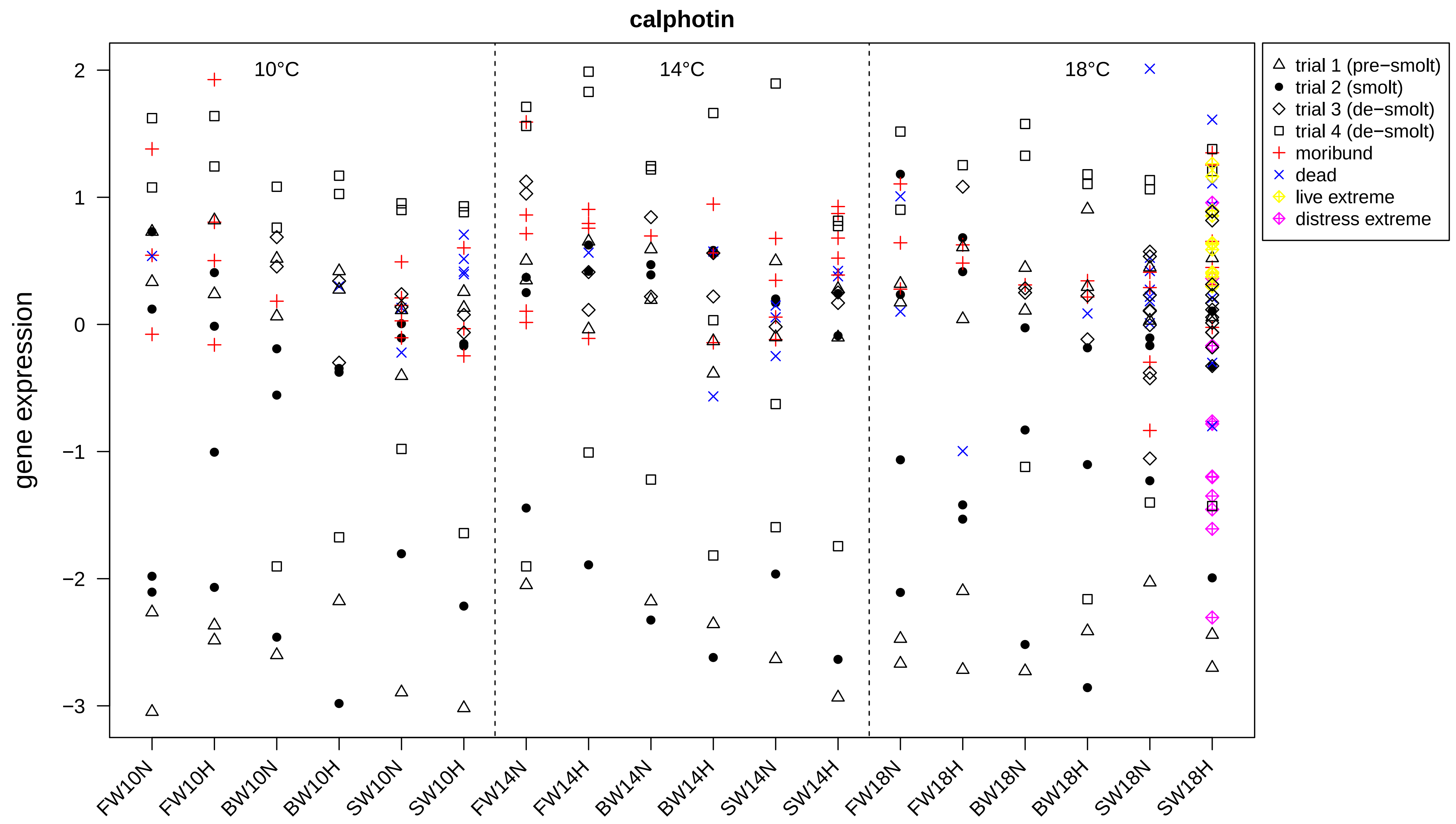
calphotin
gene expression
10°C
14°C
18°C
trial 1 (pre−smolt)
trial 2 (smolt)
trial 3 (de−smolt)
trial 4 (de−smolt)
moribund
dead
live extreme
distress extreme
2
1
0
−1
−2
−3
FW10N
FW10H
BW10N
BW10H
SW10N
SW10H
FW14N
FW14H
BW14N
BW14H
SW14N
SW14H
FW18N
FW18H
BW18N
BW18H
SW18N
SW18H

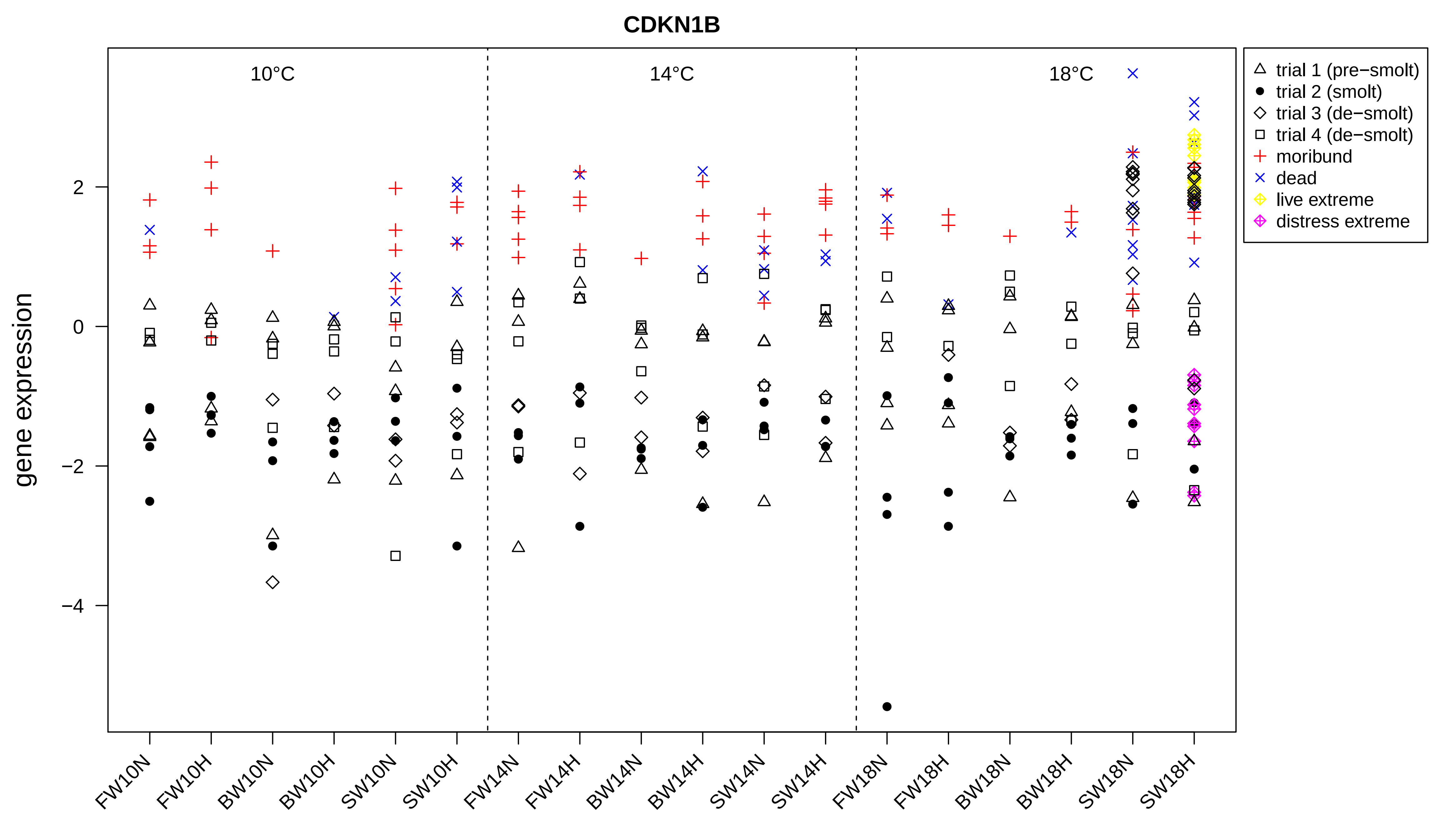
CDKN1B
10°C
14°C
18°C
gene expression
2
0
−2
−4
FW10N
FW10H
BW10N
BW10H
SW10N
SW10H
FW14N
FW14H
BW14N
BW14H
SW14N
SW14H
FW18N
FW18H
BW18N
BW18H
SW18N
SW18H
trial 1 (pre−smolt)
trial 2 (smolt)
trial 3 (de−smolt)
trial 4 (de−smolt)
moribund
dead
live extreme
distress extreme

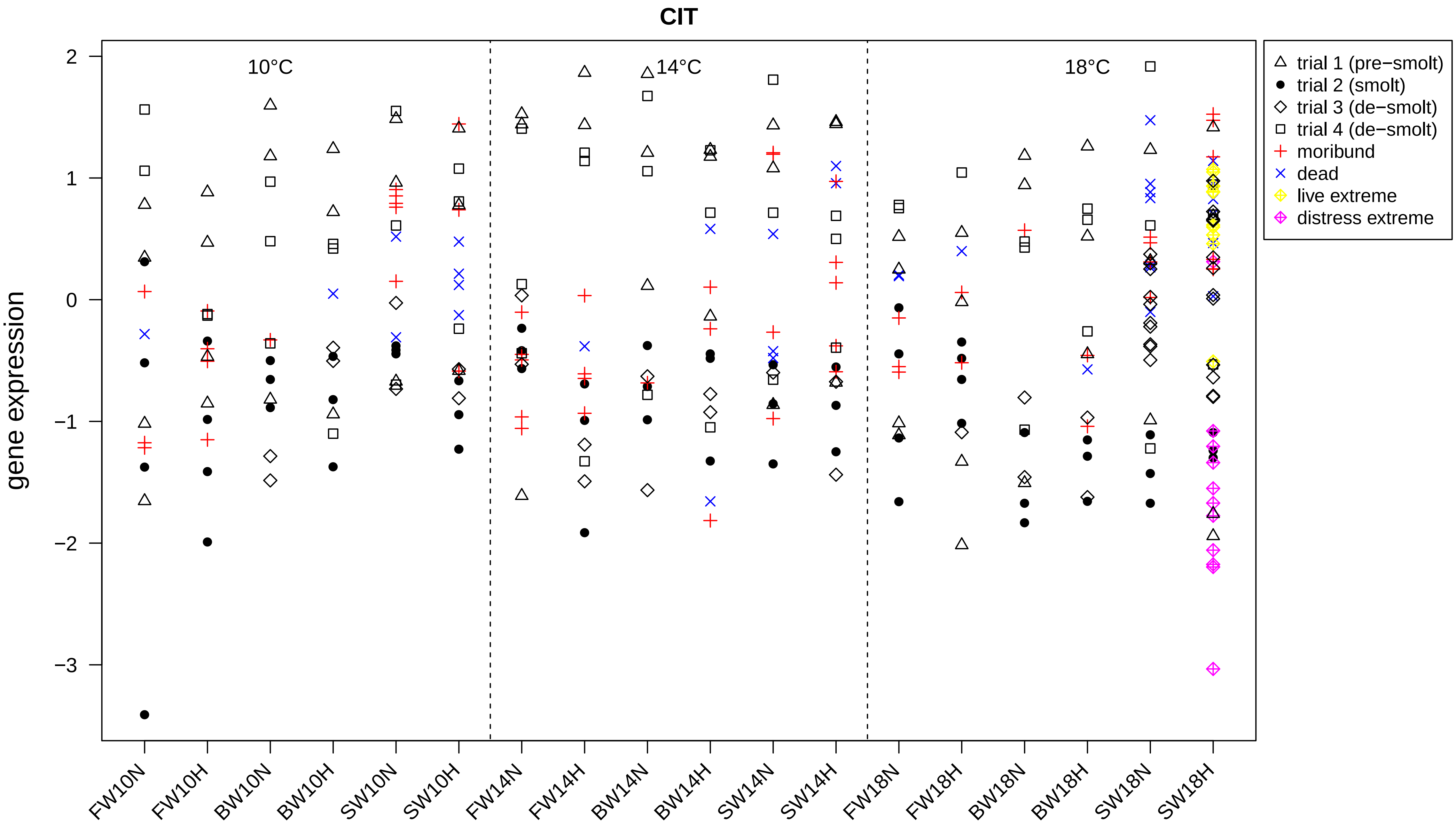

CIT
gene expression
10°C
14°C
18°C
trial 1 (pre−smolt)
trial 2 (smolt)
trial 3 (de−smolt)
trial 4 (de−smolt)
moribund
dead
live extreme
distress extreme
2
1
0
−1
−2
−3
FW10N
FW10H
BW10N
BW10H
SW10N
SW10H
FW14N
FW14H
BW14N
BW14H
SW14N
SW14H
FW18N
FW18H
BW18N
BW18H
SW18N
SW18H

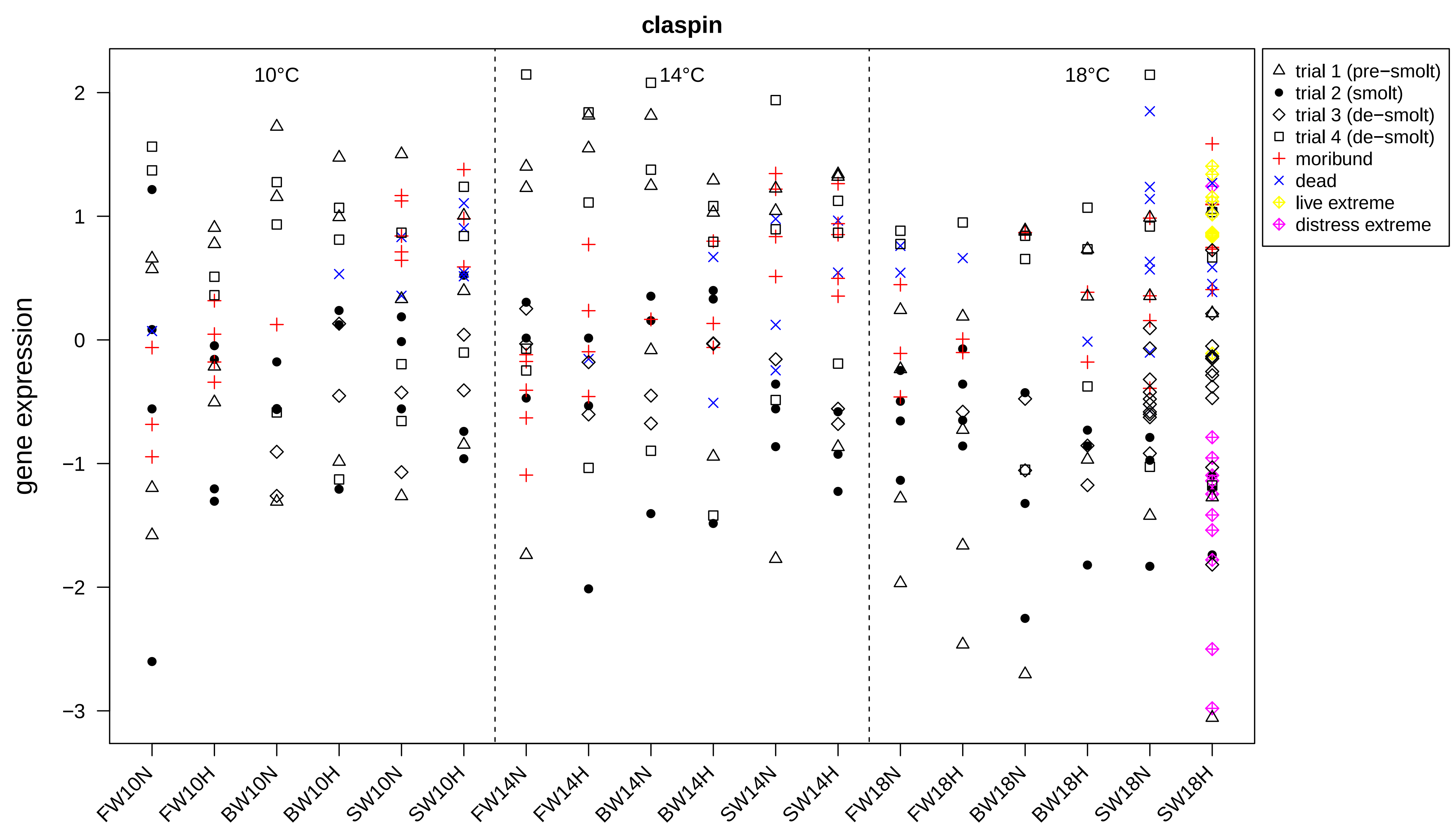
claspin
10°C
14°C
18°C
gene expression
2
1
0
−1
−2
−3
FW10N
FW10H
BW10N
BW10H
SW10N
SW10H
FW14N
FW14H
BW14N
BW14H
SW14N
SW14H
FW18N
FW18H
BW18N
BW18H
SW18N
SW18H
trial 1 (pre−smolt)
trial 2 (smolt)
trial 3 (de−smolt)
trial 4 (de−smolt)
moribund
dead
live extreme
distress extreme

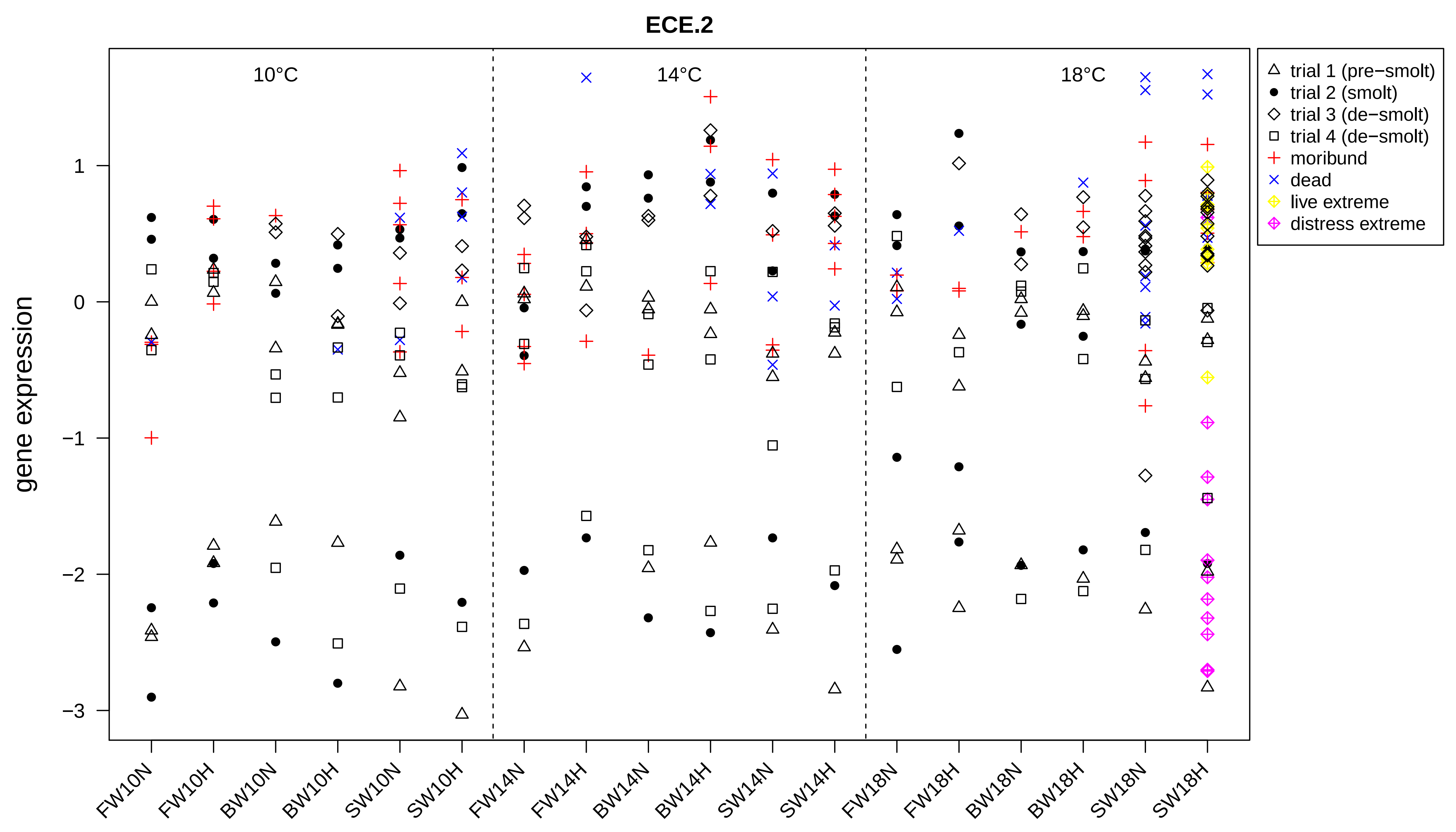
ECE.2
10°C
14°C
18°C
gene expression
1
0
−1
−2
−3
trial 1 (pre−smolt)
trial 2 (smolt)
trial 3 (de−smolt)
trial 4 (de−smolt)
moribund
dead
live extreme
distress extreme
FW10N
FW10H
BW10N
BW10H
SW10N
SW10H
FW14N
FW14H
BW14N
BW14H
SW14N
SW14H
FW18N
FW18H
BW18N
BW18H
SW18N
SW18H

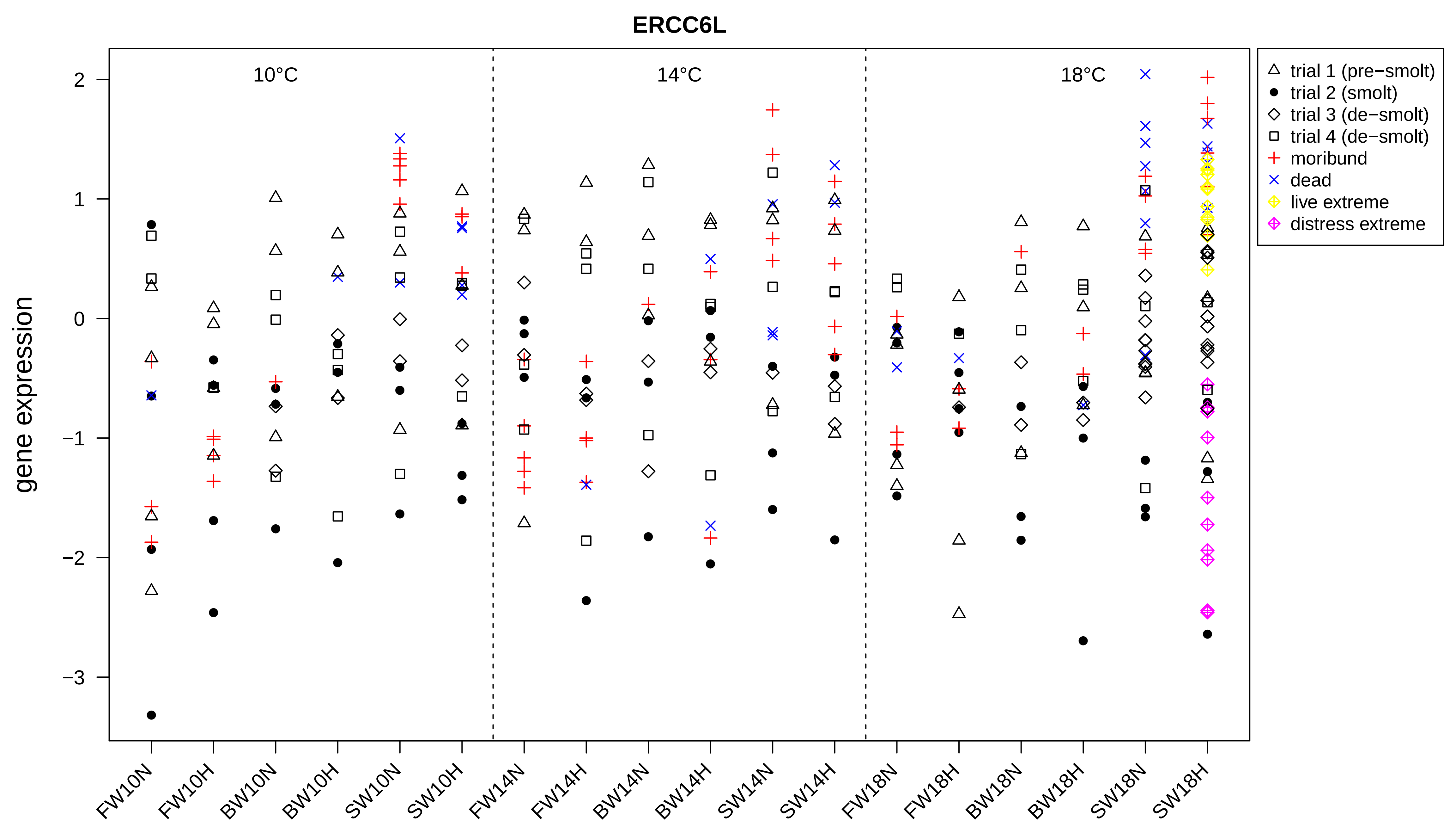
ERCC6L
10°C
14°C
18°C
gene expression
2
1
0
−1
−2
−3
FW10N
FW10H
BW10N
BW10H
SW10N
SW10H
FW14N
FW14H
BW14N
BW14H
SW14N
SW14H
FW18N
FW18H
BW18N
BW18H
SW18N
SW18H
trial 1 (pre−smolt)
trial 2 (smolt)
trial 3 (de−smolt)
trial 4 (de−smolt)
moribund
dead
live extreme
distress extreme

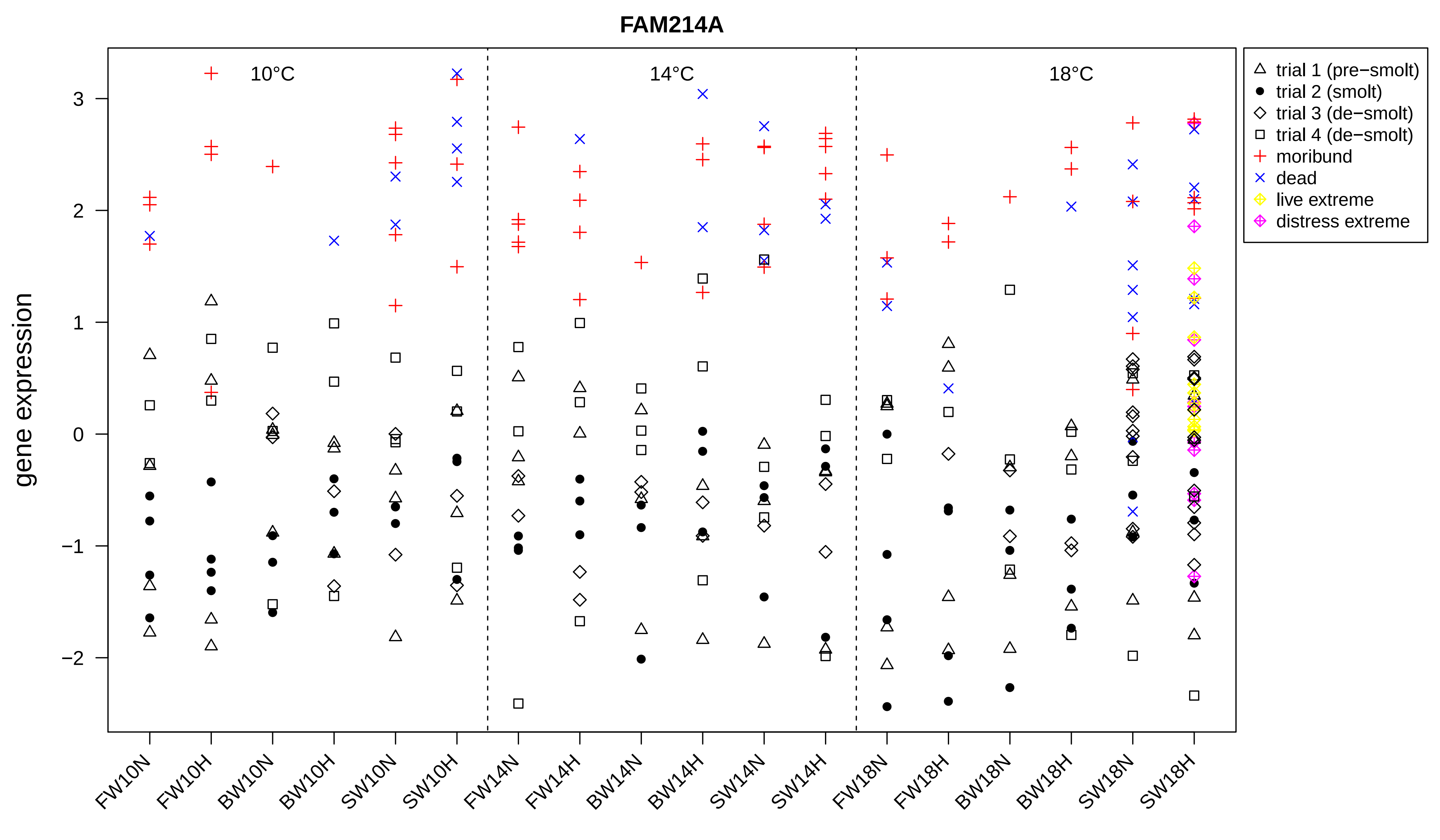
FAM214A
gene expression
10°C
14°C
18°C
trial 1 (pre−smolt)
trial 2 (smolt)
trial 3 (de−smolt)
trial 4 (de−smolt)
moribund
dead
live extreme
distress extreme
3
2
1
0
−1
−2
FW10N
FW10H
BW10N
BW10H
SW10N
SW10H
FW14N
FW14H
BW14N
BW14H
SW14N
SW14H
FW18N
FW18H
BW18N
BW18H
SW18N
SW18H

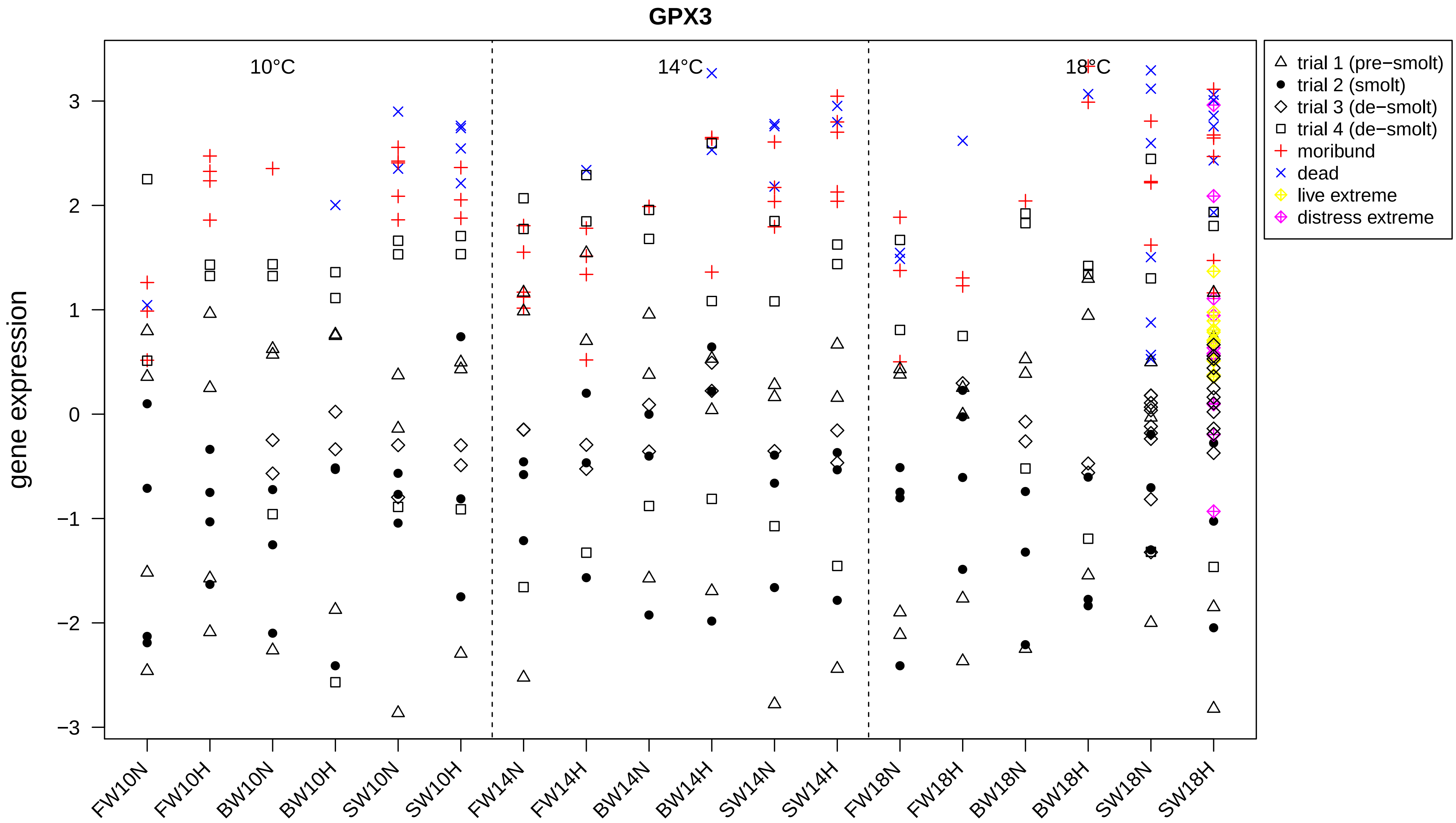
GPX3
gene expression
10°C
14°C
18°C
3
2
1
0
−1
−2
−3
FW10N
FW10H
BW10N
BW10H
SW10N
SW10H
FW14N
FW14H
BW14N
BW14H
SW14N
SW14H
FW18N
FW18H
BW18N
BW18H
SW18N
SW18H
trial 1 (pre−smolt)
trial 2 (smolt)
trial 3 (de−smolt)
trial 4 (de−smolt)
moribund
dead
live extreme
distress extreme

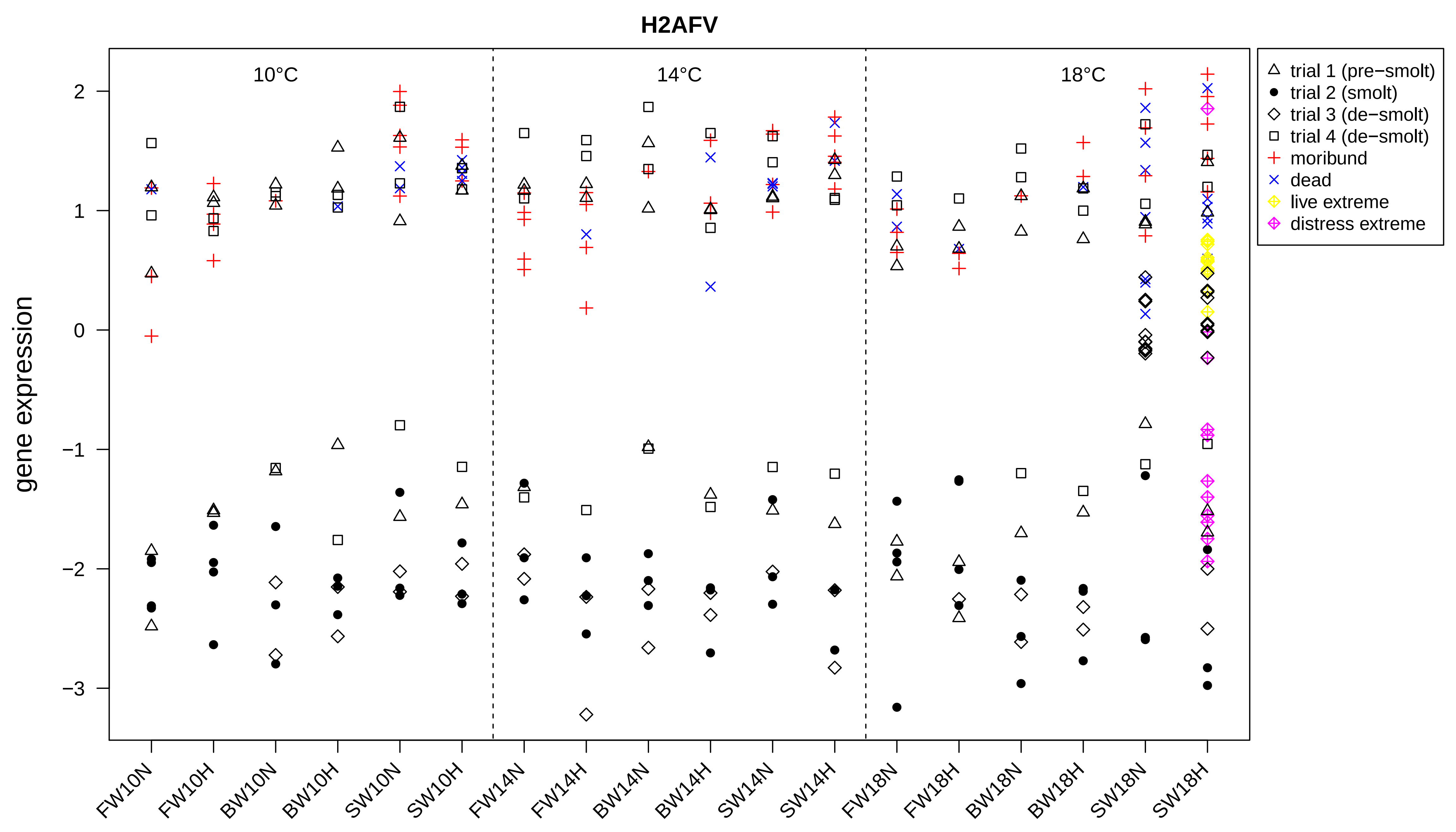

H2AFV
10°C
14°C
18°C
gene expression
2
1
0
−1
−2
−3
FW10N
FW10H
BW10N
BW10H
SW10N
SW10H
FW14N
FW14H
BW14N
BW14H
SW14N
SW14H
FW18N
FW18H
BW18N
BW18H
SW18N
SW18H
trial 1 (pre−smolt)
trial 2 (smolt)
trial 3 (de−smolt)
trial 4 (de−smolt)
moribund
dead
live extreme
distress extreme

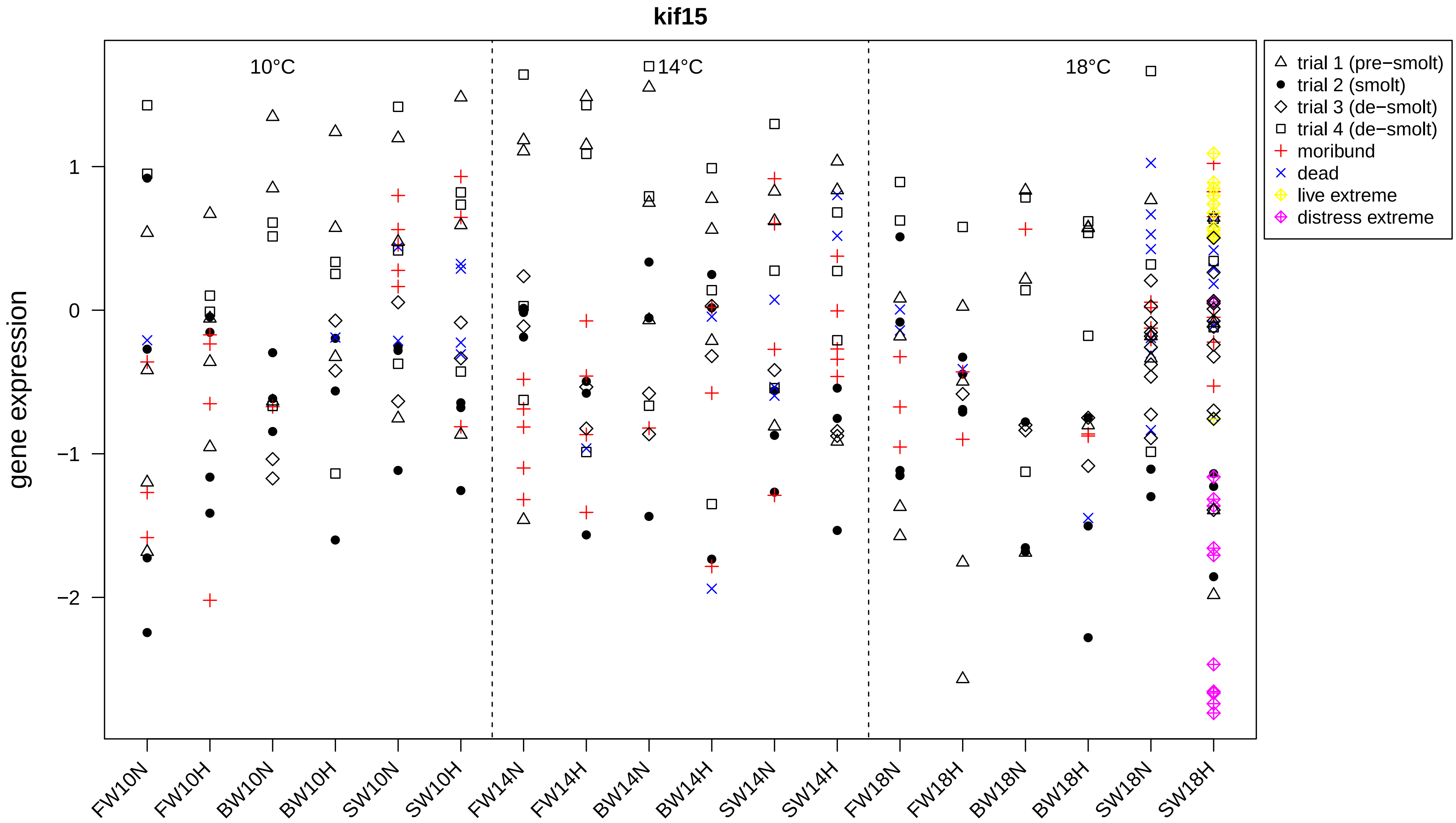

kif15
10°C
14°C
18°C
gene expression
1
0
−1
−2
FW10N
FW10H
BW10N
BW10H
SW10N
SW10H
FW14N
FW14H
BW14N
BW14H
SW14N
SW14H
FW18N
FW18H
BW18N
BW18H
SW18N
SW18H
trial 1 (pre−smolt)
trial 2 (smolt)
trial 3 (de−smolt)
trial 4 (de−smolt)
moribund
dead
live extreme
distress extreme

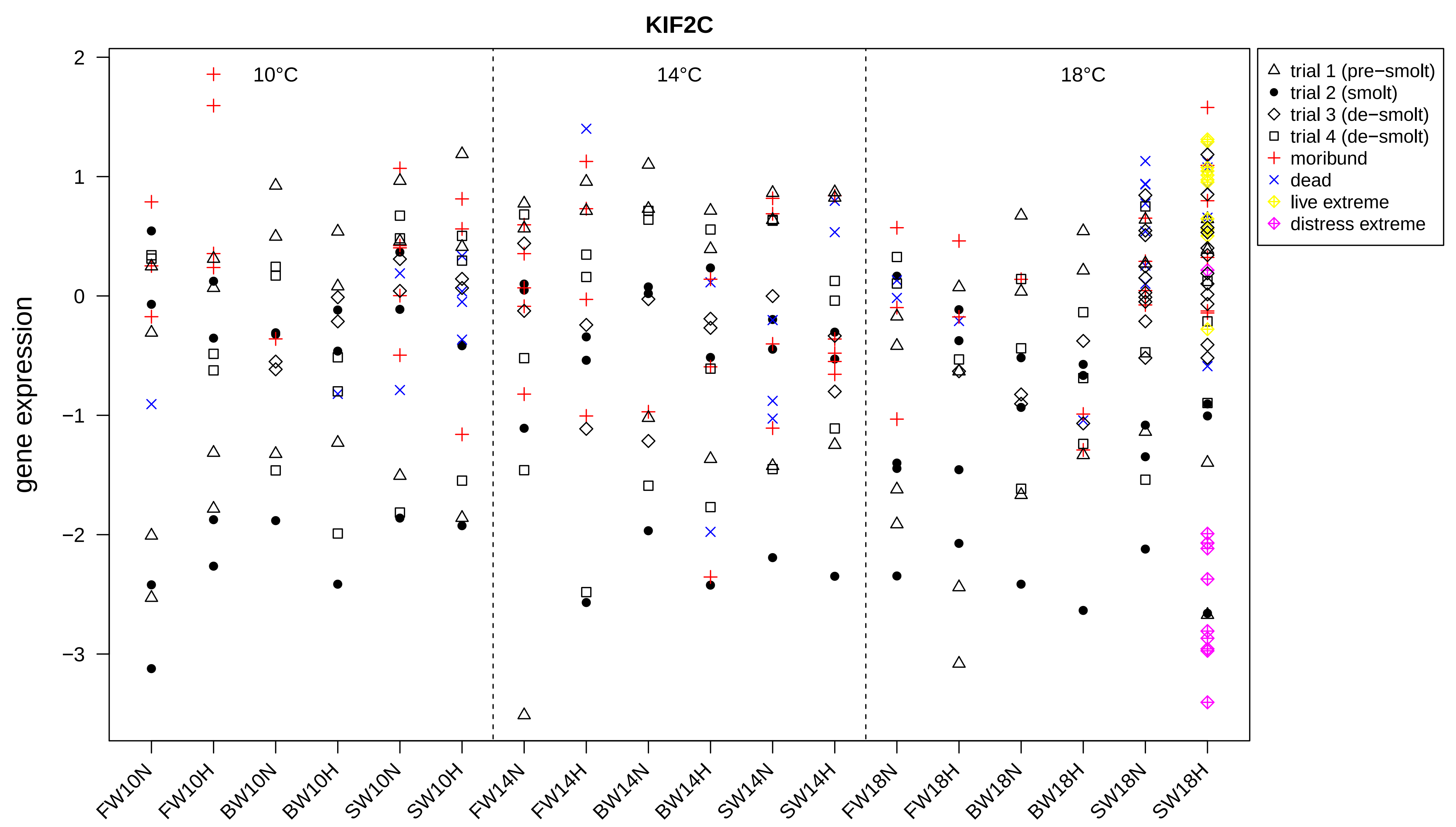
KIF2C
10°C
14°C
18°C
gene expression
2
1
0
−1
−2
−3
FW10N
FW10H
BW10N
BW10H
SW10N
SW10H
FW14N
FW14H
BW14N
BW14H
SW14N
SW14H
FW18N
FW18H
BW18N
BW18H
SW18N
SW18H
trial 1 (pre−smolt)
trial 2 (smolt)
trial 3 (de−smolt)
trial 4 (de−smolt)
moribund
dead
live extreme
distress extreme

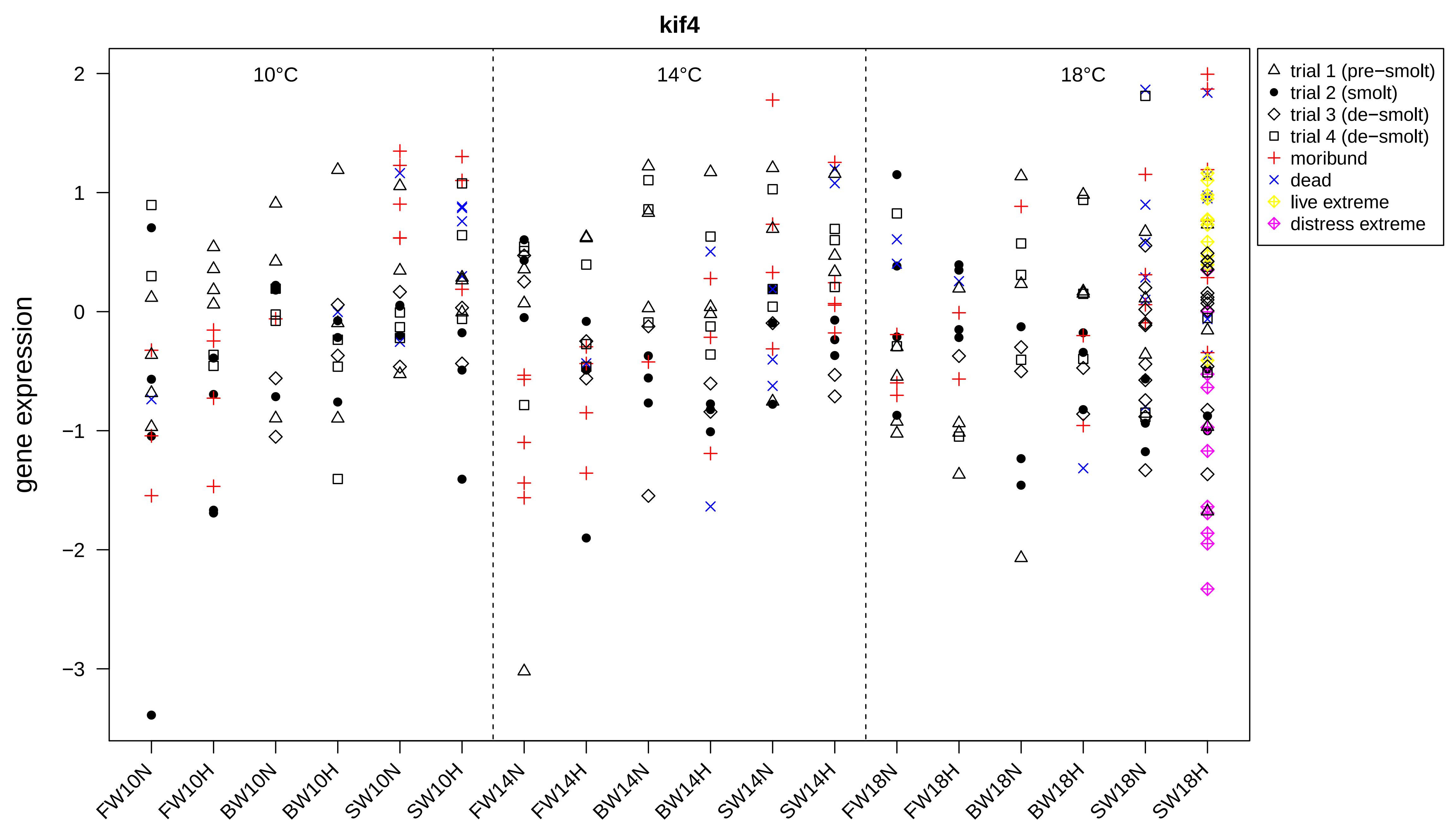

kif4
10°C
14°C
18°C
gene expression
2
1
0
−1
−2
−3
FW10N
FW10H
BW10N
BW10H
SW10N
SW10H
FW14N
FW14H
BW14N
BW14H
SW14N
SW14H
FW18N
FW18H
BW18N
BW18H
SW18N
SW18H
trial 1 (pre−smolt)
trial 2 (smolt)
trial 3 (de−smolt)
trial 4 (de−smolt)
moribund
dead
live extreme
distress extreme

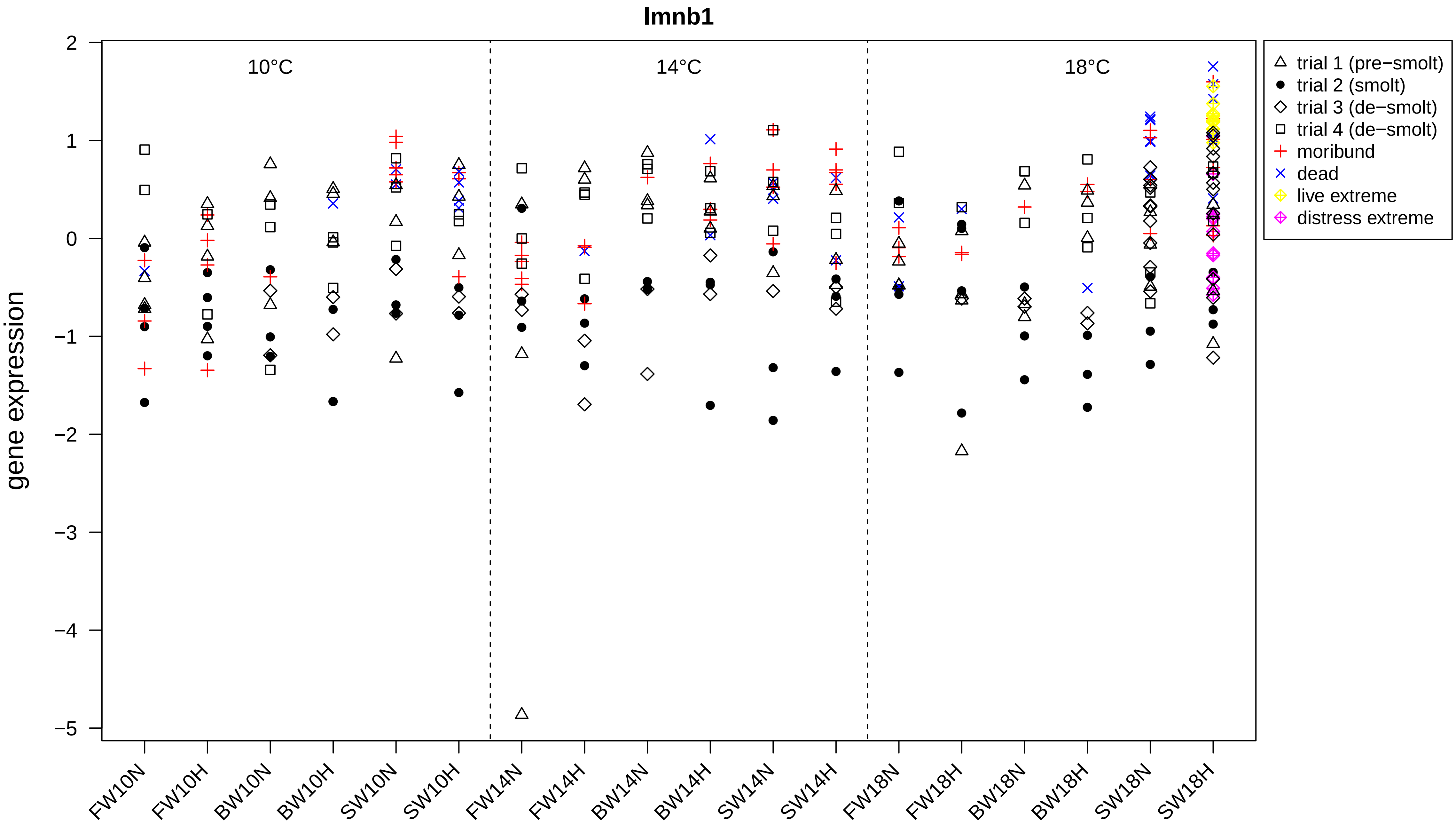

lmnb1
gene expression
10°C
14°C
18°C
2
1
0
−1
−2
−3
−4
−5
FW10N
FW10H
BW10N
BW10H
SW10N
SW10H
FW14N
FW14H
BW14N
BW14H
SW14N
SW14H
FW18N
FW18H
BW18N
BW18H
SW18N
SW18H
trial 1 (pre−smolt)
trial 2 (smolt)
trial 3 (de−smolt)
trial 4 (de−smolt)
moribund
dead
live extreme
distress extreme

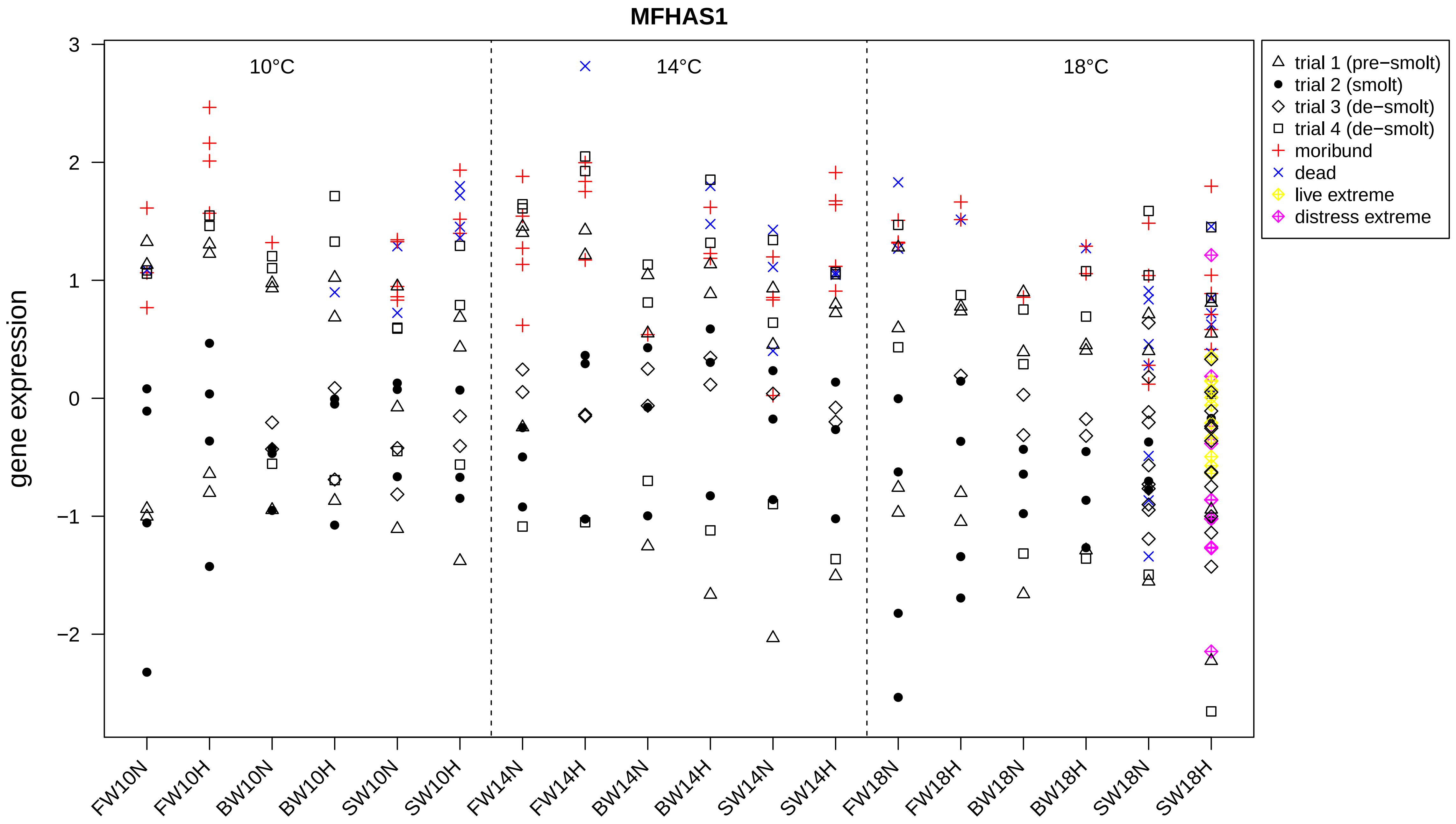
MFHAS1
gene expression
3
2
1
0
−1
−2
10°C
14°C
18°C
FW10N
FW10H
BW10N
BW10H
SW10N
SW10H
FW14N
FW14H
BW14N
BW14H
SW14N
SW14H
FW18N
FW18H
BW18N
BW18H
SW18N
SW18H
trial 1 (pre−smolt)
trial 2 (smolt)
trial 3 (de−smolt)
trial 4 (de−smolt)
moribund
dead
live extreme
distress extreme

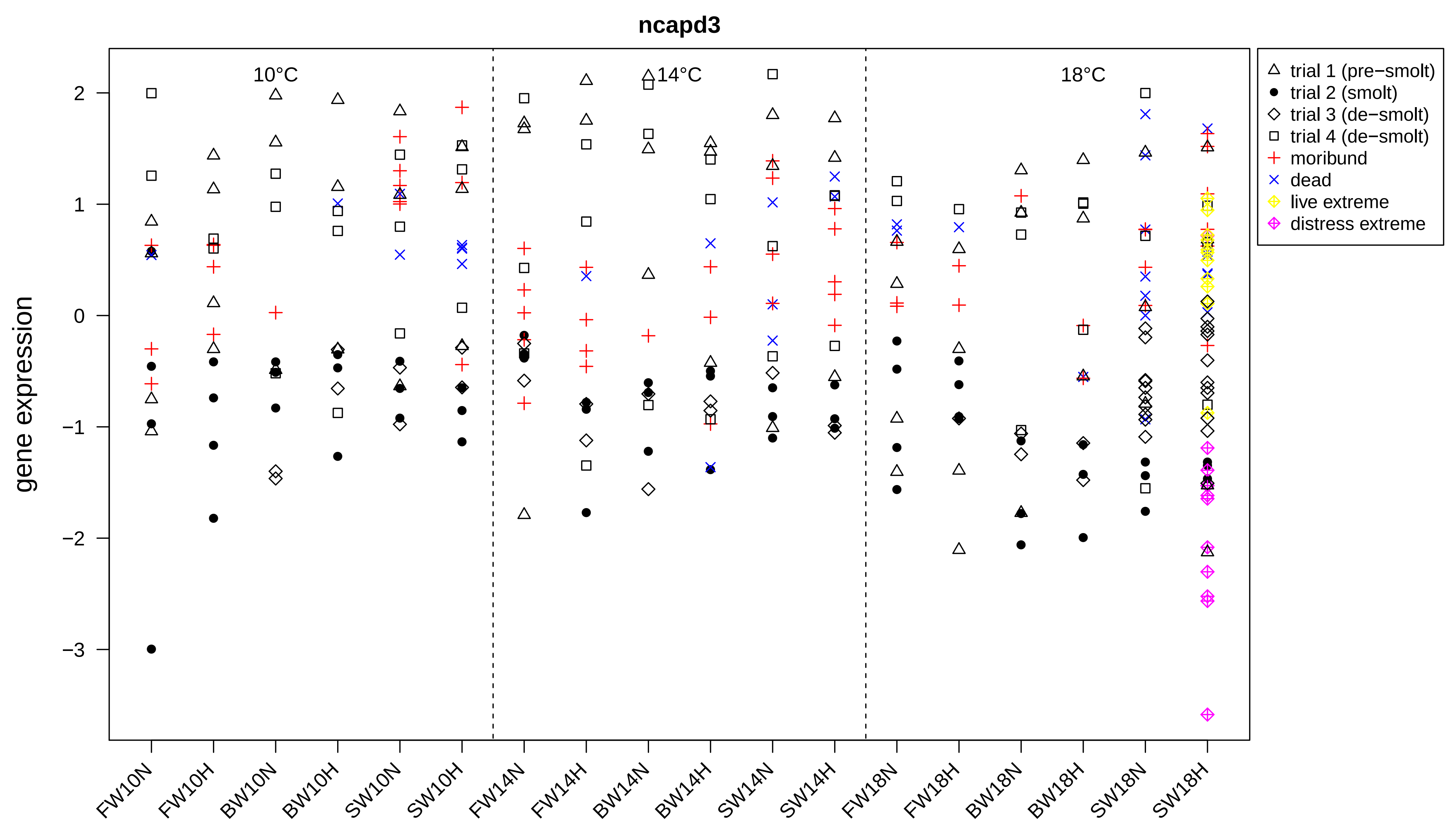

ncapd3
10°C
14°C
18°C
gene expression
2
1
0
−1
−2
−3
FW10N
FW10H
BW10N
BW10H
SW10N
SW10H
FW14N
FW14H
BW14N
BW14H
SW14N
SW14H
FW18N
FW18H
BW18N
BW18H
SW18N
SW18H
trial 1 (pre−smolt)
trial 2 (smolt)
trial 3 (de−smolt)
trial 4 (de−smolt)
moribund
dead
live extreme
distress extreme

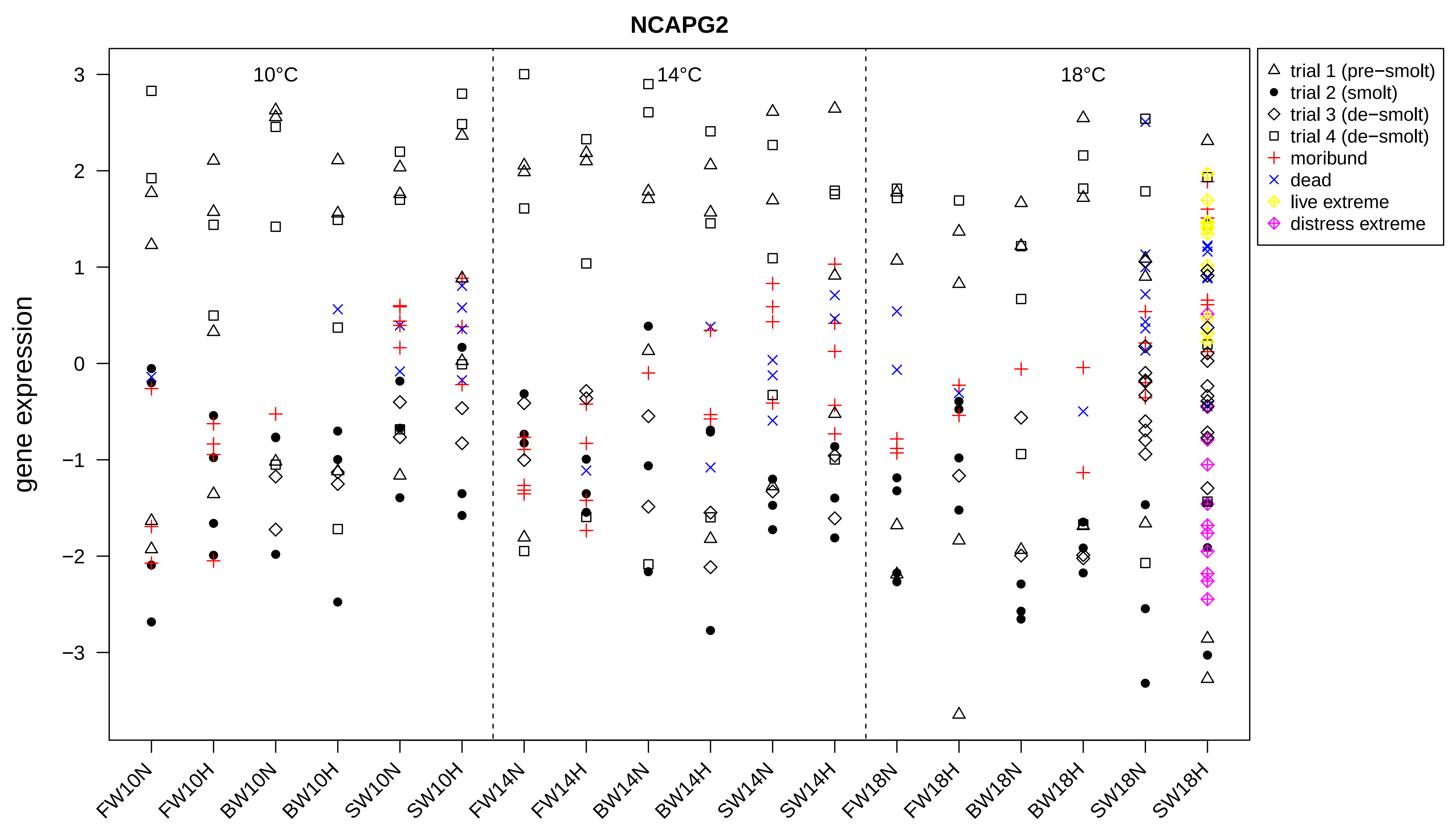

NCAPG2
10°C
14°C
18°C
gene expression
3
2
1
0
−1
−2
−3
FW10N
FW10H
BW10N
BW10H
SW10N
SW10H
FW14N
FW14H
BW14N
BW14H
SW14N
SW14H
FW18N
FW18H
BW18N
BW18H
SW18N
SW18H
trial 1 (pre−smolt)
trial 2 (smolt)
trial 3 (de−smolt)
trial 4 (de−smolt)
moribund
dead
live extreme
distress extreme

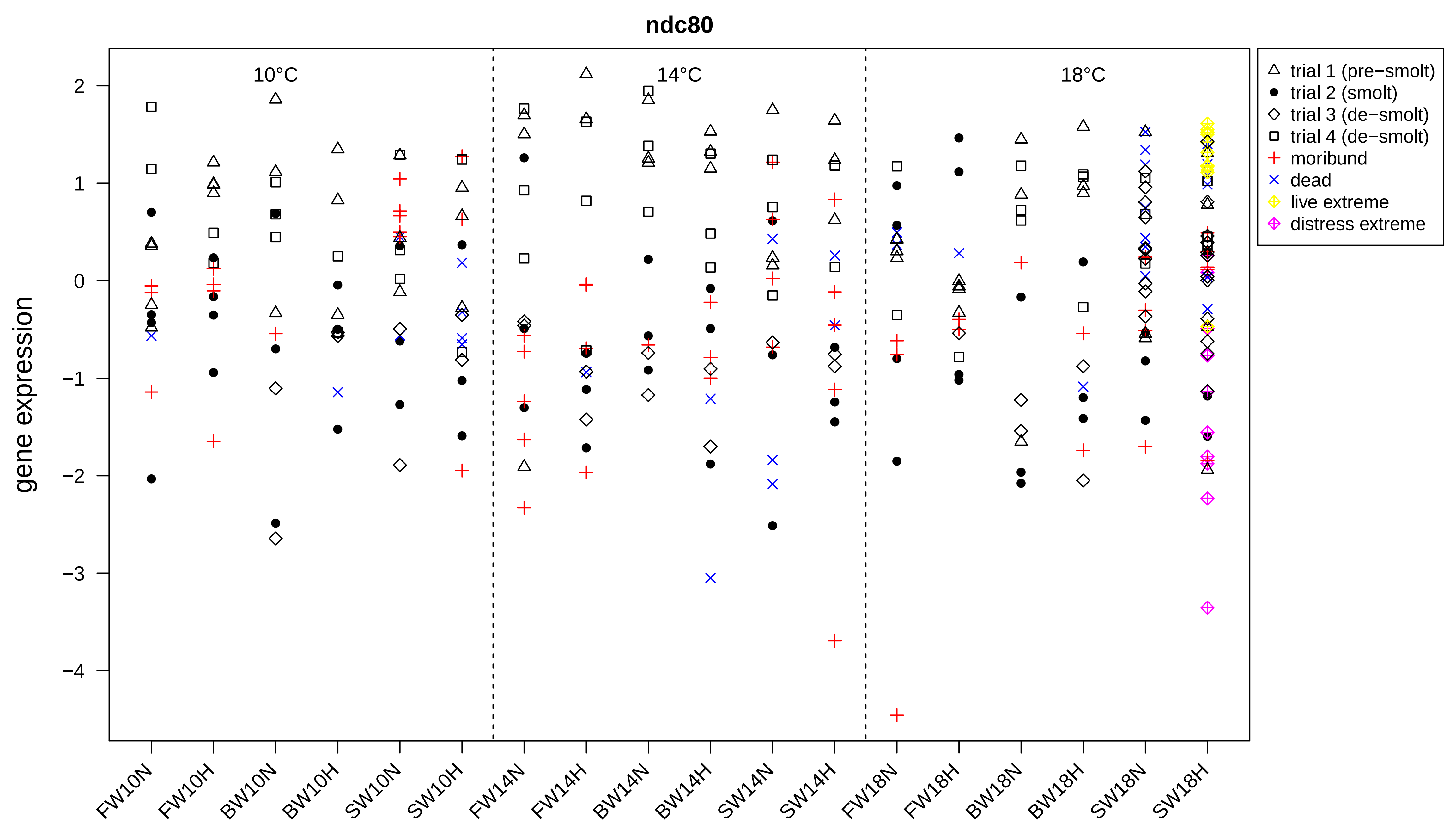
ndc80
10°C
14°C
18°C
gene expression
2
1
0
−1
−2
−3
−4
FW10N
FW10H
BW10N
BW10H
SW10N
SW10H
FW14N
FW14H
BW14N
BW14H
SW14N
SW14H
FW18N
FW18H
BW18N
BW18H
SW18N
SW18H
trial 1 (pre−smolt)
trial 2 (smolt)
trial 3 (de−smolt)
trial 4 (de−smolt)
moribund
dead
live extreme
distress extreme

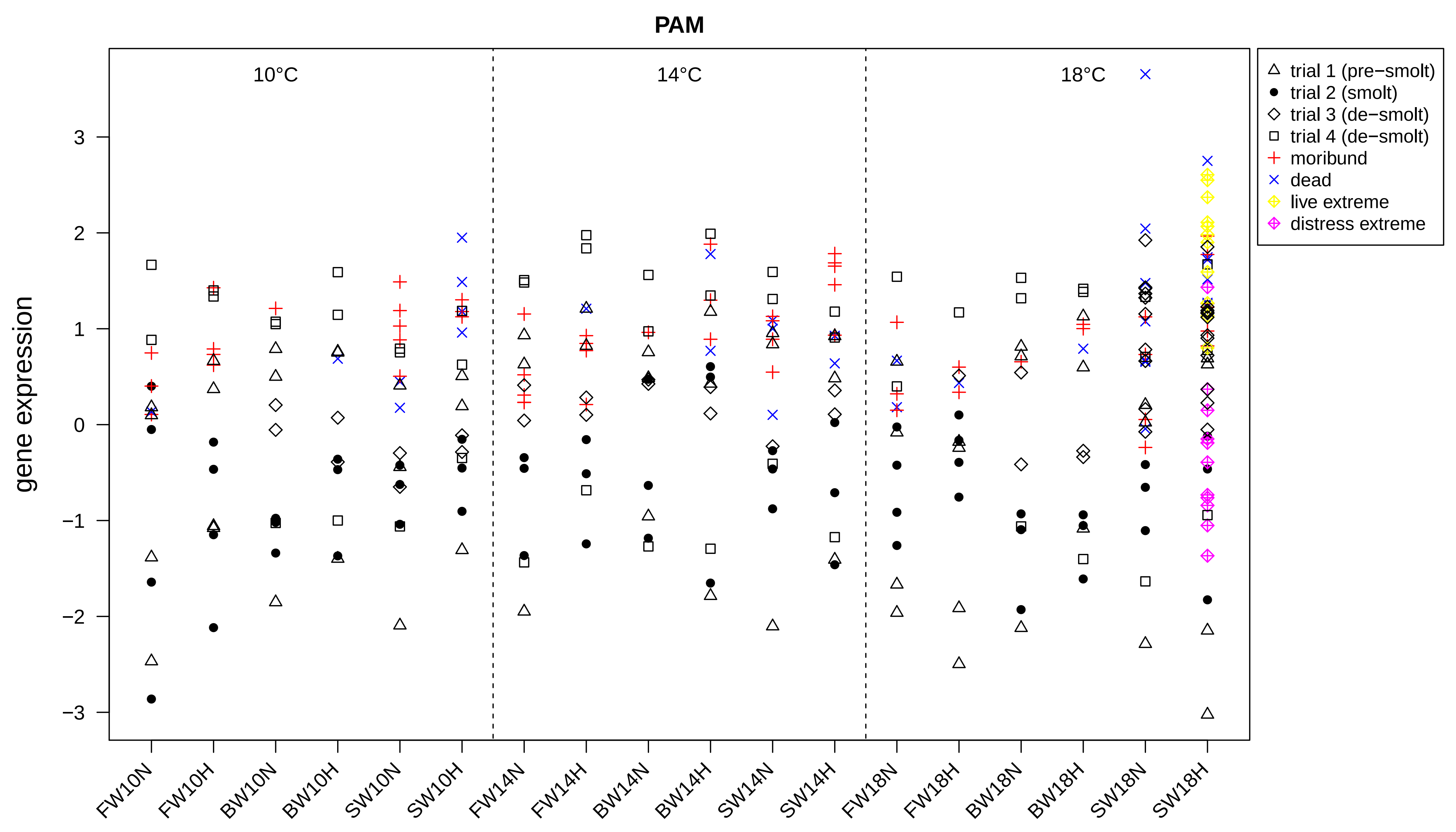
PAM
10°C
14°C
18°C
gene expression
3
2
1
0
−1
−2
−3
FW10N
FW10H
BW10N
BW10H
SW10N
SW10H
FW14N
FW14H
BW14N
BW14H
SW14N
SW14H
FW18N
FW18H
BW18N
BW18H
SW18N
SW18H
trial 1 (pre−smolt)
trial 2 (smolt)
trial 3 (de−smolt)
trial 4 (de−smolt)
moribund
dead
live extreme
distress extreme

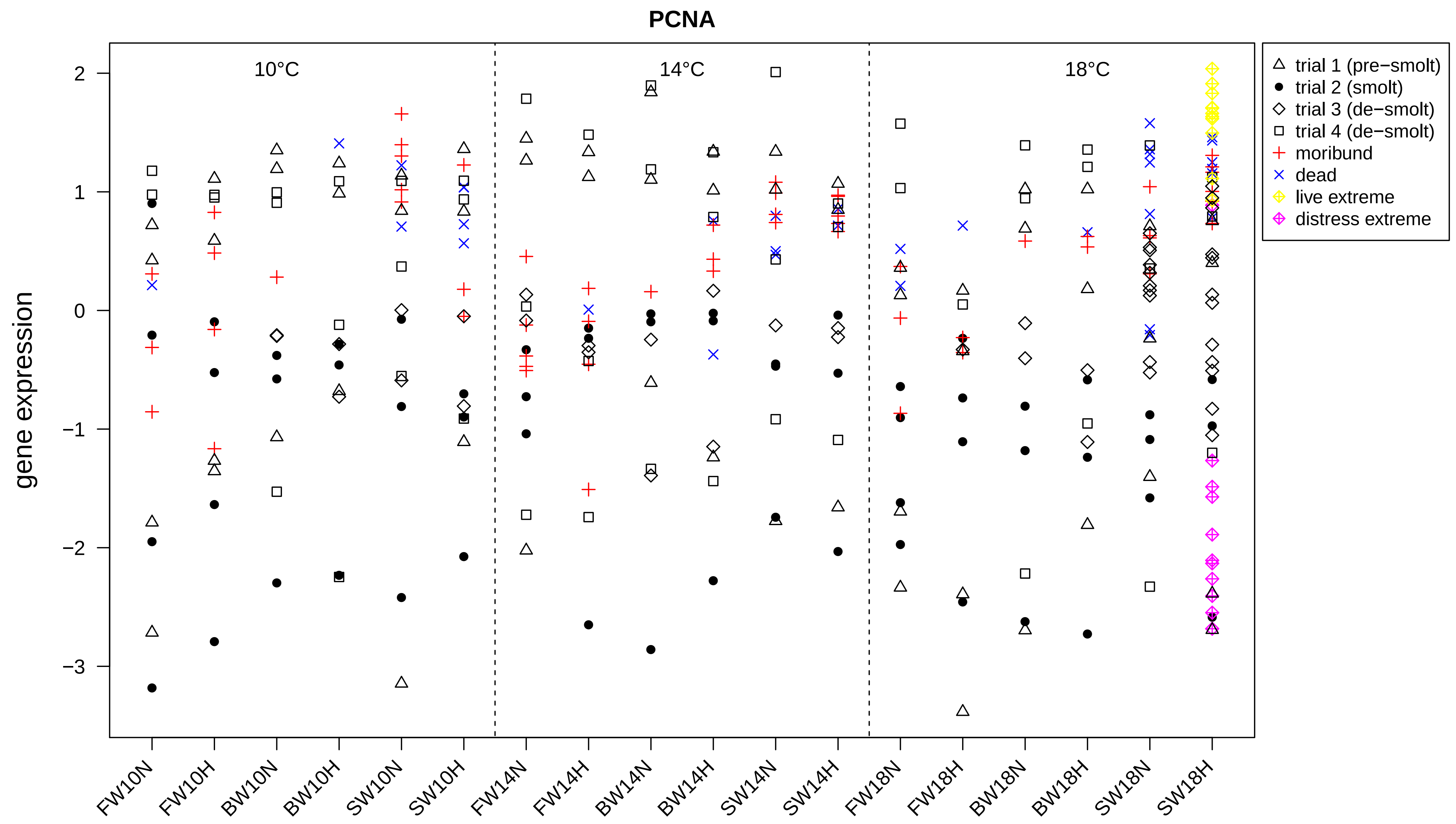
PCNA
10°C
14°C
18°C
gene expression
2
1
0
−1
−2
−3
FW10N
FW10H
BW10N
BW10H
SW10N
SW10H
FW14N
FW14H
BW14N
BW14H
SW14N
SW14H
FW18N
FW18H
BW18N
BW18H
SW18N
SW18H
trial 1 (pre−smolt)
trial 2 (smolt)
trial 3 (de−smolt)
trial 4 (de−smolt)
moribund
dead
live extreme
distress extreme

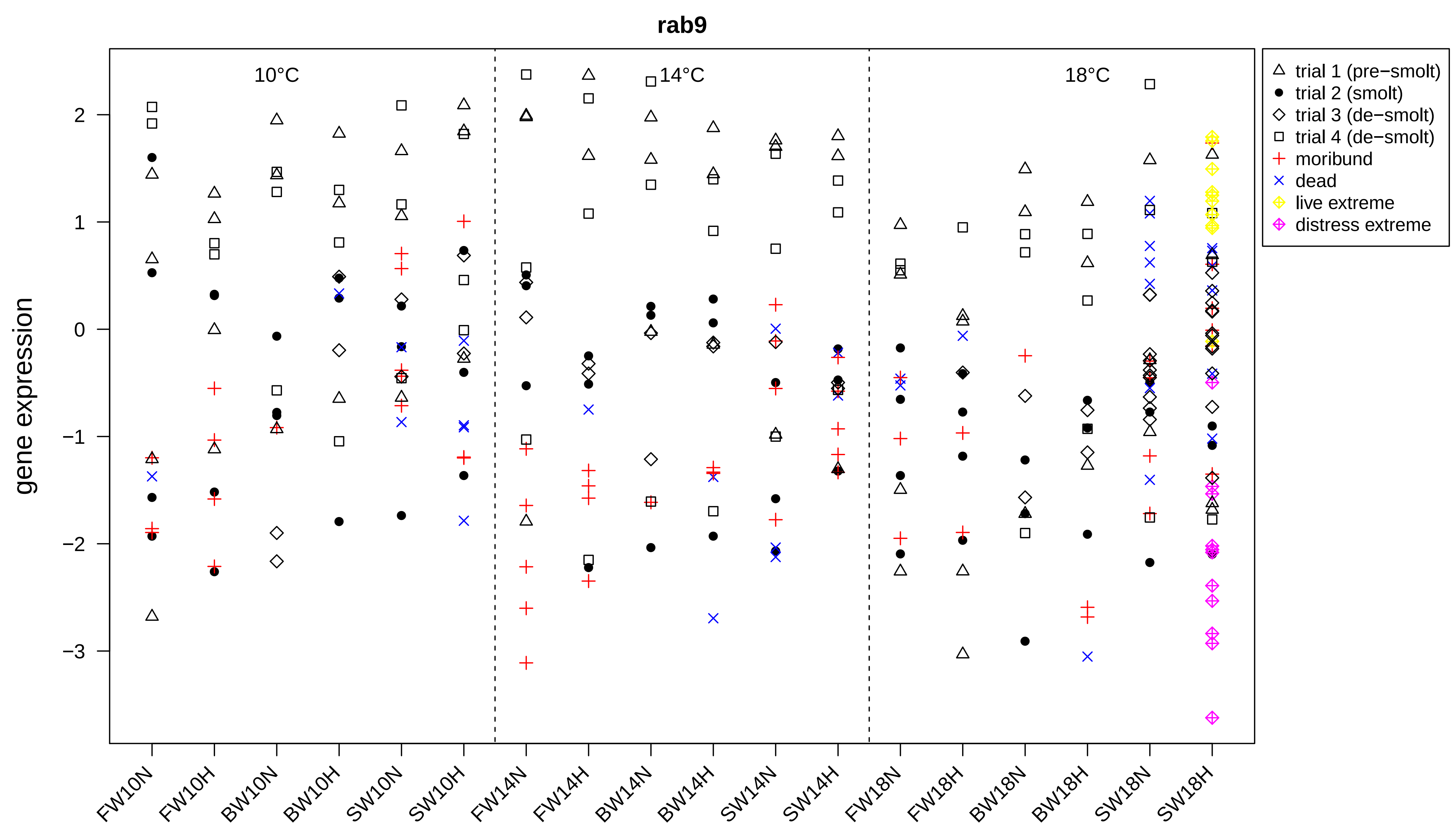
rab9
10°C
14°C
18°C
gene expression
2
1
0
−1
−2
−3
FW10N
FW10H
BW10N
BW10H
SW10N
SW10H
FW14N
FW14H
BW14N
BW14H
SW14N
SW14H
FW18N
FW18H
BW18N
BW18H
SW18N
SW18H
trial 1 (pre−smolt)
trial 2 (smolt)
trial 3 (de−smolt)
trial 4 (de−smolt)
moribund
dead
live extreme
distress extreme

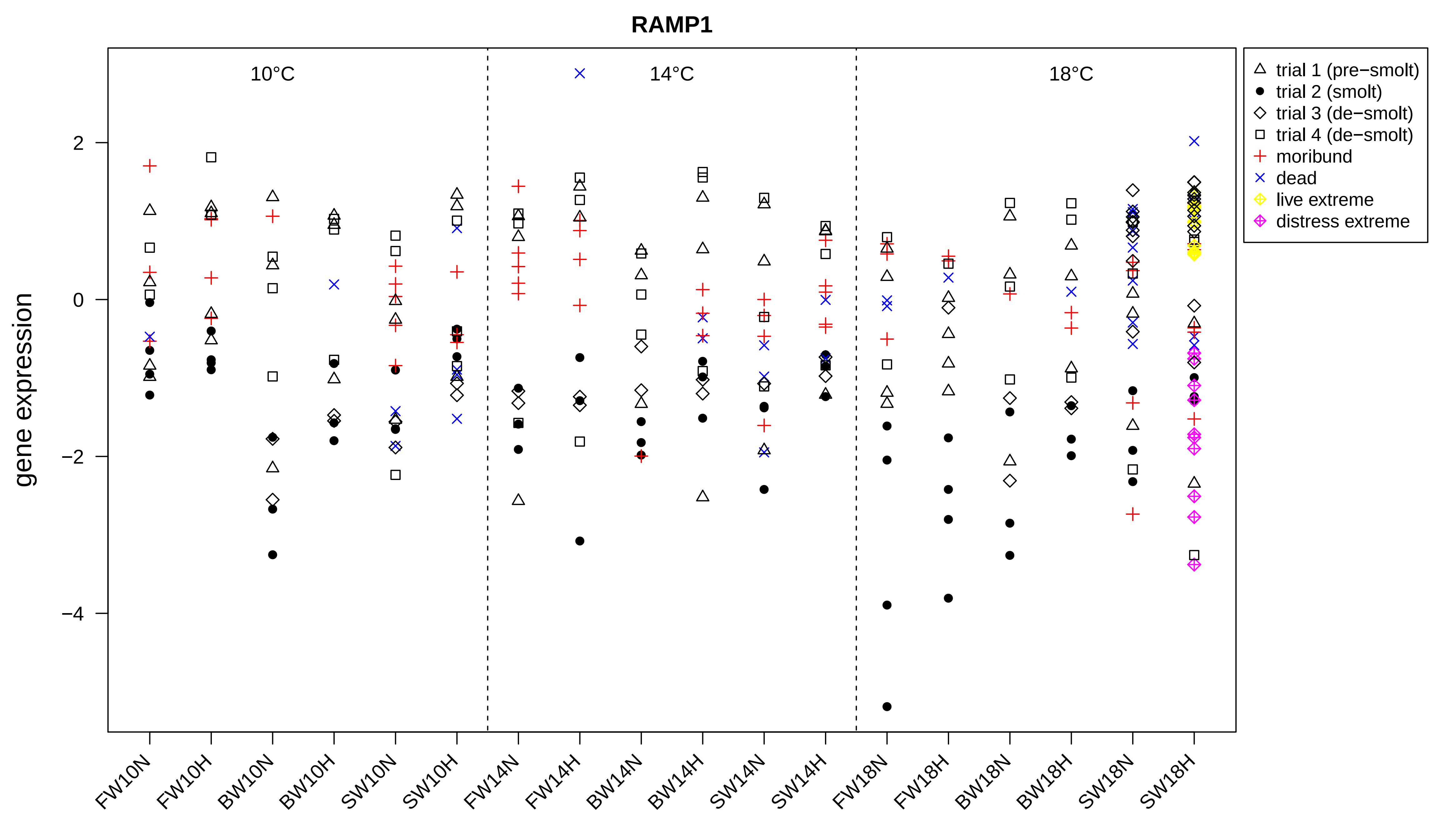
RAMP1
10°C
14°C
18°C
gene expression
2
0
−2
−4
trial 1 (pre−smolt)
trial 2 (smolt)
trial 3 (de−smolt)
trial 4 (de−smolt)
moribund
dead
live extreme
distress extreme
FW10N
FW10H
BW10N
BW10H
SW10N
SW10H
FW14N
FW14H
BW14N
BW14H
SW14N
SW14H
FW18N
FW18H
BW18N
BW18H
SW18N
SW18H

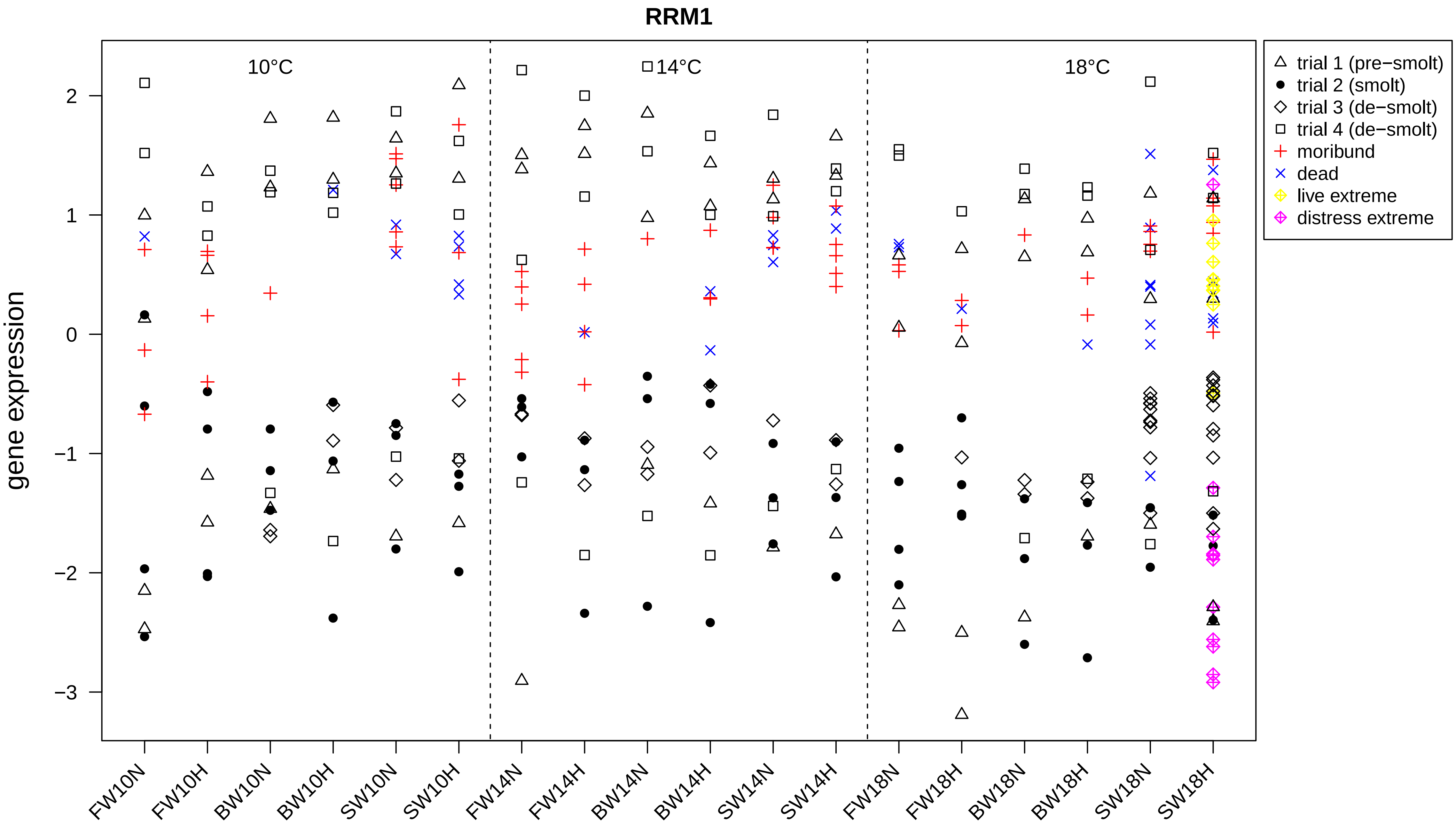

RRM1
gene expression
10°C
14°C
18°C
2
1
0
−1
−2
−3
trial 1 (pre−smolt)
trial 2 (smolt)
trial 3 (de−smolt)
trial 4 (de−smolt)
moribund
dead
live extreme
distress extreme
FW10N
FW10H
BW10N
BW10H
SW10N
SW10H
FW14N
FW14H
BW14N
BW14H
SW14N
SW14H
FW18N
FW18H
BW18N
BW18H
SW18N
SW18H

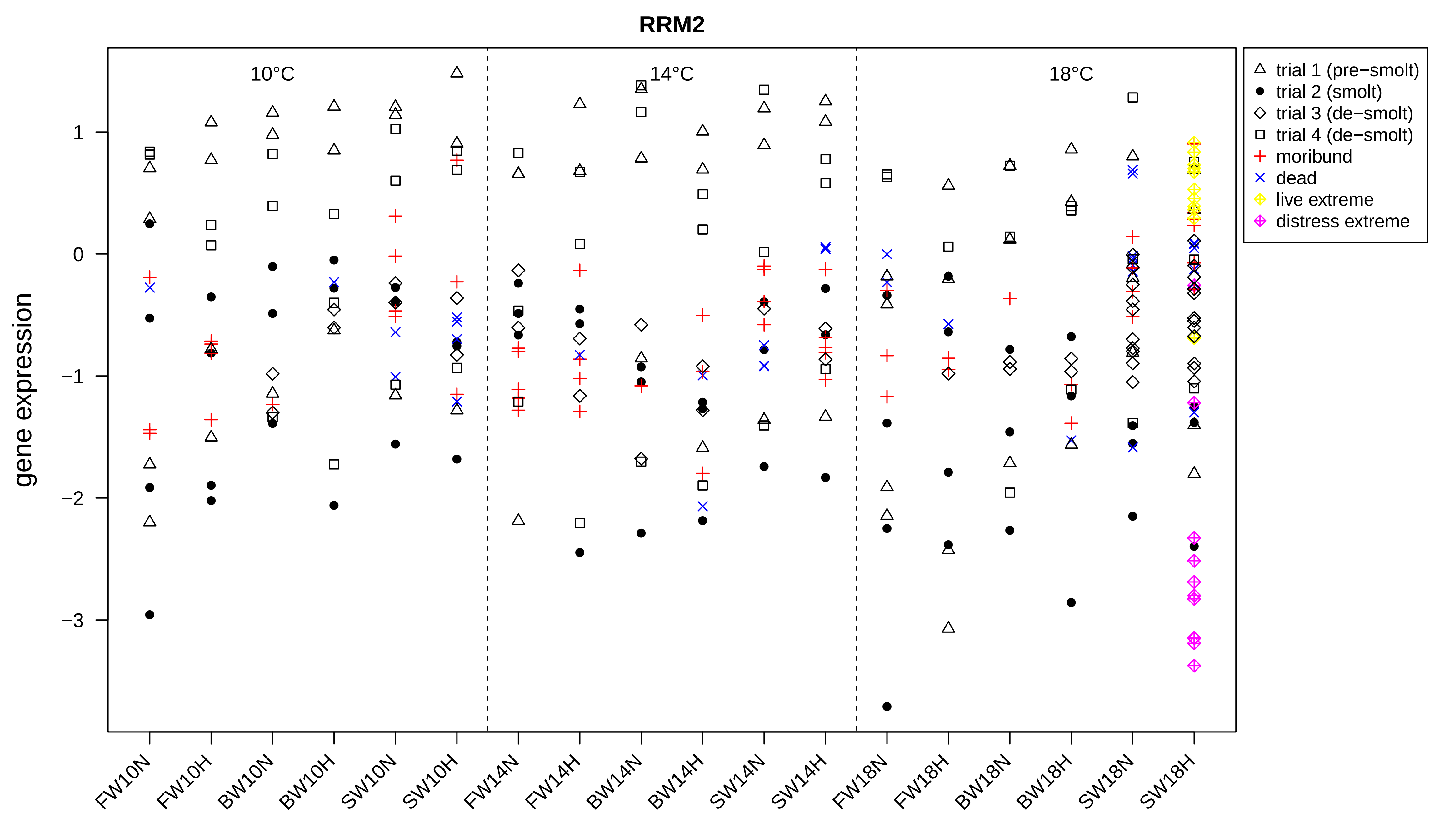

RRM2
10°C
14°C
18°C
gene expression
1
0
−1
−2
−3
FW10N
FW10H
BW10N
BW10H
SW10N
SW10H
FW14N
FW14H
BW14N
BW14H
SW14N
SW14H
FW18N
FW18H
BW18N
BW18H
SW18N
SW18H
trial 1 (pre−smolt)
trial 2 (smolt)
trial 3 (de−smolt)
trial 4 (de−smolt)
moribund
dead
live extreme
distress extreme

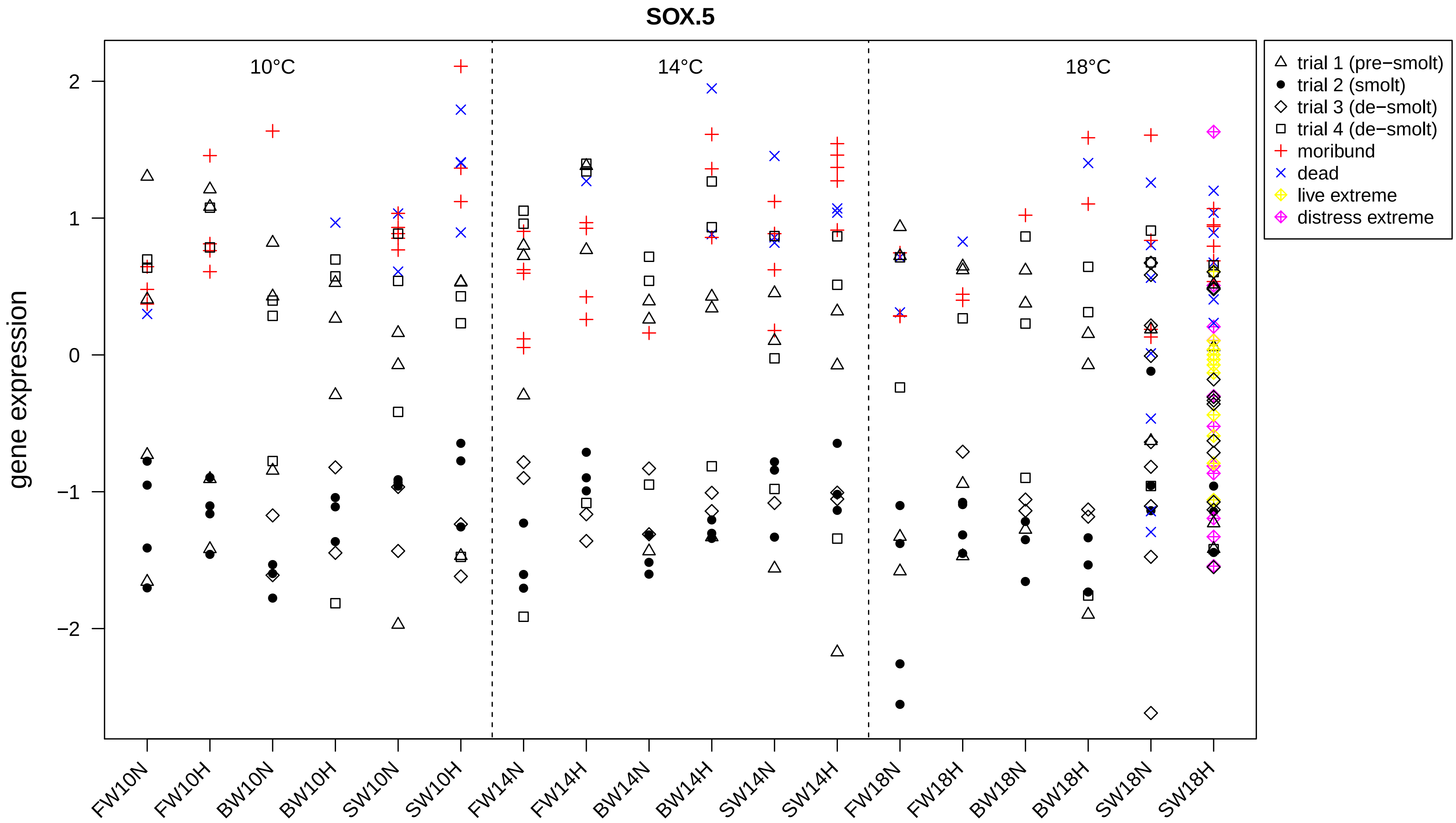
SOX.5
gene expression
10°C
14°C
18°C
2
1
0
−1
−2
FW10N
FW10H
BW10N
BW10H
SW10N
SW10H
FW14N
FW14H
BW14N
BW14H
SW14N
SW14H
FW18N
FW18H
BW18N
BW18H
SW18N
SW18H
trial 1 (pre−smolt)
trial 2 (smolt)
trial 3 (de−smolt)
trial 4 (de−smolt)
moribund
dead
live extreme
distress extreme

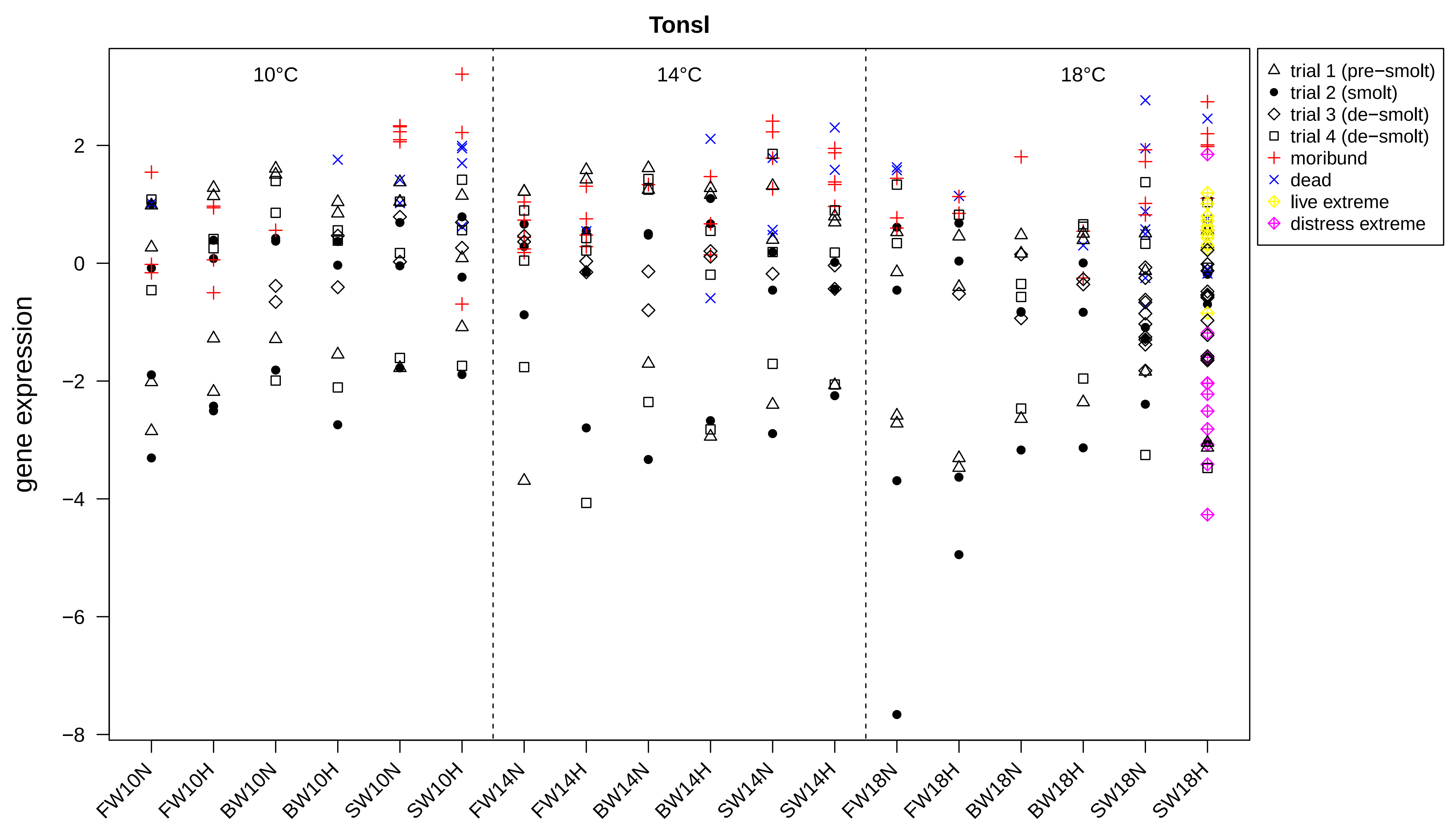

Tonsl
10°C
14°C
18°C
trial 1 (pre−smolt)
trial 2 (smolt)
trial 3 (de−smolt)
trial 4 (de−smolt)
moribund
dead
live extreme
distress extreme
gene expression
2
0
−2
−4
−6
−8
FW10N
FW10H
BW10N
BW10H
SW10N
SW10H
FW14N
FW14H
BW14N
BW14H
SW14N
SW14H
FW18N
FW18H
BW18N
BW18H
SW18N
SW18H
