## Supplemental Table 1 for "Identification of hypoxia-specific biomarkers in salmonids using RNA-sequencing and validation using high-throughput qPCR"

**Table S1.** The summary of qPCR TaqMan assay designs for 30 candidate hypoxia stress genes. Presented are the forward (F), reverse (R), and probe sequences (P).

| **Gene name** | **Assay name** | **Primers and TaqMan Probes** |
| --- | --- | --- |
| Anillin-like | Anillin | F- TGGTGAATGGATTTGGAGCTT  R- GGGTGGTTCCAATAGGACATGT  P- CGCTTCTTTGTCCTGGAG |
| Ribonucleoside-diphosphate reductase subunit M2-like | RRM2 | F- TGCTGCTAGTGATGGCATTGT  R- TTTGGAAACCATAGAAGCATCTTG  P- ATTTACACAGGAAGTCCAGG |
| Aurora kinase B-like | AURKB | F- GAAATGTGGTCGCTTCGATGA  R- CATCAGCCAACTCCTCCATGT  P- CAGCGCACTGCTAC |
| Ribonucleoside-diphosphate reductase large subunit-like | RRM1 | F- GCTGGAAGCAGGGTCTGAAG  R- GTTGGCTGCAGGCTTGGT  P- CGGGCATGTACTACCT |
| Kinesin-like protein KIF2C | KIF2C | F- CGGCCAAACTGGAAGTGGTA  R- TTCTGGCTCTTCCCTGAAAAGT  P- AACTCACACAATGGGAG |
| Non-SMC condensin II complex subunit D3 | Ncapd3 | F- TCACAGGTCCAGAAGAAAGTGTTC  R- GCGAGCGCTTCTGTTCCA  P- TCGAGAGCATGATTCC |
| Histone H2A.V | H2AFV | F- CCCTCATCAAGGCCACCAT  R- CGATCAGGGACTTGTGGATGT  P- AGGCGGTGTGATCC |
| Kinesin family member 15-like | Kif15 | F- CAGGCAGGTCTTCTCCAAGCT  R- AGTTTGGATGATAGCCTCCTTCTG  P- CACAGGATCAGACTGC |
| Rho GTPase-activating protein 11A-like | ARHGAP11A | F- TGATAATTACTCCCAACTCCAAGAGA  R- CTTGCTGGAGCCATAACTTTGA  P- AACTTCCTTTGGAATCTG |
| Rab9 effector protein with kelch motifs-like | Rab9 | F- CAGGCCCTCAATGACACCTT  R- GGAGAGCTGTGGGTGAACCA  P- CCTTCAACACAGACACC |
| ERCC excision repair 6 like, spindle assembly checkpoint helicase | ERCC6L | F- TTGTATGGTCTCCACAGAGATGGT  R- GTCTTCCCTAAGCCCATGTCAT  P- TCAAGGAGGAATCCTAG |
| Tonsoku-like protein | Tonsl | F- CCAATGCTTGCACCAATCAG  R- TCTGCCTCGCTATGTGGTACTG  P- TGCGGGTCAGGGTT |
| Citron Rho-interacting kinase-like | CIT | F- GATCTCTAGGTTTCAGCGCAAGA  R- TGAGCTCCACATCCTTTTGGT  P- ACCTGGAGTCAGTTCT |
| Proliferating cell nuclear antigen | PCNA | F- TTCTACTTGAACCAAACACAGATTCC  R- TTTCTTCAGGATGGATCCCTGTA  P- TGTTTGAGGCTCGCTTG |
| Claspin-like | Claspin | F- ATGCGGGCTGCCCTATC  R- CTCTTGAAGAACTGGTCGATGCT  P- CATGCCTGAGCCCAA |
| NDC80, kinetochore complex component | Ndc80 | F- CCGGCTCCAACACATCTTGT  R- CTTCTCCCAGTTGATCCTCTCAA  P- CACCCAGAAGTTCACTC |
| Lamin B1 | Lmnb1 | F- CGAGCTCGCTGACGCTAGA  R- CAACCTCGCGCGTTCTTT  P- ATCTCTCGATGACACCGC |
| Condensin-2 complex subunit G2-like | NCAPG2 | F- CTGTGGCACTCGGCTGTTC  R- GAGTCGAAGCCCGACCAA  P- CGCTCTAGGAAACTT |
| Chromosome-associated kinesin KIF4-like | Kif4 | F- GAATCTGCAGCACGTCATCCT  R- CAGCCAACGCCTCAATGG  P- CAGCTCCAGGATGAGAGT |
| Structural maintenance of chromosomes protein 4-like | SMC4 | F- AGCGTGTGGCTGAGTTGGAT  R- GATGTTGAAGCCAGCCATGA  P- ATCACCACAGAGAGAGACA |
| Malignant fibrous histiocytoma-amplified sequence 1 homolog | MFHAS1 | F- CCGAGGCCTGGGTGAAC  R- TCAGCTGCTCCACAGAGAAGAA  P- TCAGTGGCTGCTAGTC |
| Receptor activity-modifying protein 1-like | RAMP1 | F- CGAACCAAGTGGTGCAAGACT  R- CCGGACATGCCTGGAAGA  P- CTTCATCCAGATCCATTC |
| Peptidylglycine alpha-amidating monooxygenase_106571984 | PAM | F- TGTAGGAGATGCAGCCACAGA  R- TCTTAACAGAGCGATGTTCAGCTT  P- TGCTAAAGTTCTCCTCTGAC |
| protein family with sequence similarity 214, A | FAM214A | F- GATGAGGAGTGTGTCTGTCAGTTCTC  R- GAGCCTCAGGGCAACACTCT  P- CCAGCGGAGCACCT |
| Calphotin | Calphotin | F- CCTTCAAGCGTGTTGACATCA  R- CAGTGTTCTCCCTGTAGCACTTCA  P- TGTGTGGTGACCTGCA |
| Gutathione peroxidase 3 | GPX3 | F- AGGCCAGTCCTTCAGTGCAT  R- GGCAGGACCAGGAGGTAACA  P- TGGGCCTGGTAACC |
| Endothelin-converting enzyme 2 | ECE-2 | F- AGCAGACGCTGGGAGAGAAC  R- CACGCCTGGTACGCCTTATAG  P- CTGACAACGGAGGCC |
| Cyclin dependent kinase inhibitor 1B | CDKN1B | F- CGTCCTCAGCGAAATGGAA  R- CCATTCGAATCTCCCGTTTAAT  P- TCGATTTTTCAAGTCAAAC |
| Transcription factor SOX-5 | SOX-5 | F- GAGAGTACAAGGCCATTATGAGGAA  R- GCCCGCCGAGGACAAG  P- CGGCAGGAGATGAG |
| Acyl-coenzyme A thioesterase 9, mitochondrial | ACOT9 | F- CAAGATGCACATGTCTCAGTACCA  R- AGGGTCTCTGGCCACCATAA  P- CCAGTTCTGGATGCTAC |
