## Supplemental Table 2 for "Identification of hypoxia-specific biomarkers in salmonids using RNA-sequencing and validation using high-throughput qPCR"

**Table S2.** Correlations between 240 candidate gill hypoxia genes and their first two principal components. A principal component analysis used the scaled and normalized counts (cpm) of 240 genes. Displayed are Pearson correlations (*r*) and their significance (*p*) between the gene counts and the first two principal components (PC1 and PC2). Positive and negative correlations are upregulated and downregulated in hypoxia than normoxia, respectively. Results are sorted by *p*-value for PC1.

| Gene name | Gene description | *r*1 | *p*1 | *r*2 | *p*2 |
| --- | --- | --- | --- | --- | --- |
| LOC112218460 | anillin-like | -0.95189 | 8.81E-13 | 0.097139 | 0.651594 |
| LOC112249990 | ribonucleoside-diphosphate reductase subunit M2-like | -0.92312 | 1.33E-10 | 0.162189 | 0.448943 |
| LOC112218878 | aurora kinase B-like | -0.9088 | 8.17E-10 | 0.167254 | 0.434708 |
| LOC112228646 | ribonucleoside-diphosphate reductase large subunit-like | -0.89364 | 4.12E-09 | 0.259303 | 0.221116 |
| LOC112226944 | kinesin-like protein KIF2C | -0.89218 | 4.76E-09 | 0.23932 | 0.26004 |
| top2a | DNA topoisomerase II alpha | -0.89049 | 5.60E-09 | 0.400934 | 0.052178 |
| ncapd3 | non-SMC condensin II complex subunit D3 | -0.88624 | 8.34E-09 | 0.265898 | 0.209172 |
| LOC112262552 | histone H2A.V | -0.88545 | 8.96E-09 | 0.23774 | 0.263293 |
| kif15 | kinesin family member 15 | -0.88468 | 9.61E-09 | 0.242711 | 0.253141 |
| LOC112256655 | rho GTPase-activating protein 11A-like | -0.8838 | 1.04E-08 | 0.113267 | 0.598212 |
| ncapd2 | non-SMC condensin I complex subunit D2 | -0.88041 | 1.40E-08 | 0.392884 | 0.057543 |
| LOC112259694 | rab9 effector protein with kelch motifs-like | -0.88036 | 1.41E-08 | 0.008227 | 0.969567 |
| ercc6l | ERCC excision repair 6 like, spindle assembly checkpoint helicase | -0.8803 | 1.42E-08 | 0.258103 | 0.223336 |
| LOC112256480 | tonsoku-like protein | -0.87516 | 2.20E-08 | 0.204184 | 0.338553 |
| LOC112251434 | lamin-B2-like | -0.87474 | 2.27E-08 | 0.368745 | 0.076207 |
| LOC112215505 | citron Rho-interacting kinase-like | -0.8736 | 2.50E-08 | 0.3016 | 0.152069 |
| LOC112248619 | proliferating cell nuclear antigen | -0.87069 | 3.16E-08 | 0.123867 | 0.564172 |
| LOC112229914 | claspin-like | -0.8681 | 3.89E-08 | 0.176494 | 0.409377 |
| ndc80 | NDC80, kinetochore complex component | -0.86564 | 4.71E-08 | 0.186882 | 0.381903 |
| LOC112249909 | mitotic checkpoint serine/threonine-protein kinase BUB1 beta-like | -0.86265 | 5.91E-08 | 0.364023 | 0.080341 |
| lmnb1 | lamin B1 | -0.86212 | 6.15E-08 | 0.301504 | 0.152207 |
| LOC112228013 | condensin-2 complex subunit G2-like | -0.86067 | 6.85E-08 | 0.047134 | 0.82688 |
| LOC112231350 | chromosome-associated kinesin KIF4-like | -0.85421 | 1.09E-07 | 0.423934 | 0.038971 |
| LOC112265693 | structural maintenance of chromosomes protein 4-like | -0.8533 | 1.17E-07 | 0.448816 | 0.027813 |
| actl6a | actin like 6A | -0.84991 | 1.47E-07 | 0.085557 | 0.691 |
| LOC112253110 | kinesin-like protein KIF11-B | -0.84744 | 1.74E-07 | 0.448441 | 0.02796 |
| LOC112253851 | lymphocyte-specific helicase-like | -0.84537 | 2.00E-07 | 0.071013 | 0.741599 |
| rrm2 | ribonucleotide reductase regulatory subunit M2 | -0.84442 | 2.13E-07 | 0.189171 | 0.375994 |
| ubr7 | ubiquitin protein ligase E3 component n-recognin 7 (putative) | -0.84395 | 2.20E-07 | 0.123906 | 0.564047 |
| tyms | thymidylate synthetase | -0.84253 | 2.41E-07 | -0.09206 | 0.668785 |
| LOC112264801 | transcription factor 19-like | -0.84228 | 2.45E-07 | -0.05558 | 0.796434 |
| gtse1 | G2 and S-phase expressed 1 | -0.84024 | 2.79E-07 | 0.415735 | 0.043335 |
| espl1 | extra spindle pole bodies like 1, separase | -0.83789 | 3.24E-07 | 0.477644 | 0.018251 |
| LOC112224457 | disks large-associated protein 5-like | -0.83749 | 3.32E-07 | 0.45812 | 0.024368 |
| LOC112219045 | inner centromere protein A-like | -0.83713 | 3.40E-07 | 0.467587 | 0.021223 |
| LOC112230962 | G2/mitotic-specific cyclin-B1-like | -0.83642 | 3.55E-07 | 0.360379 | 0.083645 |
| LOC112249393 | kinesin-like protein KIF23 | -0.8302 | 5.19E-07 | 0.449583 | 0.027515 |
| LOC112220852 | DNA replication licensing factor mcm7-like | -0.8283 | 5.82E-07 | -0.04832 | 0.822594 |
| LOC112251353 | nuclear autoantigenic sperm protein-like | -0.82811 | 5.88E-07 | 0.193195 | 0.365732 |
| LOC112260003 | histone H2A.Z | -0.82671 | 6.39E-07 | 0.379012 | 0.067777 |
| LOC112228262 | deoxycytidine kinase-like | -0.82584 | 6.72E-07 | 0.295568 | 0.160843 |
| LOC112250026 | mitotic checkpoint serine/threonine-protein kinase BUB1-like | -0.82543 | 6.88E-07 | 0.446334 | 0.028795 |
| nuf2 | NUF2, NDC80 kinetochore complex component | -0.8232 | 7.82E-07 | 0.206435 | 0.333139 |
| LOC112262953 | centromere-associated protein E-like | -0.82318 | 7.83E-07 | 0.454999 | 0.025483 |
| LOC112239878 | uncharacterized LOC112239878 | -0.82244 | 8.17E-07 | 0.121091 | 0.573004 |
| sass6 | SAS-6 centriolar assembly protein | -0.82093 | 8.90E-07 | -0.05939 | 0.782826 |
| LOC112238559 | uncharacterized LOC112238559 | -0.81444 | 1.28E-06 | 0.488692 | 0.015386 |
| neil3 | nei like DNA glycosylase 3 | -0.81113 | 1.52E-06 | 0.056919 | 0.791647 |
| LOC112219506 | malignant fibrous histiocytoma-amplified sequence 1 homolog | 0.810857 | 1.55E-06 | 0.060474 | 0.778937 |
| kifc1 | kinesin family member C1 | -0.80906 | 1.70E-06 | 0.483234 | 0.016752 |
| LOC112245345 | PCNA-associated factor-like | -0.8073 | 1.86E-06 | 0.336041 | 0.108392 |
| LOC112229667 | RNA polymerase II subunit A C-terminal domain phosphatase-like | -0.80719 | 1.88E-06 | -0.09107 | 0.672133 |
| ccnf | cyclin F | -0.80538 | 2.06E-06 | 0.515212 | 0.009981 |
| LOC112256353 | cyclin-A2-like | -0.80395 | 2.22E-06 | 0.267586 | 0.206185 |
| iqgap3 | IQ motif containing GTPase activating protein 3 | -0.80357 | 2.26E-06 | 0.461015 | 0.023369 |
| LOC112229614 | anillin-like | -0.80237 | 2.40E-06 | 0.484186 | 0.016507 |
| LOC112250348 | dnaJ homolog subfamily C member 5-like | -0.8023 | 2.41E-06 | 0.302088 | 0.151375 |
| LOC112233066 | targeting protein for Xklp2-like | -0.79965 | 2.75E-06 | 0.460572 | 0.023519 |
| LOC112224264 | kinesin-like protein KIF20B | -0.79701 | 3.14E-06 | 0.382112 | 0.065378 |
| LOC112223799 | receptor activity-modifying protein 1-like | 0.79092 | 4.21E-06 | -0.17556 | 0.411914 |
| NA | E2F transcription factor 3 | -0.78962 | 4.48E-06 | -0.15758 | 0.462092 |
| kif22 | kinesin family member 22 | -0.789 | 4.62E-06 | 0.402836 | 0.050969 |
| LOC112215881 | targeting protein for Xklp2-like | -0.78678 | 5.12E-06 | 0.441463 | 0.030803 |
| LOC112264594 | anillin-like | -0.78632 | 5.23E-06 | 0.48533 | 0.016216 |
| cenpw | centromere protein W | -0.78442 | 5.71E-06 | 0.05837 | 0.786454 |
| LOC112237254 | denticleless protein homolog | -0.78403 | 5.82E-06 | -0.02643 | 0.90244 |
| pole2 | DNA polymerase epsilon 2, accessory subunit | -0.78191 | 6.41E-06 | -0.16531 | 0.440153 |
| LOC112253106 | rhotekin-2-like | -0.78139 | 6.56E-06 | 0.401285 | 0.051954 |
| cep135 | centrosomal protein 135 | -0.78072 | 6.76E-06 | 0.353064 | 0.090585 |
| LOC112228882 | protein DEK-like | -0.78045 | 6.84E-06 | 0.266359 | 0.208353 |
| LOC112250900 | chromatin assembly factor 1 subunit A-like | -0.78001 | 6.98E-06 | -0.21164 | 0.320825 |
| NA | zinc finger protein 367-like | -0.77912 | 7.26E-06 | 0.512671 | 0.010419 |
| pam | peptidylglycine alpha-amidating monooxygenase | 0.778638 | 7.42E-06 | 0.049264 | 0.819183 |
| LOC112221868 | aurora kinase C-like | -0.77809 | 7.60E-06 | 0.492525 | 0.014483 |
| NA | FAM214A-like | 0.775327 | 8.59E-06 | -0.02241 | 0.917231 |
| LOC112228990 | histone H2AX | -0.77484 | 8.78E-06 | -0.00562 | 0.979211 |
| racgap1 | Rac GTPase activating protein 1 | -0.77376 | 9.20E-06 | 0.542533 | 0.006161 |
| LOC112230732 | DNA polymerase theta-like | -0.77227 | 9.82E-06 | -0.09497 | 0.658914 |
| LOC112217492 | mitotic checkpoint serine/threonine-protein kinase BUB1-like | -0.77218 | 9.85E-06 | 0.568198 | 0.003772 |
| wdhd1 | WD repeat and HMG-box DNA binding protein 1 | -0.77202 | 9.92E-06 | -0.18485 | 0.387201 |
| LOC112218911 | calphotin-like | 0.767805 | 1.19E-05 | -0.05576 | 0.795793 |
| LOC112223116 | cell division cycle-associated protein 3-like | -0.76487 | 1.34E-05 | 0.453099 | 0.026182 |
| LOC112236066 | glutathione peroxidase 3-like | 0.764464 | 1.37E-05 | -0.02272 | 0.916063 |
| LOC112245091 | dolichyl-diphosphooligosaccharide--protein glycosyltransferase subunit STT3A | -0.76431 | 1.38E-05 | 0.118635 | 0.580867 |
| LOC112232992 | dual specificity protein kinase Ttk-like | -0.76341 | 1.43E-05 | 0.213132 | 0.31734 |
| thumpd1 | THUMP domain containing 1 | -0.76011 | 1.63E-05 | -0.10414 | 0.628187 |
| LOC112231188 | kinesin-like protein KIF20A | -0.75676 | 1.87E-05 | 0.554135 | 0.004959 |
| rtel1 | regulator of telomere elongation helicase 1 | -0.75352 | 2.13E-05 | -0.3122 | 0.137487 |
| LOC112225439 | chromatin assembly factor 1 subunit B-like | -0.75314 | 2.16E-05 | -0.34447 | 0.099275 |
| LOC112255192 | cyclin-dependent kinase 1 | -0.75261 | 2.21E-05 | 0.546964 | 0.005676 |
| LOC112216690 | endothelin-converting enzyme 2-like | 0.750775 | 2.37E-05 | 0.253743 | 0.23153 |
| eme1 | essential meiotic structure-specific endonuclease 1 | -0.74843 | 2.60E-05 | 0.328824 | 0.116672 |
| LOC112226524 | G-protein-signaling modulator 2-like | -0.74643 | 2.81E-05 | 0.26602 | 0.208956 |
| LOC112214837 | abnormal spindle-like microcephaly-associated protein homolog | -0.74507 | 2.96E-05 | 0.528526 | 0.007929 |
| cdkn1b | cyclin dependent kinase inhibitor 1B | 0.744469 | 3.02E-05 | 0.191772 | 0.369341 |
| LOC112217828 | serine/arginine-rich splicing factor 3 | -0.74175 | 3.35E-05 | -0.0636 | 0.767821 |
| NA | UNKNOWN | -0.74136 | 3.40E-05 | 0.383668 | 0.064199 |
| LOC112228552 | ras-related and estrogen-regulated growth inhibitor-like protein | 0.740487 | 3.51E-05 | 0.403917 | 0.050292 |
| cenpo | centromere protein O | -0.73815 | 3.83E-05 | -0.29485 | 0.161917 |
| LOC112220417 | suppressor APC domain-containing protein 2-like | -0.73592 | 4.16E-05 | -0.22603 | 0.288222 |
| stil | STIL, centriolar assembly protein | -0.73384 | 4.48E-05 | 0.256252 | 0.226791 |
| LOC112221983 | transcription factor E2F1-like | -0.73328 | 4.57E-05 | 0.040135 | 0.852291 |
| LOC112215028 | transcription factor SOX-5-like | 0.73291 | 4.63E-05 | 0.216739 | 0.309024 |
| LOC112241904 | serine/threonine-protein kinase PLK1-like | -0.73168 | 4.84E-05 | 0.529272 | 0.007825 |
| NA | UNKNOWN | 0.729766 | 5.19E-05 | 0.059905 | 0.78097 |
| LOC112264390 | shugoshin 1-like | -0.72835 | 5.45E-05 | 0.582269 | 0.002833 |
| gins3 | GINS complex subunit 3 | -0.72581 | 5.96E-05 | -0.12713 | 0.55387 |
| LOC112231339 | peptidyl-prolyl cis-trans isomerase FKBP2-like | -0.72281 | 6.62E-05 | -0.25111 | 0.236571 |
| LOC112223647 | angiotensin-converting enzyme-like | 0.720183 | 7.24E-05 | 0.099537 | 0.643543 |
| LOC112249961 | ectonucleotide pyrophosphatase/phosphodiesterase family member 1-like | 0.715024 | 8.62E-05 | -0.16295 | 0.446778 |
| LOC112262569 | UPF0454 protein C12orf49 homolog | 0.714297 | 8.83E-05 | 0.255886 | 0.227479 |
| NA | UNKNOWN | -0.71355 | 9.05E-05 | -0.17128 | 0.423564 |
| dna2 | DNA replication helicase/nuclease 2 | -0.71169 | 9.63E-05 | 0.224184 | 0.292286 |
| LOC112225466 | centromere protein J-like | -0.7091 | 0.000105 | 0.035955 | 0.867533 |
| LOC112228816 | barrier-to-autointegration factor | -0.70864 | 0.000106 | -0.15964 | 0.456202 |
| LOC112253028 | cGMP-dependent protein kinase 1-like | 0.70798 | 0.000109 | -0.1288 | 0.54864 |
| LOC112227719 | myb-related protein A-like | -0.70708 | 0.000112 | -0.28867 | 0.171297 |
| LOC112236577 | long-chain-fatty-acid--CoA ligase 4-like | 0.705743 | 0.000117 | -0.15747 | 0.462419 |
| LOC112248982 | DNA damage-binding protein 1-like | -0.70312 | 0.000127 | -0.25681 | 0.225749 |
| LOC112235673 | EEF1A lysine methyltransferase 4-like | -0.69907 | 0.000144 | -0.24745 | 0.243706 |
| LOC112232029 | high mobility group protein B2-like | -0.69884 | 0.000145 | -0.31407 | 0.135023 |
| pold1 | DNA polymerase delta 1, catalytic subunit | -0.69265 | 0.000176 | -0.31135 | 0.13862 |
| LOC112249379 | nicotinamide riboside kinase 2-like | 0.689692 | 0.000193 | 0.227891 | 0.284167 |
| LOC112215182 | semaphorin-3ab-like | 0.689611 | 0.000193 | 0.419927 | 0.041059 |
| LOC112242225 | uncharacterized protein C17orf62 homolog | -0.6877 | 0.000204 | -0.2815 | 0.182671 |
| LOC112251159 | regulator of G-protein signaling 4-like | 0.684114 | 0.000227 | 0.446865 | 0.028583 |
| dync1h1 | dynein cytoplasmic 1 heavy chain 1 | -0.67928 | 0.000262 | -0.35053 | 0.093086 |
| LOC112263008 | apelin receptor A | 0.679026 | 0.000264 | 0.376473 | 0.069793 |
| LOC112263729 | transcriptional repressor CTCF-like | -0.67812 | 0.000271 | 0.559862 | 0.004442 |
| LOC112232096 | synembryn-B-like | -0.67809 | 0.000271 | 0.248782 | 0.241094 |
| NA | UNKNOWN | -0.67442 | 0.000301 | -0.223 | 0.294917 |
| LOC112236709 | E3 ubiquitin-protein ligase TRIM71-like | -0.67409 | 0.000304 | -0.0196 | 0.927555 |
| LOC112249143 | ADM-like | 0.672756 | 0.000316 | 0.295191 | 0.161402 |
| LOC112247189 | hypoxia-inducible factor 1-alpha inhibitor | 0.668775 | 0.000353 | 0.482602 | 0.016916 |
| LOC112217507 | midasin-like | -0.66769 | 0.000364 | -0.16687 | 0.435784 |
| mcm8 | minichromosome maintenance 8 homologous recombination repair factor | -0.66736 | 0.000367 | -0.26904 | 0.203645 |
| NA | UNKNOWN | 0.66693 | 0.000372 | 0.474682 | 0.019089 |
| LOC112221551 | semaphorin-3ab-like | 0.660693 | 0.000441 | 0.184609 | 0.387823 |
| NA | UNKNOWN | 0.65839 | 0.000469 | 0.10629 | 0.621081 |
| LOC112260474 | Krueppel-like factor 4 | 0.65046 | 0.000579 | 0.222464 | 0.296101 |
| LOC112262024 | 14-3-3 protein zeta-like | -0.64663 | 0.000639 | -0.26286 | 0.214623 |
| LOC112252887 | nuclear receptor coactivator 4-like | 0.645517 | 0.000658 | -0.13894 | 0.517326 |
| LOC112221822 | lysine-specific demethylase 9-like | 0.641114 | 0.000736 | 0.552184 | 0.005146 |
| LOC112223699 | histone H1.0-B-like | 0.641012 | 0.000738 | 0.339695 | 0.104369 |
| NA | DENN domain-containing protein 5B-like | 0.638957 | 0.000777 | 0.121336 | 0.572222 |
| LOC112227748 | R-spondin-3-like | 0.638326 | 0.000789 | 0.101093 | 0.63834 |
| LOC112253174 | hypoxia-inducible factor 1-alpha inhibitor | 0.635982 | 0.000837 | 0.450921 | 0.027001 |
| NA | plexin-A1-like | 0.634565 | 0.000867 | -0.1139 | 0.596167 |
| LOC112217550 | connective tissue growth factor-like | 0.634026 | 0.000878 | 0.054465 | 0.800448 |
| NA | UNKNOWN | 0.633599 | 0.000888 | -0.16494 | 0.441181 |
| LOC112261313 | 15-hydroxyprostaglandin dehydrogenase [NAD(+)]-like | 0.633544 | 0.000889 | -0.31667 | 0.131641 |
| NA | intersectin 1 | -0.62847 | 0.001006 | 0.211586 | 0.320946 |
| ankrd37 | ankyrin repeat domain 37 | 0.628398 | 0.001008 | 0.363031 | 0.08123 |
| NA | PI-PLC X domain-containing protein 1-lik | -0.62603 | 0.001067 | -0.30754 | 0.143762 |
| LOC112237955 | E3 ubiquitin-protein ligase TRIM7-like | 0.625981 | 0.001068 | 0.317822 | 0.130168 |
| NA | meteorin-like protein | 0.625568 | 0.001079 | 0.390335 | 0.059328 |
| LOC112225560 | multidrug resistance protein 1B-like | 0.621334 | 0.001193 | 0.075296 | 0.726581 |
| LOC112263666 | ADM-like | 0.619626 | 0.001242 | -0.23664 | 0.265573 |
| LOC112225033 | rho GTPase-activating protein 20-like | 0.610558 | 0.001531 | 0.474172 | 0.019236 |
| LOC112245149 | uncharacterized LOC112245149 | 0.608741 | 0.001596 | 0.009464 | 0.964991 |
| LOC112252728 | activating molecule in BECN1-regulated autophagy protein 1-like | -0.60802 | 0.001622 | 0.152281 | 0.477482 |
| slc18a1 | solute carrier family 18 member A1 | 0.607421 | 0.001644 | 0.137195 | 0.522646 |
| plcd1 | phospholipase C delta 1 | 0.605253 | 0.001726 | -0.28573 | 0.175898 |
| LOC112224548 | interferon gamma receptor 1-like | 0.604946 | 0.001738 | 0.470388 | 0.020359 |
