## Supplemental Table 3 for "Identification of hypoxia-specific biomarkers in salmonids using RNA-sequencing and validation using high-throughput qPCR"

**Table S3.** The results of correlation analysis of 29 discovered genes between the log_2_ fold change from the RNA-seq data and the log_2_ fold change of the qRT-PCR platform for the same 24 samples used for both platforms. The significant correlated genes are highlighted p<0.05.

| **Genes** | **r** | **P value** |
| --- | --- | --- |
| ACOT9 | -0.0523 | 0.8081 |
| Anillin | 0.0867 | 0.6872 |
| ARHGAP11A | 0.5970 | 0.0021 |
| AURKB | 0.6987 | 0.0001 |
| calphotin | 0.3230 | 0.1237 |
| CDKN1B | -0.0207 | 0.9235 |
| CIT | 0.7014 | 0.0001 |
| claspin | 0.6611 | 0.0004 |
| ECE.2 | 0.3970 | 0.0547 |
| ERCC6L | 0.8735 | 0.0000 |
| FAM214A | 0.3222 | 0.1247 |
| GPX3 | 0.2172 | 0.3080 |
| H2AFV | 0.3792 | 0.0677 |
| kif15 | 0.7882 | 0.0000 |
| KIF2C | 0.7459 | 0.0000 |
| kif4 | 0.7808 | 0.0000 |
| lmnb1 | 0.1685 | 0.4313 |
| MFHAS1 | -0.0612 | 0.7762 |
| ncapd3 | 0.6504 | 0.0006 |
| NCAPG2 | 0.3365 | 0.1079 |
| ndc80 | 0.3812 | 0.0660 |
| PAM | 0.1595 | 0.4565 |
| PCNA | 0.5479 | 0.0056 |
| rab9 | 0.5700 | 0.0036 |
| RAMP1 | 0.4223 | 0.0398 |
| RRM1 | 0.5576 | 0.0046 |
| RRM2 | 0.7520 | 0.0000 |
| SOX.5 | 0.0249 | 0.9079 |
| Tonsl | 0.6624 | 0.0004 |
