## Supplemental Fig. caption for "Identification of hypoxia-specific biomarkers in salmonids using RNA-sequencing and validation using high-throughput qPCR"

**Supplemental Figures caption**

**Figure S1.** Individual-based expression for all 29 discovered genes after six days of treatment. Salinity treatment symbols are SW, BW, and FW for seawater, brackish, and freshwater groups. Temperature treatment symbols are 10, 14, and 18 °C groups. Dissolved oxygen treatment symbols are N for normoxia and H for hypoxia groups.

**Figure S2.** Canonical plots of the first two principal components using 4, 11 and 6 hypoxia biomarkers returned by the Shrunken Centroid method for pre-smolt fish in panels (a) freshwater (FW), (b) brackish water (BW), and (c) seawater (SW), respectively. Temperature treatment symbols are 10, 14, and 18 °C groups. Principal component analysis was performed on the training set (left panel) and then applied to the testing set (right panel). Centroids are represented by the largest point of the same colour. Arrows represent loading vectors of the biomarkers using the training set.

**Figure S3.** Canonical plots of the first two principal components using 9, 4 and 5 hypoxia biomarkers returned by the Shrunken Centroid method for smolt fish in panels (a) freshwater (FW), (b) brackish water (BW), and (c) seawater (SW), respectively. Temperature treatment symbols are 10, 14, and 18 °C groups. Principal component analysis was performed on the training set (left panel) and then applied to the testing set (right panel). Centroids are represented by the largest point of the same colour. Arrows represent loading vectors of the biomarkers using the training set.

**Figure S4.** Canonical plots of the first two principal components using 6, 3 and 15 hypoxia biomarkers returned by the Shrunken Centroid method for de-smolt fish in panels (a) freshwater (FW), (b) brackish water (BW), and (c) seawater (SW), respectively. Temperature treatment symbols are 10, 14, and 18 °C groups. Principal component analysis was performed on the training set (left panel) and then applied to the testing set (right panel). Centroids are represented by the largest point of the same colour. Arrows represent loading vectors of the biomarkers using the training set.

**Figure S5.** Canonical plots of the first two principal components using 4 and 14 hypoxia biomarkers returned by the Shrunken Centroid method for distressed (moribund/recently dead) fish in panels (a) freshwater (FW) and (b) seawater (SW), respectively. Temperature treatment symbols are 10, 14, and 18 °C groups. Principal component analysis was performed on the training set (left panel) and then applied to the testing set (right panel). Centroids are represented by the largest point of the same colour. Arrows represent loading vectors of the biomarkers using the training set.
